# Two different epigenetic pathways detected in wild three-spined sticklebacks are involved in salinity adaptation

**DOI:** 10.1101/649574

**Authors:** Melanie J. Heckwolf, Britta S. Meyer, Robert Häsler, Marc P. Höppner, Christophe Eizaguirre, Thorsten B. H. Reusch

## Abstract

While environmentally inducible epigenetic marks are discussed as one mechanism of transgenerational plasticity, environmentally stable epigenetic marks emerge randomly. When resulting in variable phenotypes, stable marks can be targets of natural selection analogous to DNA sequence-based adaptation processes. We studied both postulated pathways in natural populations of three-spined sticklebacks (Gasterosteus aculeatus) and sequenced their methylomes and genomes across a salinity cline. Consistent with local adaptation, populations showed differential methylation (pop-DMS) at genes enriched for osmoregulatory processes. In a two-generation experiment, 62% of these pop-DMS were insensitive to salinity manipulation, suggesting that they could be stable targets for natural selection. Two-thirds of the remaining inducible pop-DMS became more similar to patterns detected in wild populations from the corresponding salinity, and this pattern accentuated over consecutive generations, indicating a mechanism of adaptive transgenerational plasticity. Natural DNA methylation patterns can thus be attributed to two epigenetic pathways underlying the rapid emergence of adaptive phenotypes in the face of environmental change.

## Introduction

Recent advances in epigenetics have started to challenge our understanding of genetic diversity, inheritance and adaptive evolution (1–3). It has been suggested that epigenetic modification, for example via DNA methylation, histone modification or small RNAs, provides an additional pathway to create phenotypic diversity and ultimately contribute to rapid evolutionary adaptation (4–6). Several theoretical models posit that the heritable proportion of these molecular modifications can contribute to adaptation via two distinct information pathways (5, 7, 8). Firstly, selection-based epigenetic marks would emerge as spontaneous epimutations that remain stable across subsequent generations, although their overall stability seems to be 2-3 orders of magnitude lower as DNA base changes (7, 9). Similar to adaptation from DNA sequence-based variation these epimutations can underlie phenotypes that are targets of natural selection and thereby carry information on past selection regimes without directly responding to the current environment (5, 8, 10). Secondly, detection-based effects describe inducible epigenetic marks at defined genomic locations which are under environmental control (7). This way, the inheritance of information on environmental cues experienced by previous generations represents one rapid and reliable mechanism underlying transgenerational plasticity which is hypothesized to buffer populations under sudden environmental change (7, 11). Distinguishing between these mechanisms is crucial, since they have very different evolutionary implications. While stable epigenetic marks are in line with key tenets of the Modern Synthesis including random variation and genetic inheritance, directional processes via inducible epigenetic marks are better captured by the Extended Evolutionary Synthesis and its assumptions on non-random phenotypic variation and inclusive inheritance (12). Given an ongoing and sometimes controversial discussion of whether or not there is a need of an extended evolutionary synthesis (2), providing empirical evidence of both pathways and assessing their proportional contribution to epigenetic inheritance in nature is paramount. While the significance and principal difference of both transmission pathways is clear (4, 5, 7), empirical evidence on the proportional existence and adaptive value of these pathways is lacking. One prime objective of this study was thus to assess whether these two epigenetic pathways can be detected in nature and to test whether short term acclimation responses match patterns of DNA methylation variation of natural populations. If transgenerational experiments result in DNA methylation profiles closer to those of locally adapted natural populations, this would provide evidence that DNA methylation is mechanistically involved in adaptive transgenerational plasticity.

Studying the adaptation to ocean salinity is particularly well suited to identify selection-based and detection-based effects because spatio-temporal patterns in ocean salinity are rather stable compared to for instance temperature. Since salinity change imposes strong physiological stress with well-defined cellular effects (13), natural salinity gradients offer unparalleled opportunities to use local patterns of epigenetic variation as background against which direction and magnitude of experimental salinity manipulations can be tested. One suitable ecosystem to follow such a space-for-time approach is the Baltic Sea, a semi-enclosed marginal sea that has been dubbed a “time machine” for many predicted perturbations associated with global change (14).

Taking advantage of the Baltic Sea salinity gradient, we sequenced the methylomes (reduced representation bisulfite sequencing, RRBS) as well as whole genomes of spined-three sticklebacks (Gasterosteus aculeatus) from three locally adapted populations (15) in- and outside the Baltic Sea (6, 20 and 33 PSU). In this paper we focus on the patterns of (epi)genomic variation, while transgenerational phenotypic effects in the same populations have been described previously (16). Some key response variables are presented here to validate adaptive transgenerational plasticity.

Baltic stickleback populations are genetically differentiated from one another (genome wide average pairwise FST = 0.028 (15)) and show patterns consistent with local adaptation to salinity regimes in controlled common garden experiments (16, 17). Furthermore, sticklebacks are known for their adaptive transgenerational plasticity in response to variation in temperature (18) and changes in DNA methylation levels at osmoregulatory genes in response to within generation salinity acclimation (19, 20). However, it remains unclear whether DNA methylation mediate adaptive transgenerational plasticity, one possible mechanism allowing adaptive phenotypes to rapidly emerge in the face of environmental change. To address this question, we complemented our field survey with a two-generation salinity acclimation experiment using the mid salinity population (20 PSU) to quantify the proportion of stable (potentially selection-based) and inducible (potentially detection-based) DNA methylation within- and across generations (Figure 1). We focused on the methylation of cytosines at cytosine-phosphate-guanine dinucleotides (CpG sites), the most common methylation motif in vertebrates (21) with partial inheritance potentially facilitating adaptation in natural populations (11).

**Fig. 1:**
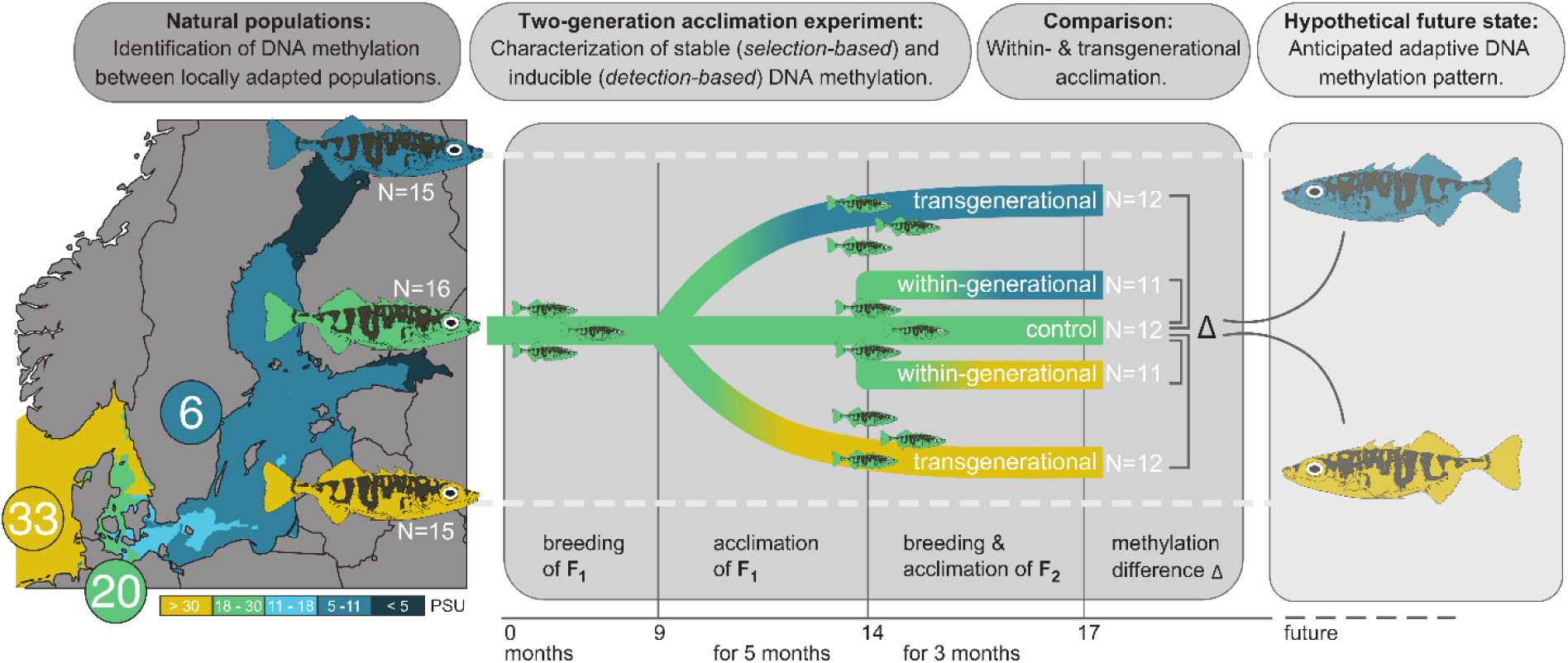
Experimental space-for-time approach. We characterized DNA methylation profiles (via Reduced Representation Bisulfite Sequencing, RRBS) and whole genomes (Whole Genome Sequencing, WGS) of fish from three populations of wild caught three-spined sticklebacks locally adapted to 6 (blue, N = 15), 20 (green, N = 16) and 33 PSU (yellow, N = 15). We also collected sticklebacks from the mid salinity location (20 PSU) and acclimated laboratory bred offspring of these fish within one (‘within-generational’) or over two (‘transgenerational’) generations to decreased (6 PSU) or increased (33 PSU) salinity, and maintained a control group at its original salinity (N = 11-12 per group, see details in Figure). Differential methylation within and across generations was assessed and compared to natural populations locally adapted to the corresponding salinity, serving as the hypothetical future DNA methylation state to capture long term adaptation processes.

We tested three inter-related hypotheses: (i) Stickleback populations originating from different salinity areas (6, 20 and 33 PSU) show differentially methylated CpG sites (hereafter pop-DMS), consistent with patterns of local adaptation. (ii) Such pop-DMS include both types of methylation sites: experimentally stable sites (potentially selection-based) and experimentally inducible sites (potentially detection-based). (iii) Upon transgenerational salinity acclimation, inducible DNA methylations become more similar to the patterns of natural populations at corresponding salinities. When associated with beneficial phenotypic effects, the latter findings would be evidence of a mechanism of adaptive transgenerational plasticity.

## Results

### Identifying differentially methylated CpG sites between stickleback populations along a natural salinity cline

Differentially methylated CpG sites between stickleback populations (hereafter pop-DMS) were determined via reduced representation bisulfite sequencing (RRBS) in 46 wild caught sticklebacks from 3 different sites that varied in average salinity (Sylt, 33 practical salinity units = PSU; Kiel, 20 PSU; Nynäshamn, 6 PSU; Figure 1). After quality and coverage filtering, we obtained 525,985 CpG sites present in all groups, corresponding to ∼4% of all CpG sites in the stickleback genome. Between pairs of wild caught populations, we detected 1,470 (comparison 20 vs. 6 PSU) and 1,158 (20 vs. 33 PSU) pop-DMS according to our significance threshold. The distribution of these sites was random with regard to the genomic features (promoter, exon, intron, and intergenic; 20 vs. 6 PSU: X23 = 3.36, P = 0.340; 20 vs. 33 PSU: X23 = 1.61, P = 0.656; Supplementary Material: Table S1) and chromosomal regions (Supplementary Figure S1A). Among these pop-DMS, 1,098 (20 vs. 6 PSU) and 871 (20 vs. 33 PSU) were located close to (< 10 kb from transcription start sites) or within genes and thereby associated with 655 and 510 genes, respectively. Many of these genes were involved in basic biological processes such as DNA repair and strand renaturation, as well as chromosome condensation and separation (Supplementary Figure S2). Particularly relevant and concordant with previous findings of local salinity adaptation (15), these genes were enriched for osmoregulatory processes such as ion transport and channel activity, renal water homeostasis and absorption, as well as urine volume regulation (Figure 2). Genes associated with ≥ 10 pop-DMS are listed in Table 1 (for all genes, see Supplementary Table S2A and S2B).

**Fig. 2:**
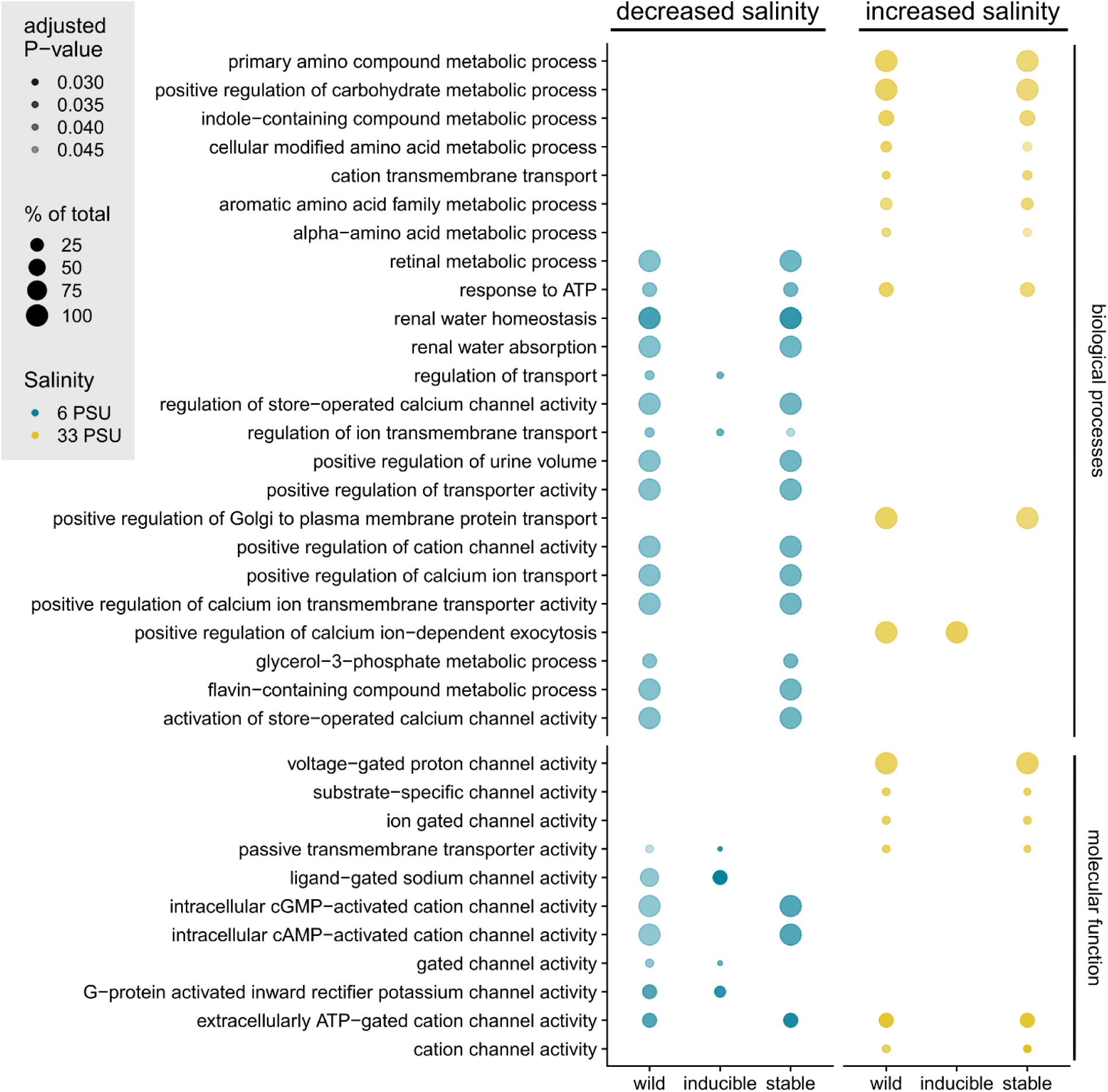
Gene ontology terms for biological processes and molecular functions. Gene ontology (GO) terms for biological processes and molecular functions under salinity increase (yellow, 20 vs. 33 PSU) and decrease (blue, 20 vs. 6 PSU) associated with differentially methylated sites between populations (pop-DMS) are presented. The graph is split into GO terms associated with pop-DMS from natural stickleback populations across a salinity cline (wild) and their experimental inducibility (inducible and stable) in a two-generation acclimation experiment. The size of the circles refers to the number of genes of this term present in our groups (in %) and the transparency to the FDR corrected P-value (darker circles refer to a lower adjusted P-value). This subset is filtered for GO terms including the following keywords: channel, transport, water, chloride, potassium, homeostasis, ion-dependent, urine, ATP, metabolic, see Figure S2 (Supplementary Material) for the full figure.

**Table 1:**
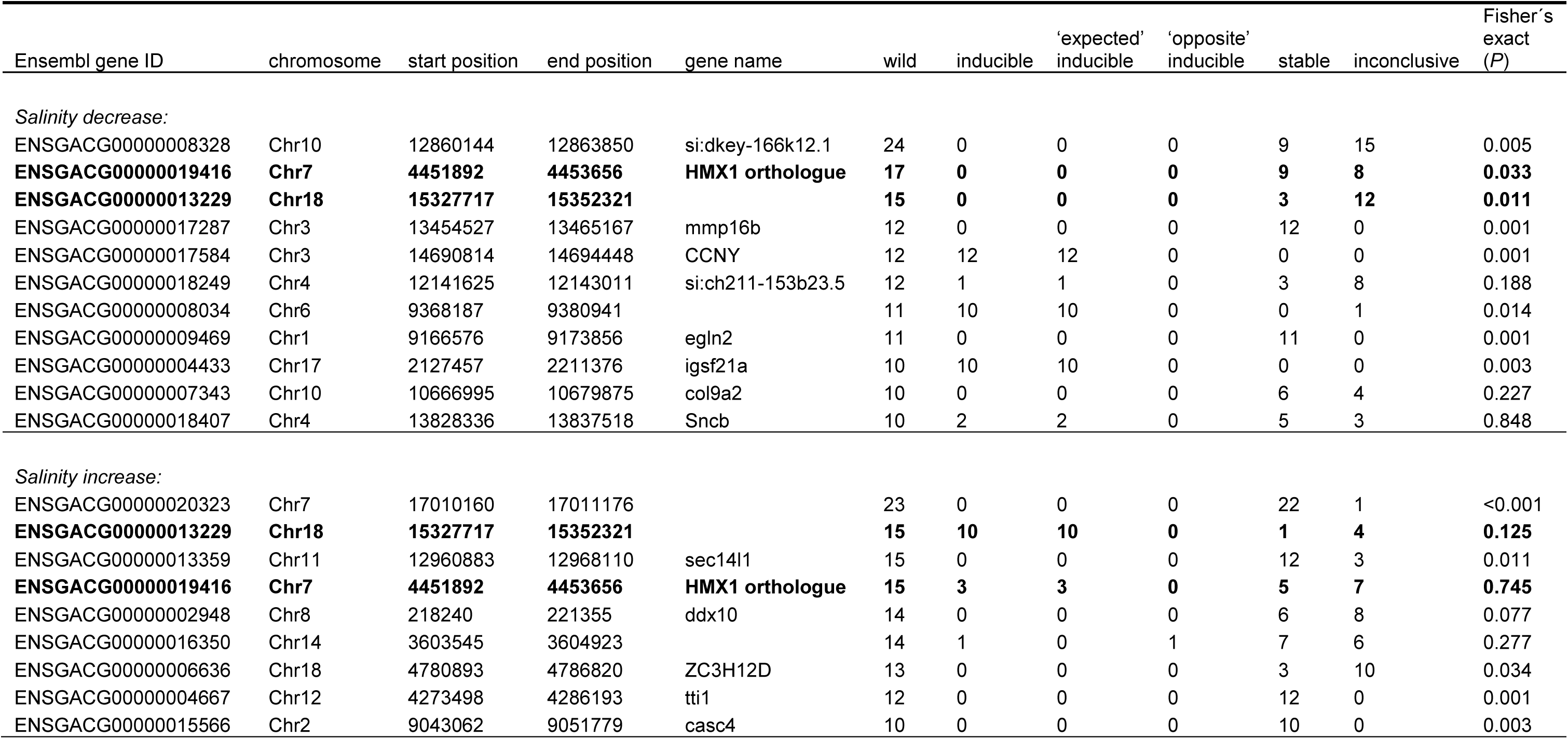
Differentially methylated genes across natural populations along a salinity cline. Genes derived from DNA methylation comparisons between natural populations associated with ≥ 10 pop-DMS (decreased salinity: KIE (20 PSU) vs. NYN (6 PSU); increased salinity: KIE (20 PSU) vs. SYL (33 PSU)). Ensembl gene ID and name as well as the position on the chromosome are listed. The numbers refer to the numbers of DMS in the population comparison (wild). These DMS were classified into ‘inducible’, ‘inconclusive’ and ‘stable’ sites according to their behavior in a two-generation salinity acclimation experiment with laboratory bred sticklebacks from the mid salinity population (20 PSU) exposed to experimental salinity increase or decrease (33 and 6 PSU respectively). Further, inducible sites were distinguished whether they matched methylation levels of the locally adapted population (‘expected’) or not (‘opposite’). Genes written in bold vary in both population comparisons. We used a Fisher’s exact test to assess whether pop-DMS associated to the same gene are correlated in their response to experimental salinity change (non-random distribution among the categories stable, inducible, inconclusive) and reported corresponding P-values. For a full table on all genes associated with 1 or more pop-DMS see Table S2A and S2B (Supplementary Material).

### Characterizing stable and inducible DNA methylation in a two-generation salinity acclimation experiment

In order to distinguish between stable and inducible DNA methylation we then conducted a two-generation salinity acclimation experiment with laboratory bred sticklebacks from the mid salinity population (Figure 1). We considered pop-DMS to be stable when both the within- and the transgenerational acclimation group were not differentially methylated compared to the control group (q-value ≥ 0.0125). On the other hand, if a pop-DMS was differentially methylated between at least one of the acclimation groups (within- and transgenerational) compared to the control group (q-value < 0.0125; methylation difference ≥ 15%) this site was considered inducible. Pop-DMS with a significant q-value not exceeding the threshold of differential DNA methylation (15%) were treated as a separate category, hereafter inconclusive. After two generations of salinity acclimation, we found that the majority of the pop-DMS remained stable, regardless of the direction of salinity change (926 pop-DMS = 63% at decreased salinity; 694 pop-DMS = 60% at increased salinity). A smaller number of pop-DMS (13%) were inducible, as they showed a significant change in CpG methylation upon experimental salinity decrease (198 pop-DMS) or increase (148 pop-DMS). An additional 24% and 27% (346 and 316 pop-DMS respectively) differed significantly between experimental treatment groups, but did not exceed the minimum threshold in differential DNA methylation of 15% employed in this study. The number of inducible pop-DMS (13%) derived from comparisons between natural populations was significantly higher compared to what would be expected from a random subset of CpG sites across the genome (< 1%; 1000 replicates; salinity decrease: X22 = 1090.7, P < 0.001; salinity increase: X22 = 967.7, P < 0.001).

### Stable and inducible methylated sites are associated with different functional gene categories

We then assessed both stable and inducible pop-DMS for associations with different putative gene function (Figure 2, Supplementary Figure S2). Genes associated with stable pop-DMS (452 and 329 under salinity decrease and increase, respectively) were enriched for basic biological processes (e.g. DNA repair and chromosome separation; Supplementary Figure S2), but also for osmoregulatory functions (e.g. ion channel activity; Figure 2). Furthermore, under increased salinity, many metabolic processes were among the stable pop-DMS (Figure 2). Inducible pop-DMS were associated with genes (100 and 82 under salinity decrease and increase, respectively) that were primarily enriched for other osmoregulatory functions regulating ion transmembrane transport (Figure 2, Supplementary Figure S2). Therefore, stable and inducible pop-DMS are not only affecting different genes but also different gene ontologies with very little overlap (Figure 2).

### Assessing the role of inducible DNA methylation in nature

Next, we investigated whether multiple pop-DMS associated with the same gene showed a correlated response to experimental salinity acclimation, which would result in a non-random distribution of pop-DMS within genes among the categories ‘stable’, ‘inducible’ and ‘inconclusive’. We found that in 13 out of 20 genes with ≥ 10 pop-DMS, these pop-DMS responded in a synchronous way across the gene upon salinity acclimation (Table 1, Fisher’s exact test, P < 0.05), suggesting that genomic sites are pre-defined and not randomly hit by epi-mutations. Secondly, we tested whether a change at inducible pop-DMS in experimental fish increased the similarity to methylation patterns in natural populations locally adapted to their salinity conditions. Of the 198 (decreased salinity) and 148 (increased salinity) inducible pop-DMS, 130 (66%) and 101 (68%), respectively, became more similar to methylation levels of the field population locally adapted to the corresponding salinity (hereafter ‘expected’ direction). Conversely, at 68 (34%, decreased salinity) and 47 (32%, increased salinity) inducible pop-DMS, experimental fish showed methylation changes in the opposite direction, meaning the similarity to methylation levels observed in the natural populations was reduced (hereafter ‘opposite’ direction).

Since genomic variation can have a strong cis-regulatory impact on epigenomic variation (22), populations may respond with similar DNA methylation changes to salinity at regions of low genetic population divergence, while at regions of high genetic population divergence DNA methylation changes might differ between populations. Thus, we hypothesized that opposite inducible pop-DMS are associated with higher genomic (DNA sequence-based) differentiation, while we anticipated the reverse at expected inducible pop-DMS. Accordingly, we re-sequenced whole genomes of the same wild caught individuals we used for RRBS and calculated the degree of genomic differentiation per inducible pop-DMS as mean FST value (± 5 kb window) between populations. In line with our hypothesis, the populations from Kiel and Nynäshamn (decreased salinity) were genetically more differentiated at opposite inducible pop-DMS than at expected sites (δ.mean.FST = -0.014, P = 0.002; Figure 3A and 3C). A similar trend, yet not significant, was found between the populations from Kiel and Sylt (increased salinity: δ.mean.FST = -0.005, P = 0.153; Figure 3B and 3D). To understand if selection might have played a role in shaping the differences between increased and decreased salinity we tracked survival rates from fertilized eggs to the three months old offspring and compared them between treatment groups. Mortality differed significantly between the treatment groups (GLMM, X24 = 66.159, P < 0.001; Figure 4A, Supplementary Material: Table S3A) with increased mortality under increased salinity, while mortality under decreased salinity was generally low and did not differ from the control group (Figure 4A, Supplementary Material: Table S3A).

**Fig. 3:**
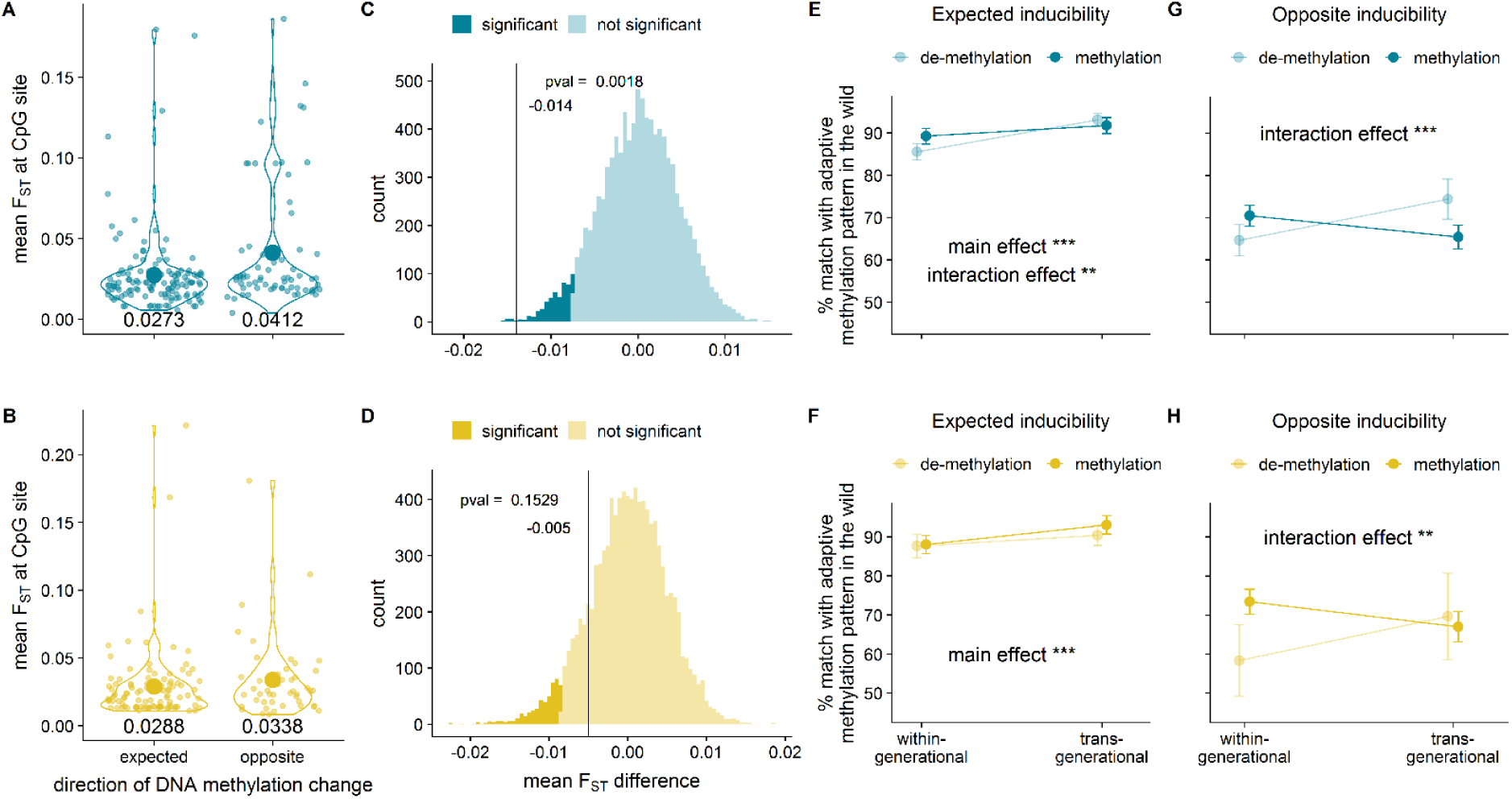
The duration of acclimation (within-vs. transgenerational) and level of genomic differentiation between the populations influence DNA methylation at inducible sites. Figure 3A and 3B show mean FST values for pop-DMS (with a ± 5 kb window) inducible under experimental salinity decrease (top, blue) and increase (bottom, yellow), that either shifted methylation levels towards the values observed in the field (expected) or the opposite direction (opposite). A randomization test (with 10,000 bootstraps) was performed for the difference between expected and opposite mean FST value (δ.mean.FST = ‘expected’ mean FST – ‘opposite’ mean FST; Figure 3C and 3D). Under the one tailed hypothesis of increased genetic differentiation at opposite sites and an alpha of 0.05 the P-value was calculated as values smaller than the true difference divided by 10,000 bootstraps. In Figure 3E-H the y-axis shows the percentage match between the within- and transgenerational acclimation group in relation to the methylation differentiation level found in natural populations at inducible pop-DMS. This value was obtained by calculating the difference between the methylation change in the experiment (meth.diff.exp in %; control vs. within-generational or control vs. transgenerational) and the difference in methylation between natural populations (meth.diff.wild in %) as δ.meth.diff = 100 – (meth.diff.wild – meth.diff.exp). Mean values ± 95% confidence interval are shown for within- and transgenerational acclimation to decreased and increased salinity at expected and opposite inducible sites. Colors refer to the direction of DNA methylation change (hypomethylation or hypermethylation). Values closer to 100 indicate a shift in methylation pattern towards adaptive methylation levels found in natural populations and stars indicate the significance level (P ≤ 0.001 ‘***’; P ≤ 0.01 ‘**’) for the comparison between within- and transgenerational acclimation. ‘Main effect’ refers to an effect of acclimation (within- or transgenerational) and ‘interaction effect’ to an interaction of acclimation and methylation direction (hypo- or hypermethylation).

**Fig. 4:**
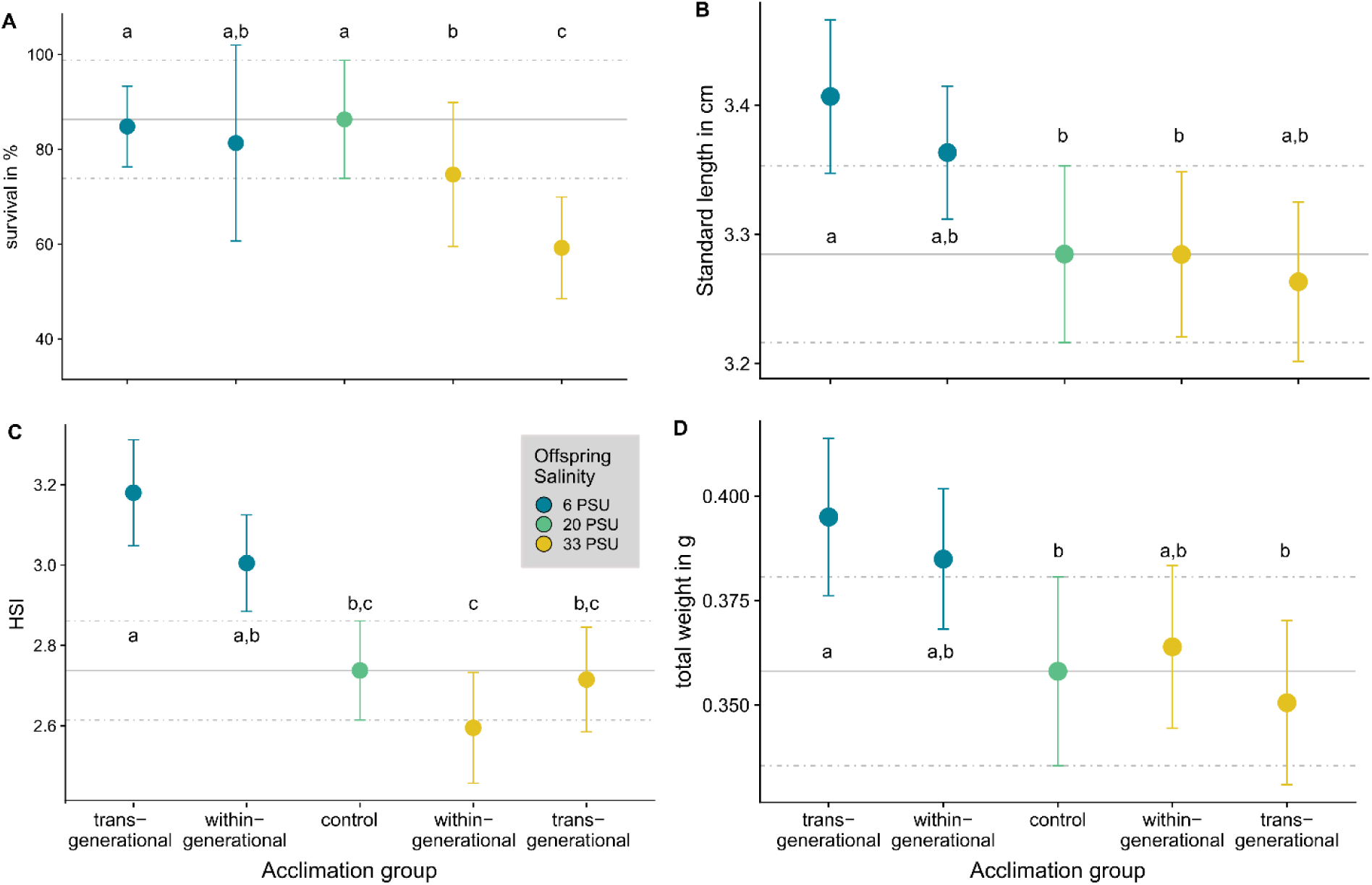
Effects of salinity acclimation on fitness correlated factors. For all five acclimation groups (control group (20 PSU), within- and transgenerational acclimation to 6 or 33 PSU) survival rates in percent (A), standard length in cm (B), hepatosomatic index (C) and total weight in g (D) are displayed. Letters indicate significant differences resulting from Tukey HSD post hoc tests (Supplementary Material: Table S3).

### Comparing within- and transgenerational acclimation effects on inducible DNA methylation

We then tested whether or not DNA methylation followed patterns predicted under adaptive transgenerational plasticity, namely that salinity acclimation over two instead of only one consecutive generations would enhance the similarity at inducible pop-DMS with patterns found among wild populations at corresponding salinities. To do so, we calculated the percentage match (δ.meth.diff, Figure 3E-H) between the within- and transgenerational acclimation groups in relation to the anticipated adaptive methylation level at inducible pop-DMS. In line with our hypothesis, we found that transgenerational compared to within-generation salinity manipulation increased the δ.meth.diff (for ‘expected’ inducible methylation: decreased salinity: F1,256 = 30.42, P < 0.001; increased salinity: F1,198 = 10.39, P = 0.001; Figure 3E and F). Remarkably, under decreased experimental salinity, we found an interaction of ‘methylation direction’ (hyper- or hypomethylation) and ‘acclimation’ (within- and transgenerational) affecting the δ.meth.diff (ANOVA, δ.meth.diff ∼ methylation direction * acclimation, F1,256 = 7.69, P = 0.006; Figure 3E). Here, transgenerational acclimation increased the similarity of hypomethylated sites to methylation levels found in natural populations, while hypermethylated sites showed similar values within- and across generations (Figure 3E). While for ‘expected’ inducible sites this effect was only present under decreased salinity, at ‘opposite’ inducible sites transgenerational acclimation to decreased and increased salinity enhanced the δ.meth.diff at hypomethylated sites (Figure 3G and H; ANOVA, δ.meth.diff ∼ methylation direction * acclimation, decreased salinity: F1,132 = 19.89, P < 0.001; increased salinity: F1,90 = 9.85, P = 0.002).

To infer the effect of DNA methylation differences on offspring fitness, we compared fitness correlated proxies between control, within- and transgenerational acclimation groups. Given that osmoregulation is energetically very costly we assessed the total weight, standard length (SDL) and the hepatosomatic index (HSI) as a proxy for energy reserves in form of liver glycogen storage. SDL (GLMM, X24 = 9.965, P = 0.041; Figure 4B, Supplementary Material: Table S3B) and total weight (GLMM, X24 = 11.518, P = 0.021; Figure 4D, Supplementary Material: Table S3D) differed only slightly between the treatment groups. Highly significant differences between the treatment groups were detected for the HIS (GLMM, X24 = 22.688, P < 0.001; Figure 4C, Supplementary Material: Table S3C) with an increased HSI under decreased salinity compared to the control group. This supports previous findings, showing that osmoregulation at 6 PSU is energetically less demanding (16, 17). Under increased salinity the HSI was lower compared to decreased salinity in the within-generation acclimation group, while a transgenerational acclimation to increased salinity partially removed this difference. Even though this effect was not significant, we observed a trend towards higher mean HSI in the transgenerational acclimation group compared to the within-generational acclimation group at the same salinity (Figure 4C, Supplementary Material: Table S3C).

## Discussion

This study investigated whether two postulated pathways of epigenetic inheritance (selection-based and detection-based) can be identified in natural populations, and whether the functions of the associated genes indicate a contribution to salinity adaptation in three-spined sticklebacks. Consistent with modeled selection dynamics for selection-based DNA methylation sites (7), we identified stable pop-DMS between populations enriched for osmoregulatory functions. As suggested by models on the role of epigenetic and genetic changes in adaptive evolution (5), such randomly originating DNA methylation sites could serve as potential targets for natural selection, resulting in differential DNA methylation of osmoregulatory genes between locally adapted populations. Phenotypic variation resulting from stable DNA methylation sites with expected epimutation rates of approximately 10-4 (estimated for A. thaliana (9)) could allow populations to explore the fitness landscape faster than under DNA sequence based genetic variation alone (mutation rate ∼10-8) (5, 23). Despite the random nature of such epimutations and the high gene flow among stickleback populations (15), the regionally synchronized signal of methylation differentiation in proximity to and within osmoregulatory genes suggests to be driven by local salinity environments. Furthermore, since local adaptation is 10-times more likely to involve changes at the gene expression than at the amino acid sequence level (24, 25), it is conceivable that differential DNA methylation and consequently different regulation of osmoregulatory genes may contribute to local adaptation to salinity. Remarkably, one of the top candidate genes differentially methylated between populations from 20 and 6 PSU was eda (Ectodysplasin A), a well-described gene involved in lateral plate formation (26). Salinity and calcium are significant drivers of plate morphology (27) in proposed conjunction with predation (28). Our findings suggest that repeated and parallel selection for the low plated eda allele in response to low saline habitats (29–31), including the Baltic Sea (15, 32), may not only be due to DNA base pair changes but also involve methylation mechanisms. Previous studies have suggested that energetic cost for Baltic sticklebacks increase with increasing difference between treatment and isosmotic salinity conditions (∼11 PSU (33)) (16, 17). In line with these findings, we observed many metabolic processes to be associated with stable pop-DMS under increased salinity, which is also reflected in the lower hepatosomatic index under increased salinity. Taken together, our results suggest that, along with genetic differentiation, the stable fraction of differentially methylated genes follow patterns consistent with local salinity adaptation across stickleback populations (Figure 2, Table 1; Supplementary Material: Figure S2, Table S2A and S2B). Previous studies have shown that patterns of local adaptation in DNA methylation can have a genomic basis in form of cis- and trans-acting genomic loci (22, 34). Whether the pop-DMS in this study represent an independent mechanism for local adaptation or whether they are a consequence of DNA-sequence based genetic differentiation would need further functional genomic investigation.

For the second postulated pathway on detection-based epigenetic inheritance (7), we identified a significantly higher number of experimentally inducible pop-DMS than expected by chance. Interestingly, multiple DMS associated with the same gene were mostly synchronized in their response (Table 1). Their association with different osmoregulatory genes compared to stable pop-DMS reflects a salinity-mediated plastic response potentially contributing to the molecular basis of adaptive phenotypic plasticity by allowing individuals to regulate their ion balance relative to the seawater medium instantaneously without requiring any further genetic adaptation at the population level. Interestingly, more than two-thirds of the inducible pop-DMS changed methylation to become more similar to the population locally adapted to the corresponding salinity. Remarkably, the similarity of these pop-DMS methylation levels between the naturally adapted and the experimentally acclimated population increased across generations. This strongly suggests that adaptive transgenerational plasticity via DNA methylation plays a role in salinity acclimation in wild animal populations that is changing incrementally with increasing exposure time to the environmental cue. Since we used a split-clutch design for the breeding experiment and mortality levels within the salinity decrease experiment were not different, we can assume that these groups have a similar genomic background. Therefore, we can rule out that most of the inducible pop-DMS simply follow patterns of genomic differentiation and assume that these sites are independently responding to the existing environmental pressure.

The induction of methylation sites has been widely discussed as potential buffer for environmental changes (11, 18, 35), which would allow populations to persist in the face of disturbances by moving phenotypes in the direction of optimal fitness faster than genetic changes alone (5, 23). Further work should be directed as to whether the inducible methylation changes would also become persistent and thus move effectively into the stable pop-DMS category, or whether these changes are reversible. We observed that the potential for adaptive transgenerational plasticity differed among methylation directions (hypo- and hypermethylated sites, Figure 3E, G and H), with a higher potential for transgenerational plasticity at hypomethylated sites. Generally, the spontaneous addition of a methyl-group to a cytosine is 2.5 times more likely than the removal (23) making a targeted de-methylation more difficult. In the transgenerational acclimation group, the methylation reprogramming including extensive methylation erasure and de novo methylation during gamete formation and zygote development could serve as a mechanistic basis to enhance de-methylation of CpG sites (36, 37).

The genetic background is considered one important source for epigenomic variation via cis- and trans-regulatory mechanisms (22, 38, 39). Genomic characterization around each experimentally inducible pop-DMS revealed a relationship between the level of population genetic differentiation (FST) and the propensity of the experimental population to approach the methylation level of the low salinity population (NYN) under experimental salinity decrease (Figure 3A and C). Here, experimentally induced DNA methylation becomes more similar to the methylation in natural populations only in genomic regions with low genetic differentiation. On the other hand, when experimentally induced methylation differences to the low salinity population increase (Figure 3A and C), this occurs in a more divergent genomic background, suggesting that the genome has undergone past selection leading to DNA-based local adaptation, rendering epigenetic modifications less relevant (5). This finding underlines the importance of incorporating the genomic background in the interpretation of DNA methylation patterns. Nevertheless, it remains to be tested what happens to all induced DNA methylation sites with selection over multiple generations.

Taken together, our study provides the first empirical evidence that stable and inducible DNA methylation exist in wild animal populations and follow predictions from evolutionary theory according to a selection- and a detection-based epigenetic pathway to promote adaptation to novel environments (5, 7) (Figure 5). The latter process, in particular, may provide an extremely fast (only two generations) mechanism to promote adaptive phenotypes which may be particularly important under predicted climate change (Figure 5). While the selection-based pathway follows the core assumptions of the Modern Synthesis (random variation and genetic inheritance), the detection-based pathway is better captured by the Extended Evolutionary Synthesis (non-random variation and inclusive inheritance). Because their evolutionary implications are very different, any future transgenerational or epigenetic studies should distinguish between these pathways, which will also contribute urgently needed data to the debate on an extended evolutionary synthesis. Whether epigenetic marks such as differentially methylated sites can permanently be attributed to one of the two categories or rather represent a continuum of stability levels and directionality will need further experimental testing over multiple generations.

**Fig. 5:**
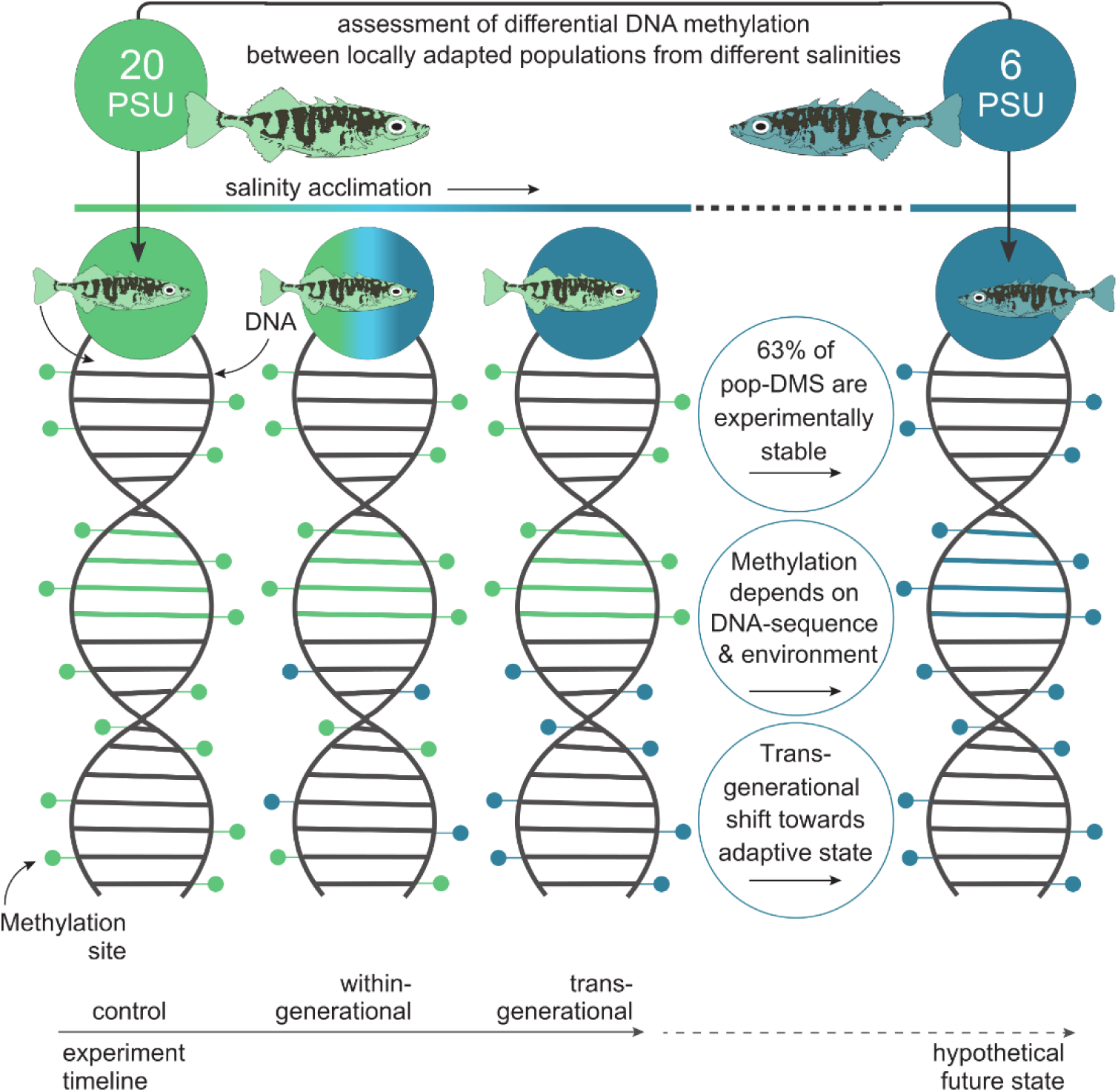
Graphical summary of experimental design and main results. We used the Baltic Sea salinity gradient to study the role of DNA methylation in local salinity adaptation and the response to salinity change in a space-for-time approach. To assess the potential future acclimatization and adaptation processes of the natural stickleback population from 20 PSU (Kiel / green) to the predicted desalination(63), we compared differences in DNA methylation at CpG sites between wild caught and laboratory bred sticklebacks. Following the experiment timeline (bottom), we compared methylation levels of the experimental control group from 20 PSU, to within- and transgenerational acclimation of 20 PSU sticklebacks to 6 PSU (DNA from left to right). The population locally adapted to 6 PSU serves as the hypothetical future state in which salinities will decrease (blue, DNA on the right). The three main results are written in the circles with schematically and horizontally corresponding DNA methylation changes. (i) 63% of the DMS between the populations remained stable under experimental salinity change. (ii) The direction of experimental methylation change was dependent not only on the treatment but also on the degree of genetic differentiation between the populations (see Figure 3A-D for results). (iii) Transgenerational salinity acclimation shifted DNA methylation patterns closer to the anticipated adaptive state found in the hypothetical future population (see Figure 3E-H for results). For clarity, only one (6 PSU) of the two foreign salinity regimes tested (6 and 33 PSU) is shown, indicated by the yellow fish on the top left (see Figure 1 for full experimental design).

## Materials and Methods

### Animal Welfare

All catches were performed under legal authorization issued by the German ‘Ministry of Energy Transition, Agriculture, Environment, Nature and Digitalization’ in Schleswig-Holstein (MELUR – V242-7224.121-19), by the Danish ‘Ministry of Food, Agriculture and Fisheries of Denmark’ (Case no: 14-7410-000227), by the Estonian ‘Ministry of the Environment’ (Keskkonnaministeerium - eripüügiluba nr 28/2014) and by the Swedish Sea and Water Authority (Havs och Vattenmyndigheten). Ethical permission for the experiments required by German law was given by the MELUR: V312-7224.121-19).

### Survey and experimental design

For the field survey, we collected juvenile three-spined sticklebacks (Gasterosteus aculeatus; 31.68 ± 14.25 mm) from three different salinity regimes inside and outside the Baltic Sea (Sylt (SYL), Germany (55°00’58.3’’N, 8°26’22.0’’E), 33 PSU, N = 16; Kiel (KIE), Germany (54°26’11.8 N 10°10’20.2 E), 20 PSU, N = 16; Nynäshamn (NYN), Sweden (58°52’44.7 N 17°56’06.2 E), 6PSU, N = 16) in September 2014. Once collected, fish were immediately euthanized using tricaine methane sulfonate solution (MS222), photographed, measured (length and total weight) and stored in RNA-later (24h at 7°C, afterwards at -20°C). A cut along the ventral side ensured that the RNA-later solution would diffuse into all tissues. Conserved specimen were later dissected in the lab and gill tissue was separated, since gills represent a main osmoregulatory organ in fishes. For the acclimation experiment, we collected live adult fish from Kiel (20 PSU), which were crossed in our facilities at GEOMAR to obtain ten F1 laboratory bred families, herein referred to as ‘parental generation’. At nine months post-hatch we split each family into three salinity treatment groups of 10 fish each: one at 33 PSU, one at 6 PSU, and one control group at 20 PSU. The salinity transition was performed within 10 days by 3 PSU steps every second day. Over the entire time each group was fed ad libitum and kept in a 20-L aquarium connected to one of three filter tanks per salinity treatment. After 5 months under treatment conditions, six pure crosses per salinity treatment group were performed in vitro, herein referred to as ‘offspring generation’ (F2). Offspring and parental generation were kept at 18 °C water temperature and a 15:9 light/dark cycle (L/D). The last eight weeks before performing the F2 crosses, the F1 generation underwent an artificial winter to trigger reproduction (2 weeks at 12 °C, 12:12 L/D; 4 weeks 6 °C, 8:16 L/D, 2 weeks at 12 °C, 12:12 L/D). Upon fertilization, clutches were split and separated into different treatments (Figure 1). At three months post-hatch, laboratory bred F2 sticklebacks were euthanized using MS222, photographed, dissected and their gill tissue was stored in RNA-later. The age at sampling matched the estimated age of the wild caught juveniles (3 months). Additionally, to the 48 wild caught individuals from Kiel, Nynäshamn and Sylt that were used in above field survey, we sequenced whole genomes from gill tissue of an additional three populations of sticklebacks, namely from Falsterbo, Sweden (55°24’46.6 N 12°55’52.3 E; 10 PSU; N = 16), Letipea (59°33’07.6 N 26°36’29.7 E; 4 PSU; N = 16) and Barsta (62°51’47.1 N 18°23’51.0 E; 5 PSU; N = 16).

### Mortality and hepatosomatic index

Mortality was monitored throughout the experiment to account for possible non-random mortality. Three months post-hatch, we assessed the standard length (SDL), total weight and liver weight of the experimental F2 generation and calculated the hepatosomatic index (HSI = liver weight / total weight * 100), which is a proxy for energy reserves in form of glycogen storage in the liver. We analyzed the effect of treatment (5 treatment groups, Figure 1) on the survival rate per family as a ratio of ‘alive’ vs ‘dead’ fish, using glmer implemented in the R package ‘lme4’ (40) with Binomial error and ‘crossing’ as well as ‘climate chamber’ as random effects. The effect of treatment on HSI, SDL and total weight was analyzed fitting three individual linear mixed effect models, using lmer in ‘lme4’ (40) with Gaussian error and ‘crossing’ as well as ‘tank’ nested within ‘climate chamber’ as random effects. Tukey HSD post hoc tests were run using the glht function implemented in the package ‘multicomp’ (41) to identify significant differences between treatment groups.

### DNA extraction

For the field survey, DNA extraction of gill tissue (N = 16 individuals per population) was performed using the DNeasy Blood & Tissue Kit (Qiagen). Further purification of the extracted DNA was done with NucleoSpin® gDNA Clean-up (Macherey-Nagel). For laboratory bred F2 offspring of the two-generation acclimation experiment, dual extraction of whole RNA and DNA was performed from gill tissue (N = 11-12 individuals per treatment group, Figure 1) stored in RNAlater using the AllPrep DNA/RNA mini kit (Qiagen). Purity and quality of the extracted DNA was estimated using a NanoDrop ND-1000 spectrophotometer (Thermo Fisher Scientific) and a standard agarose gel (1% Agarose/TAE). DNA concentration was assessed using the Qubit® 2.0 Fluorometer (Thermo Fisher Scientific). To obtain a balanced sex ratio (50/50), we determined the gender of the individuals using a sex-specific genetic polymorphism in isocitrate dehydrogenase (IDH) with a modified protocol from Peichel et al. (2004) (42). For the PCR (settings: once 94°C for 3 minutes; 30 cycles of 94°C for 30 sec, 54°C for 20 sec, 72°C for 30 sec; once 72°C for 5 minutes), 1 µL forward and reverse primer (5µM) was used with 4.9 µL water, 1 µL 10x buffer, 1 µL dNTPs (0.5 µM), and 0.1 µL DreamTaq (5 U/µL). The resulting PCR products were visualized with a capillary electrophoresis on the 3100 ABI sequencer and a LIZ500 size standard. While males show a heterogametic signal with two bands (at approximately 300 bp and 270 bp), females lack the band at 270 bp.

### Library preparation and sequencing (Whole genome sequencing, WGS)

For whole genome sequencing, the ‘TruSeq Nano DNA’ (Illumina) library preparation kit was used according to the manufacturer’s protocol by the Sequencing Facility of the IKMB, University of Kiel. Ultrasonication was conducted with a ‘Covaris E220’ (Covaris) to shear the input DNA (100 ng per sample and 350 bp insert size). Before the enrichment with a PCR step (8 cycles), fragmented and bead purified DNA was ligated with adenylate at the blunt 3’ ends (End repair and A-tailing) and with indexing adapters. Fragments were cleaned with MagSi-NGS Prep Plus Beads (Steinbrenner). Paired-end sequencing of the quality-controlled and multiplexed libraries was performed on the Illumina Hiseq 4000 platform (2 x 150 bp reads).

### Quality assessment, data filtering and mapping (WGS)

The command line tools of Picard v.2.7.1 (Broad Institute 2016) were used to (i) reformat the Fastq to uBAM file format and to add further values (read group etc.) to the SAM header using FastqToSam), to (ii) mark the location of adapter sequences using MarkIlluminaAdapters, and to (iii) reconvert the sequences to Fastq format with SamToFastq. The stickleback genome (here Broad/gasAcu1) was indexed with bwa index and used as a reference for the mapping with bwa mem (43) v.07.12-r1044. To retain the meta-information from the uBAMs we used MergeBamAlignment. Picard was also used to identify duplicates with MarkDuplicates. Basic statistics were generated with CollectWgsMetrics, CollectInsertSizeMetrics and AlignmentSummaryMetrics and summarized with MultiQC v1.0.dev0 (44). A total number of 4,463,070,154 high-quality reads (mapping quality > Q20) was mapped resulting in a mean depth of 13.84x (sd. 2.02x) and mean insert size 383.07 bp (sd. 9.40 bp, Supplementary Table S3). GATK v 3.7 HaplotypeCaller (45) was run to determine the likelihoods of the haplotypes per sample, i.e. to call SNPs and indels, which were than processed with GenotypeGVCFs for a joint genotyping. SNPs were selected using hard filters for quality and extracted from the raw genotypes with a combination of the SelectVariants, VariantsToTable and VariantFiltration commands. VCFtools (46) was used in a next step, removing SNPs with a minimum quality score (minQ) below 20 and a minor allele frequency (maf) greater than or equal 0.0049.

### Library preparation and sequencing (reduced representation bisulfite sequencing, RRBS)

The library preparation for methylation analyses followed the Smallwood and Kelsey reduced representation bisulfite sequencing (RRBS) protocol (47). A total of 100-250 ng purified DNA was digested with the methylation-insensitive MspI restriction enzyme, which cuts at the “CCGG” motif and thereby enriches for CpG regions. DNA end-repair and A-tailing was conducted and un-tailed CEGX spike-in controls (Cambridge Epigenetix) were added. These are DNA oligos of known sequence and with known cytosine modification, which can be used for downstream assessment of bisulfite conversion efficiency. After adapter ligation, bisulfite conversion was conducted using the EZ-96 DNA Methylation-Gold Kit (Zymo Research) according to the manufacturer’s protocol. PCR amplification with 19 cycles were performed. Quality control of purified PCR products was performed on a 2200 TapeStation System (Agilent) and high-quality libraries were pooled and diversified with 15% PhiX. Single-end sequencing with 100 bp read length was conducted on a HiSeq 2500 sequencer (Illumina).

### Quality assessment, data filtering and mapping (RRBS)

In total 106 individuals (48 wild caught and 58 experimental fish) of balanced sex ratio were DNA sequenced at an average of 19.8 ± 3.5 million reads for experimental fish and 11.4 ± 2.1 million reads for wild-caught fish (Supplementary Table S4). De-multiplexed fastq files were quality checked using FastQC v0.11.5 (48) and Multiqc v1.3 (44). Adapters were removed with cutadapt v1.9.1 (49) using multiple adapter sequences (NNAGATCGGAAGAGCACAC, AGATCGGAAGAGCACAC, ATCGGAAGAGCACAC) with a minimum overlap of one base pair between adapter and read. This was necessary to remove primer dimers and avoid false methylation calls systematically caused by the RRBS end-repair step during library preparation, if the end repair step adds artificial cytosins. Simultaneously, cutadapt was used to trim low quality bases (-q 20) from the 3’-end and remove trimmed reads shorter than 10 bases. An air bubble during sequencing caused the bases 66-72 of ten tiles of one lane (affecting 12 individuals) to have low quality values, which were removed in a custom awk script. Two poor quality individuals (a Sylt and a Nynäshamn female) did not meet our strict quality requirements (e.g.: ≥ 5 million reads, mapping efficiency > 52%) and showed biases in the proportion of bases per position compared to other individuals (plot in fastqc per base sequence content). Therefore, we excluded these two libraries from downstream analysis resulting in 15 instead of 16 individuals from Sylt and Nynäshamn (Figure 1). Bisulfite conversion efficiency was assessed from the spike-in controls (Cambridge Epigenetix), using the cegxQC software (50). Overall conversion levels were 2.4 ± 1.8% conversion of methylated cytosines and 99.6 ± 0.5% conversion of un-methylated cytosines, which is in line with expected conversion rates (Supplementary Table S4). We used Bismark v0.17.0 (51) to index the UCSC stickleback reference genome (Broad/gasAcu1) and to generate the bisulfite alignments with Bowtie2 v2.3.3 at default settings. Bismark was also used to extract the methylation calls. Average mapping efficiency was 63.7 ± 2.4% (Supplementary Table S4).

### Identification of differentially methylated sites

The methylation calls were analyzed in R v3.4.1 (52) using the package methylKit v1.3.8 (53). CpG loci were filtered for a minimum coverage of 10 reads per site. To account for potential PCR bias, we additionally excluded all sites in the 99.9th percentile of coverage. To improve the methylation estimates, we corrected for SNPs, which could have led to a wrong methylation call. The excluded positions were derived with custom written perl scripts from C-to-T and G-to-A-SNPs with genotype quality of 20 and a minimum allele frequency of 0.005 (see above) from the 96 wild caught individuals with a combination of custom written Perl and R-scripts using packages from methylkit (53) and GenomicRanges (54) (Supplementary File Summary_scripts.txt). After normalizing coverage values between samples, using normalizeCoverage implemented in methylKit, we excluded all sites that were present in fewer than 9 individuals per treatment group from downstream analysis. As previously shown, sex specific methylation affects < 0.1% of CpG sites on autosomal chromosomes, but > 5% of CpGs on the sex chromosome (19). Therefore, to exclude a potential sex bias, we removed all CpG sites located on the sex chromosomes (chromosome 19), resulting in a high-quality dataset with 525,985 CpG sites. Finally, by checking the first six principal components of the resulting PCA and running an ANOVA on the filtered dataset, we confirmed the absence of an effect of sex on global methylation pattern (F124,1 = 2.611, P = 0.109). However, the PCA revealed a bias in methylation pattern by families over all experimental groups. Therefore, to identify differentially methylated CpG sites (DMS) between treatment groups, we performed pairwise comparisons (Supplementary Table S5) fitting a logistic regression model per CpG site with calculateDiffMeth in methylKit using family as covariate for the experimental groups. A Chi-square test was applied to assess significance levels of DMS and P-values were corrected to q-values for multiple testing using the SLIM (sliding linear model) method (55). Additionally, we accounted for multiple use of groups in pairwise comparisons and adjusted the alpha for the q-value according to Bonferroni correction to 0.0125 (= 0.05 / 4). Ultimately, CpG sites were considered to be differentially methylated with a q-value < 0.0125 and a minimum weighted mean methylation difference of 15%. To ensure that the DMS obtained are not laboratory artefacts, we used calculateDiffMeth implemented in methylKit and compared the wild population from Kiel to the experimental control group (Kiel population from 20 PSU at 20 PSU). The resulting 11,828 DMS were excluded from the DMS obtained by the pairwise comparisons mentioned above (Supplementary Table S5). DMS were plotted across the genome for the comparison between Kiel vs Nynäshamn (20 vs. 6 PSU, blue fish) and Kiel vs Sylt (20 vs. 33 PSU, yellow fish) using ggplot2 (56) und hypoimg (57) (Supplementary Figure S1).

### Assessment of inducibility and gene association of DMS

By comparing wild caught individuals from the mid salinity population (20 PSU, KIE) to the populations sampled at low (6 PSU, NYN) and high (33 PSU, SYL) salinity in the field, we obtained 1,470 (KIE-NYN) and 1,158 (KIE-SYL) pairwise pop-DMS. We first tested whether these pop-DMS distinguishing natural populations are inducible or stable at the respective salinity in the experiment. A pop-DMS was considered stable when the within- and the transgenerational acclimation group did not significantly differ in methylation to the control group (q-value ≥ 0.0125). On the other hand, pop-DMS were considered inducible when at least one of the acclimation groups was differentially methylated compared to the control group (q-value < 0.0125; methylation difference ≥ 15%). Pop-DMS with a significant q-value not exceeding the threshold of differential DNA methylation (15%) will be referred to as ‘inconclusive’ hereafter. We used a randomization test to ensure that the number of inducible sites obtained did not occur by chance. To this end, we randomly sampled 1,470 (KIE-NYN) and 1,158 (KIE-SYL) pop-DMS from the complete dataset (1,000 replicates). A Chi-square test was used to assess whether our observed number of inducible, stable and inconclusive sites differs from a random set of sites (averaged over replicates). Finally, we tested whether the weighted mean methylation difference (meth.diff, in percentage) between wild populations matches the inducible methylation difference by subtracting the ‘meth.diff’ in the experiment (exp) from the ‘meth.diff’ between wild caught populations (wild):

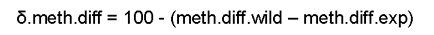

As we subtracted this difference from 100, values closer to 100 indicated higher similarity of experimentally inducible methylation with the postulated adaptive DNA methylation pattern in natural populations. By comparing the ‘δ.meth.diff’ for within- and transgenerational acclimation using an ANOVA, we can assess whether there is a difference in inducibility of methylation to match patterns found in wild caught populations. All analyses were run separately for decreased (6 PSU; KIE-NYN) and increased (33 PSU; KIE-SYL) salinity.

In order to detect potential functional associations of the observed changes in DNA methylation state, we classified the genomic region of a pop-DMS based on their nearest transcription start site (TSS) using annotateWithGeneParts and getAssociationWithTSS implemented in genomation v1.4.2 (58). We distinguished between promoter (1500 bp upstream and 500 bp downstream of TSS), exon, intron and intergenic regions. To be associated to a gene, the pop-DMS had to be either inside the gene or, if intergenic, not further than 10 kb away from the TSS. We excluded three pop-DMS that were on a different reference scaffold then the gene they were associated to on the chrUn linkage group (that merges scaffolds into one large artificial chromosome). Using the genes with associated pop-DMS, we applied a conditional hypergeometric GO term enrichment analysis (FDR corrected P-value ≤ 0.05) with the ensembl stickleback annotation dataset ‘gaculeatus_gene_ensembl’ and all genes that were associated to any sequenced CpG site as universe. We identified overrepresented biological processes, molecular functions and cellular components using the packages GOstats v2.5 (59) and GSEABase v1.46 (60) and corrected for multiple testing using the FDR method implemented in goEnrichment v1.0 (61) in R v3.6 (52). Figures were produced using ggplot2 v3.2 (56).

### Estimation of DNA sequence based genetic differentiation at differentially methylated sites

In order to evaluate the genetic differentiation up- and downstream (in sum 10 kb) of the pop-DMS position, we calculated the mean FST values (≤ 60% missing data and depth ≥ 5) from whole genome sequencing data of the exact same individuals with vcftools v.0.1.15 (62). We hypothesized that inducible CpG positions matching the methylation difference expected from the profile of the wild populations are genetically more similar between the populations than sites that changed in the opposite direction. To test this one-tailed hypothesis we applied a randomization test (with 10,000 bootstraps) on the mean FST difference between the two groups (expected and opposite):

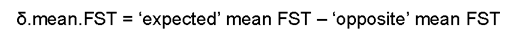

We plotted the 10,000 delta mean FST values and calculated a P-value by dividing the proportion of values smaller than the true difference by the number of bootstraps. Figures were produced using ggplot2 v3.2 (56).

## Supporting information

Summary of Scripts

## Acknowledgments

Many thanks to Fabian Wendt, Jakob Gismann, Linda Sartoris, Christoph Giez, Zuzanna Zagrodzka, Laura Niewendick, Max Bettendorff, Bastian Poerschke, and Tobias Strickmann for taking care of the experimental animals and assisting the sampling procedures. Thanks to Florian Rüppel for planning and constructing the recirculating aquaria systems. We thank the many field helpers for catching wild sticklebacks. Many thanks also to Susanne Landis who visualized our ideas in Figure 1 and 5. We kindly acknowledge the financial support for the BAMBI project (Grant Agreement number: call 2012-76) by BONUS, the joint Baltic Sea research program, funded by the European Union and the Federal Ministry of Education and Research in Germany to TBHR and CE (reference number 03F0680A). Further, BSM’s research was generously supported by the Excellence Cluster ‘The Future Ocean’ (EXC 80) and the sequencing partly by the Excellence Cluster ‘Inflammation at Interfaces’ (EXC 306) and ‘Origins and Function of Metaorganisms’ (CRC1182), subprojects Z3 & INF, by the Deutsche Forschungsgemeinschaft (DFG). Both ‘The Future Ocean’ and ‘Inflammation at Interfaces’ are funded within the framework of the Excellence Initiative by the Deutsche Forschungsgemeinschaft (DFG) on behalf of the German federal and state governments.

## Author contributions

CE and TBHR conceived and designed the study with contributions for the bisulfite sequencing strategy from BSM. BSM and MJH planned and carried out the fieldwork at the German locations, BSM supervised the sampling in Estonia and Sweden, which was carried out by a great team of the BONUS-BAMBI project. MJH, with the help from CE and TBHR, planned and supervised the breeding and acclimatization experiment. MJH and BSM conducted wet laboratory work (DNA extractions and quality assessment). MPH and RH and conducted the library preparation and sequencing. MJH and BSM analyzed the data and drafted the manuscript together with equal contributions. All co-authors discussed and interpreted the results and contributed to the final version of the manuscript.

## Competing interests

The authors declare no competing interests.

## Data Availability

Fastq raw reads of genomes and methylation sequencing will be deposited in GenBank. Raw data on mortality and phenotypic effects are available at PANGAEA (https://doi.org/10.1594/pangaea.892493).

## Notes

#### Summary of Updates

Results on phenotypic effects (effects of salinity on standard length, total weight and hepatosomatic index) and mortality were added.

