## Supplementary Information for "Two different epigenetic pathways detected in wild three-spined sticklebacks are involved in salinity adaptation"

20 **Figure S1:**

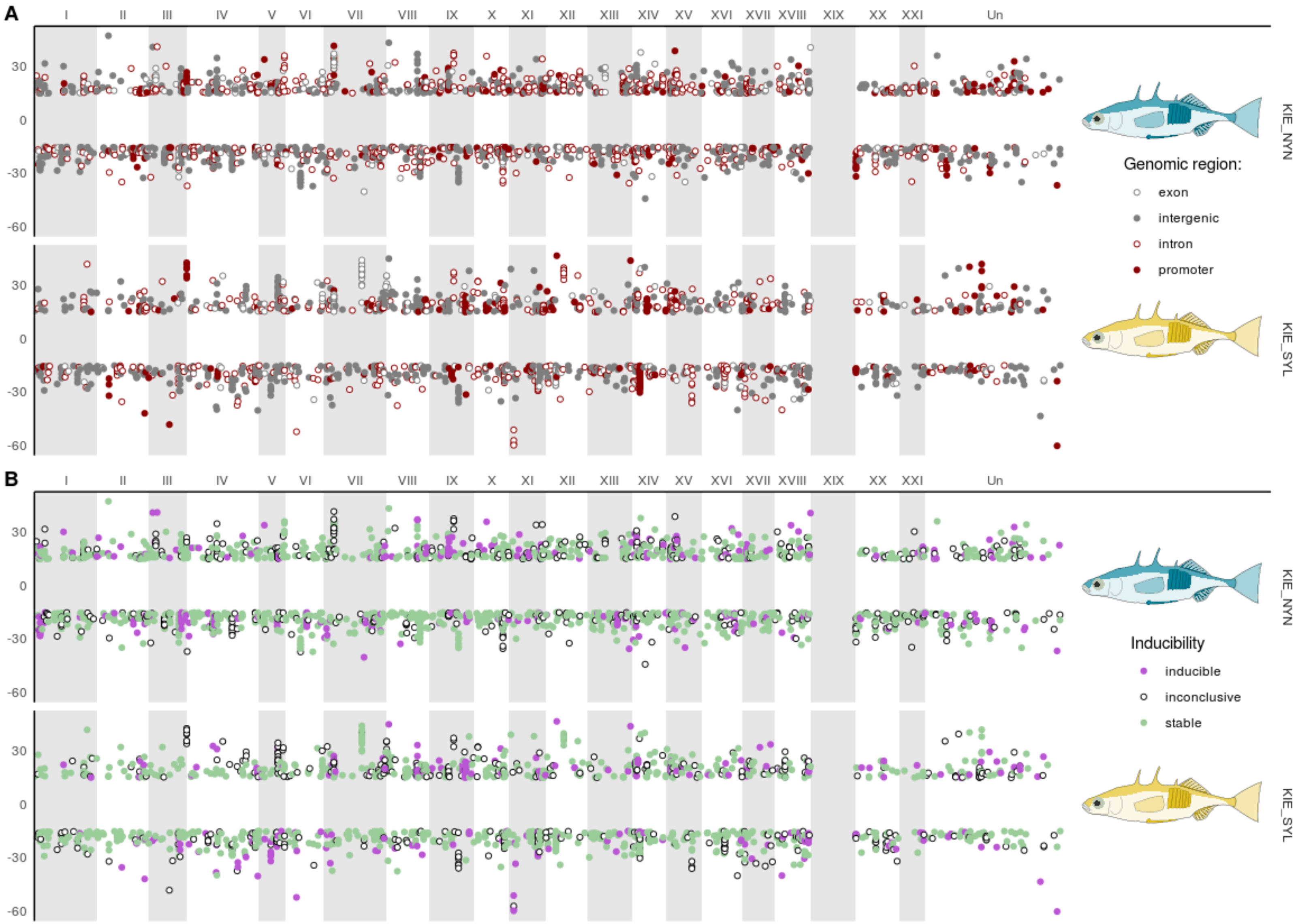

21

**Fig. S1:** Significant DMS throughout the genome for comparison between Kiel vs. Nynäshamn (20 vs. 6 PSU, blue fish) and Kiel vs. Sylt (20 vs. 33 PSU, yellow fish). Each DMS is one dot and the y axis represents the methylation difference in percent (%). DMS $> 0$ are hypermethylated in Nynäshamn / Sylt compared to Kiel and DMS $< 0$ are hypomethylated in Nynäshamn / Sylt compared to Kiel. No DMS are shown on the sex chromosome (number 19), since this was filtered out to reduce the sex bias (see Methods for details). The “chromosome unknown” (Un) represents a size-sorted collection of un-assembled scaffolds. Figure S1A and S1B show the same DMS. In S1A the DMS are colored according to their genomic location (exon, intergenic, intron, promoter) and in S1B according to their stability (stable, inconclusive, experimentally inducible).

**Figure S2:**

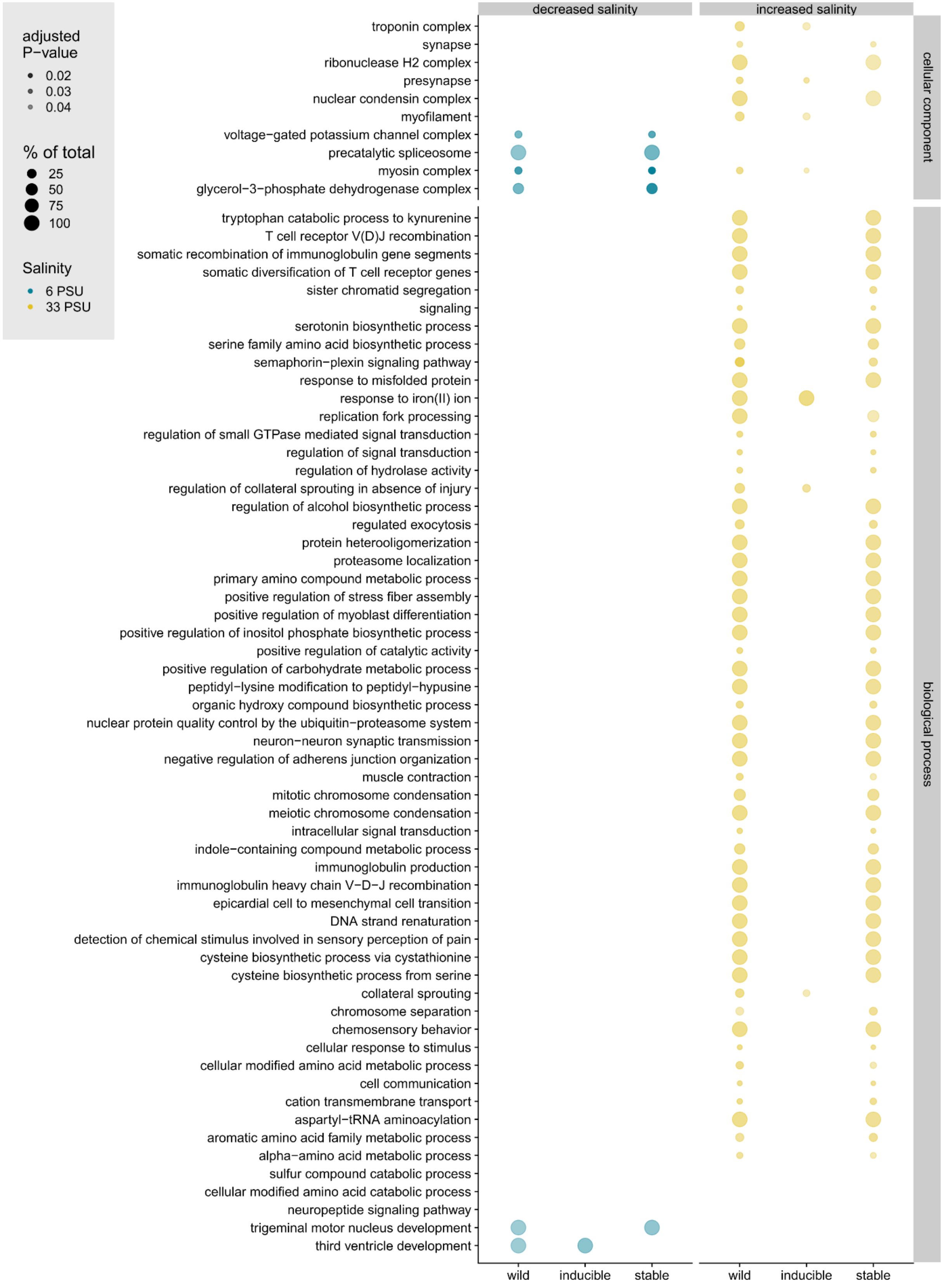
adjusted P−value
0.02
0.03
0.04
% of total
25
50
75
100
Salinity
6 PSU
33 PSU
decreased salinity
increased salinity
cellular component
biological process
troponin complex
synapse
ribonuclease H2 complex
presynapse
nuclear condensin complex
myofilament
voltage−gated potassium channel complex
precatalytic spliceosome
myosin complex
glycerol−3−phosphate dehydrogenase complex
tryptophan catabolic process to kynurenine
T cell receptor V(D)J recombination
somatic recombination of immunoglobulin gene segments
somatic diversification of T cell receptor genes
sister chromatid segregation
signaling
serotonin biosynthetic process
serine family amino acid biosynthetic process
semaphorin−plexin signaling pathway
response to misfolded protein
response to iron(II) ion
replication fork processing
regulation of small GTPase mediated signal transduction
regulation of signal transduction
regulation of hydrolase activity
regulation of collateral sprouting in absence of injury
regulation of alcohol biosynthetic process
regulated exocytosis
protein heterooligomerization
proteasome localization
primary amino compound metabolic process
positive regulation of stress fiber assembly
positive regulation of myoblast differentiation
positive regulation of inositol phosphate biosynthetic process
positive regulation of catalytic activity
positive regulation of carbohydrate metabolic process
peptidyl−lysine modification to peptidyl−hypusine
organic hydroxy compound biosynthetic process
nuclear protein quality control by the ubiquitin−proteasome system
neuron−neuron synaptic transmission
negative regulation of adherens junction organization
muscle contraction
mitotic chromosome condensation
meiotic chromosome condensation
intracellular signal transduction
indole−containing compound metabolic process
immunoglobulin production
immunoglobulin heavy chain V−D−J recombination
epicardial cell to mesenchymal cell transition
DNA strand renaturation
detection of chemical stimulus involved in sensory perception of pain
cysteine biosynthetic process via cystathionine
cysteine biosynthetic process from serine
collateral sprouting
chromosome separation
chemosensory behavior
cellular response to stimulus
cellular modified amino acid metabolic process
cell communication
cation transmembrane transport
aspartyl−tRNA aminoacylation
aromatic amino acid family metabolic process
alpha−amino acid metabolic process
sulfur compound catabolic process
cellular modified amino acid catabolic process
neuropeptide signaling pathway
trigeminal motor nucleus development
third ventricle development
wild
inducible
stable
wild
inducible
stable

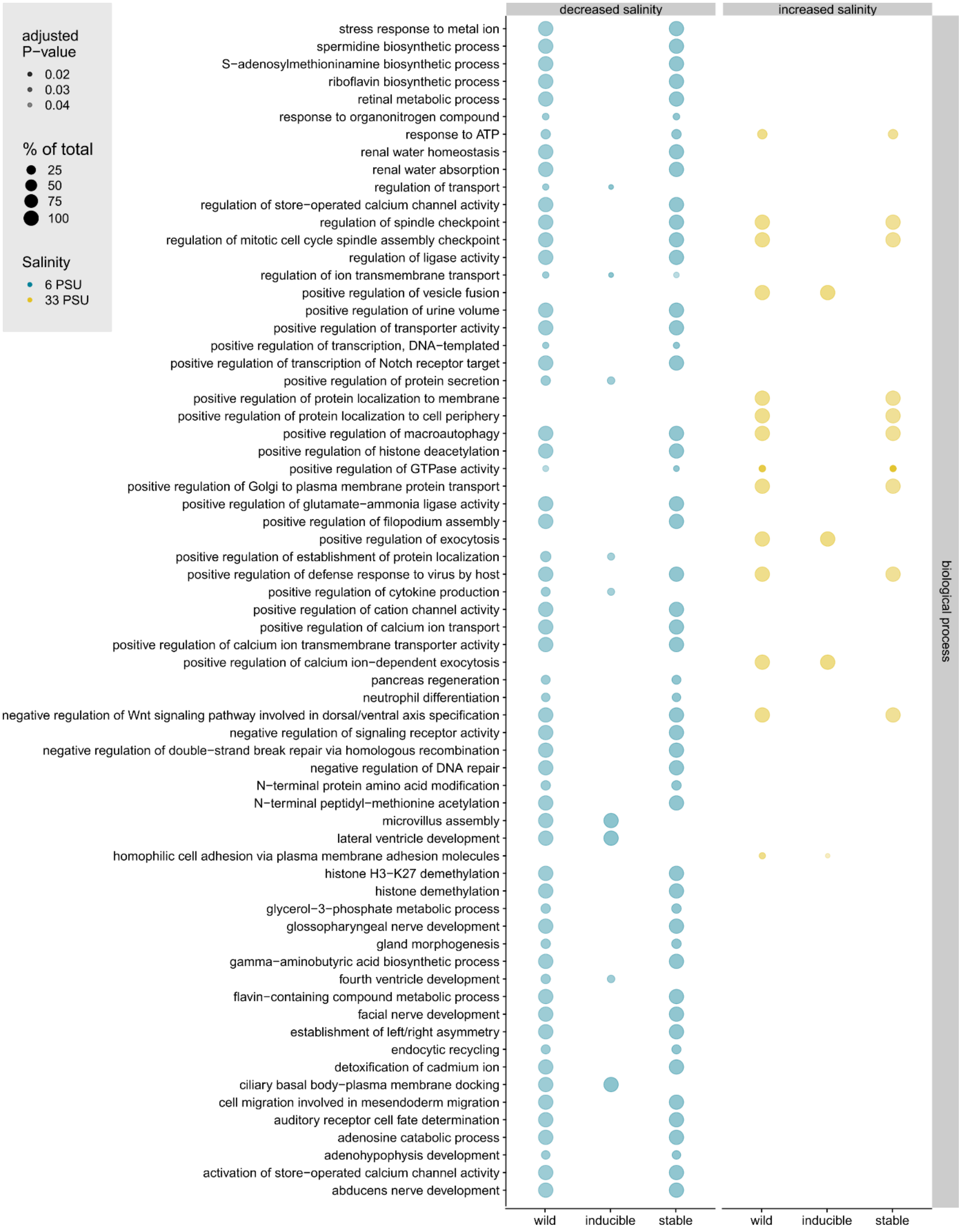
adjusted P−value
0.02
0.03
0.04
% of total
25
50
75
100
Salinity
6 PSU
33 PSU
decreased salinity
increased salinity
stress response to metal ion
spermidine biosynthetic process
S−adenosylmethioninamine biosynthetic process
riboflavin biosynthetic process
retinal metabolic process
response to organonitrogen compound
response to ATP
renal water homeostasis
renal water absorption
regulation of transport
regulation of store−operated calcium channel activity
regulation of spindle checkpoint
regulation of mitotic cell cycle spindle assembly checkpoint
regulation of ligase activity
regulation of ion transmembrane transport
positive regulation of vesicle fusion
positive regulation of urine volume
positive regulation of transporter activity
positive regulation of transcription, DNA−templated
positive regulation of transcription of Notch receptor target
positive regulation of protein secretion
positive regulation of protein localization to membrane
positive regulation of protein localization to cell periphery
positive regulation of macroautophagy
positive regulation of histone deacetylation
positive regulation of GTPase activity
positive regulation of Golgi to plasma membrane protein transport
positive regulation of glutamate−ammonia ligase activity
positive regulation of filopodium assembly
positive regulation of exocytosis
positive regulation of establishment of protein localization
positive regulation of defense response to virus by host
positive regulation of cytokine production
positive regulation of cation channel activity
positive regulation of calcium ion transport
positive regulation of calcium ion transmembrane transporter activity
positive regulation of calcium ion−dependent exocytosis
pancreas regeneration
neutrophil differentiation
negative regulation of Wnt signaling pathway involved in dorsal/ventral axis specification
negative regulation of signaling receptor activity
negative regulation of double−strand break repair via homologous recombination
negative regulation of DNA repair
N−terminal protein amino acid modification
N−terminal peptidyl−methionine acetylation
microvillus assembly
lateral ventricle development
homophilic cell adhesion via plasma membrane adhesion molecules
histone H3−K27 demethylation
histone demethylation
glycerol−3−phosphate metabolic process
glossopharyngeal nerve development
gland morphogenesis
gamma−aminobutyric acid biosynthetic process
fourth ventricle development
flavin−containing compound metabolic process
facial nerve development
establishment of left/right asymmetry
endocytic recycling
detoxification of cadmium ion
ciliary basal body−plasma membrane docking
cell migration involved in mesendoderm migration
auditory receptor cell fate determination
adenosine catabolic process
adenohypophysis development
activation of store−operated calcium channel activity
abducens nerve development
wild
inducible
stable
wild
inducible
stable
biological process
33

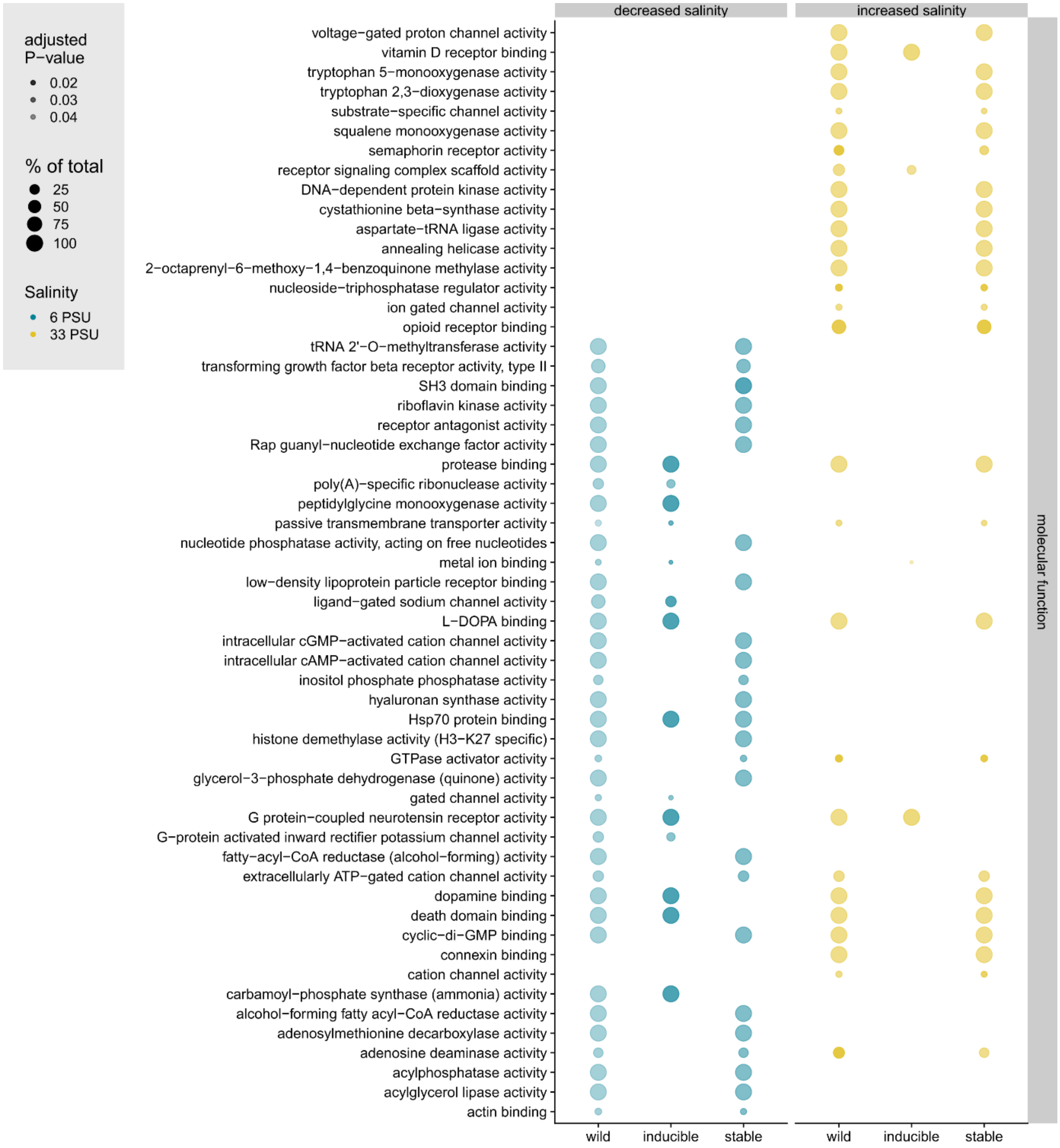

**Fig. S2:** Gene ontology (GO) terms for biological processes, cellular components and molecular functions under salinity increase (yellow, 20 vs. 33 PSU) and decrease (blue, 20 vs. 6 PSU) associated with differentially methylated sites between populations (pop-DMS) are presented. The graph is split into GO terms associated with pop-DMS from natural stickleback populations across a salinity cline (wild) and their experimental inducibility (inducible and stable) in a two-generation acclimation experiment. The size of the circles refers to the number of genes of this term present in our groups (in %) and the transparency to the *P*-value (darker circles refer to a lower *P*-value).

### Supplementary Tables

**Table S1:** Relative distribution of DMS among genomic features. Given are DMS between field populations (KIE vs. NYN for 20 vs. 6 PSU and KIE vs. SYL for 20 vs. 33 PSU), as well as experimentally inducible and stable sites for increased and decreased salinity separately. 525,985 CpGs were sequenced.

| Comparison | | Genomic feature (%) | | | |
|---|---|---|---|---|---|
| | | exon | intergenic | intron | promoter |
| DMS under salinity decrease | KIE vs. NYN | 11.2 | 45.9 | 31.5 | 11.4 |
| | inducible | 9.1 | 49.5 | 30.3 | 11.1 |
| | stable | 11.8 | 44.8 | 33.5 | 9.9 |
| DMS under salinity increase | KIE vs. SYL | 13.6 | 43.3 | 30.1 | 13.0 |
| | inducible | 9.5 | 42.5 | 31.8 | 16.2 |
| | stable | 13.0 | 42.9 | 32.6 | 11.5 |
| CpGs sequenced in total | | 15.1 | 40.7 | 25.4 | 18.8 |

**Table S2A:** Differentially methylated genes between populations from Kiel (20 PSU) and Nynäshamn (6 PSU): For genes associated with DMS, Ensembl gene ID and gene name as well as the position on the chromosome are listed. The numbers refer to the numbers of DMS in the population comparison (wild), these DMS were classified into ‘inducible’, ‘inconclusive’ and ‘stable’ sites according to their behavior in a two-generation salinity acclimation experiment with laboratory bred sticklebacks from Kiel (20 PSU) exposed to experimental salinity decrease (6 PSU) (see Methods for details). Further, inducible sites were distinguished whether they matched methylation levels of the locally adapted population (‘*expected*’) or not (‘*opposite*’).

| Ensembl gene ID | chromosome | start position | end position | gene name | wild | inducible | expected inducible | opposite inducible | stable | inconclusive |
|---|---|---|---|---|---|---|---|---|---|---|
| ENSGACG00000008328 | ChrX | 12860144 | 12863850 | si:dkey-166k12.1 | 24 | 0 | 0 | 0 | 9 | 15 |
| ENSGACG00000019416 | ChrVII | 4451892 | 4453656 | HMX1 orthologue | 17 | 0 | 0 | 0 | 9 | 8 |
| ENSGACG00000013229 | ChrXVIII | 15327717 | 15352321 | | 15 | 0 | 0 | 0 | 3 | 12 |
| ENSGACG00000017287 | ChrIII | 13454527 | 13465167 | mmp16b | 12 | 0 | 0 | 0 | 12 | 0 |
| ENSGACG00000017584 | ChrIII | 14690814 | 14694448 | CCNY (1 of many) | 12 | 12 | 12 | 0 | 0 | 0 |
| ENSGACG00000018249 | ChrIV | 12141625 | 12143011 | si:ch211-153b23.5 (1 of many) | 12 | 1 | 1 | 0 | 3 | 8 |
| ENSGACG00000008034 | ChrVI | 9368187 | 9380941 | | 11 | 10 | 10 | 0 | 0 | 1 |
| ENSGACG00000009469 | ChrI | 9166576 | 9173856 | egln2 | 11 | 0 | 0 | 0 | 11 | 0 |
| ENSGACG00000004433 | ChrXVII | 2127457 | 2211376 | igsf21a | 10 | 10 | 10 | 0 | 0 | 0 |
| ENSGACG00000007343 | ChrX | 10666995 | 10679875 | col9a2 | 10 | 0 | 0 | 0 | 6 | 4 |
| ENSGACG00000018407 | ChrIV | 13828336 | 13837518 | sncb | 10 | 2 | 2 | 0 | 5 | 3 |
| ENSGACG00000011118 | ChrI | 12219197 | 12219822 | | 9 | 0 | 0 | 0 | 6 | 3 |
| ENSGACG00000011999 | ChrXIII | 13539291 | 13591502 | pbx3b | 9 | 6 | 0 | 6 | 2 | 1 |
| ENSGACG00000016734 | ChrIV | 3512927 | 3533277 | ANKRD50 | 9 | 0 | 0 | 0 | 9 | 0 |
| ENSGACG00000002963 | ChrXX | 130883 | 136797 | | 8 | 0 | 0 | 0 | 2 | 6 |
| ENSGACG00000014253 | ChrIII | 3157734 | 3197547 | carmil3 | 8 | 0 | 0 | 0 | 3 | 5 |

| | | | | | | | | | | |
|---|---|---|---|---|---|---|---|---|---|---|
| ENSGACG00000015837 | ChrXIV | 1372274 | 1375579 | | 8 | 0 | 0 | 0 | 8 | 0 |
| ENSGACG00000017653 | ChrIX | 8567630 | 8572176 | | 8 | 6 | 0 | 6 | 1 | 1 |
| ENSGACG00000003164 | ChrV | 2512372 | 2514082 | | 7 | 0 | 0 | 0 | 7 | 0 |
| ENSGACG00000016341 | ChrIV | 229120 | 240408 | si:dkeyp-110e4.11 | 7 | 0 | 0 | 0 | 3 | 4 |
| ENSGACG00000006040 | ChrXV | 2368176 | 2420656 | RYR3 (1 of many) | 6 | 0 | 0 | 0 | 1 | 5 |
| ENSGACG00000007557 | ChrXX | 8805057 | 8875965 | | 6 | 0 | 0 | 0 | 3 | 3 |
| ENSGACG00000008805 | ChrX | 13142212 | 13171897 | ST3GAL1 (1 of many) | 6 | 0 | 0 | 0 | 6 | 0 |
| ENSGACG00000013273 | ChrXIII | 16131688 | 16140819 | arhgap25 | 6 | 0 | 0 | 0 | 6 | 0 |
| ENSGACG00000017845 | ChrIX | 8726948 | 8730779 | lgi2b | 6 | 0 | 0 | 0 | 0 | 6 |
| ENSGACG00000018576 | ChrIX | 13131695 | 13177741 | tusc3 | 6 | 0 | 0 | 0 | 6 | 0 |
| ENSGACG00000001150 | scaffold_237 | 44323 | 48391 | | 5 | 0 | 0 | 0 | 2 | 3 |
| ENSGACG00000007077 | ChrXII | 9114925 | 9116176 | cort | 5 | 0 | 0 | 0 | 0 | 5 |
| ENSGACG00000009737 | ChrV | 11530314 | 11534646 | WIPI1 (1 of many) | 5 | 0 | 0 | 0 | 5 | 0 |
| ENSGACG00000010295 | ChrXI | 8105454 | 8118846 | | 5 | 0 | 0 | 0 | 3 | 2 |
| ENSGACG00000011405 | ChrI | 13139185 | 13152967 | stim1a | 5 | 0 | 0 | 0 | 4 | 1 |
| ENSGACG00000014351 | ChrXI | 14886837 | 14931008 | | 5 | 0 | 0 | 0 | 4 | 1 |
| ENSGACG00000017710 | ChrIII | 15790255 | 15791829 | | 5 | 0 | 0 | 0 | 5 | 0 |
| ENSGACG00000018311 | ChrIV | 12800220 | 12810446 | eda | 5 | 3 | 3 | 0 | 0 | 2 |
| ENSGACG00000001679 | ChrXXI | 614128 | 678875 | mpp7a | 4 | 0 | 0 | 0 | 4 | 0 |
| ENSGACG00000003086 | ChrV | 2254186 | 2259462 | trim8b | 4 | 0 | 0 | 0 | 4 | 0 |
| ENSGACG00000004201 | ChrXXI | 9588132 | 9594424 | LPIN2 (1 of many) | 4 | 0 | 0 | 0 | 4 | 0 |
| ENSGACG00000004882 | ChrXXI | 10624323 | 10627058 | | 4 | 2 | 2 | 0 | 1 | 1 |
| ENSGACG00000004909 | ChrXIII | 2375287 | 2400699 | si:dkey-84j12,1 | 4 | 0 | 0 | 0 | 4 | 0 |
| ENSGACG00000005913 | ChrX | 8577242 | 8596962 | elp2 | 4 | 0 | 0 | 0 | 3 | 1 |
| ENSGACG00000006038 | ChrXVII | 4531155 | 4544248 | znfx1 | 4 | 0 | 0 | 0 | 3 | 1 |
| ENSGACG00000006851 | ChrV | 9125837 | 9132564 | | 4 | 4 | 0 | 4 | 0 | 0 |
| ENSGACG00000008558 | ChrVIII | 9695227 | 9702494 | si:ch211-43f4.1 | 4 | 0 | 0 | 0 | 3 | 1 |
| ENSGACG00000010182 | ChrXIII | 9952936 | 9956281 | zgc:110353 | 4 | 0 | 0 | 0 | 4 | 0 |

| | | | | | | | | | | |
|---|---|---|---|---|---|---|---|---|---|---|
| ENSGACG00000010247 | ChrVI | 12459328 | 12462277 | si:ch211-248a14.8 | 4 | 0 | 0 | 0 | 4 | 0 |
| ENSGACG00000013411 | ChrIII | 1381685 | 1420334 | | 4 | 0 | 0 | 0 | 4 | 0 |
| ENSGACG00000013470 | ChrXX | 16251960 | 16257880 | CTSS | 4 | 0 | 0 | 0 | 4 | 0 |
| ENSGACG00000017082 | ChrII | 19781208 | 19783294 | csrp3 | 4 | 0 | 0 | 0 | 3 | 1 |
| ENSGACG00000017422 | ChrXIV | 7780909 | 7787080 | ambp | 4 | 0 | 0 | 0 | 4 | 0 |
| ENSGACG00000018444 | ChrIV | 14030866 | 14039363 | adam19b | 4 | 0 | 0 | 0 | 4 | 0 |
| ENSGACG00000018465 | ChrIX | 12301295 | 12303829 | NOCT (1 of many) | 4 | 0 | 0 | 0 | 1 | 3 |
| ENSGACG00000019155 | ChrIV | 22038773 | 22045261 | mcm10 | 4 | 0 | 0 | 0 | 4 | 0 |
| ENSGACG00000020636 | ChrVII | 21643034 | 21651293 | cfb | 4 | 2 | 2 | 0 | 1 | 1 |
| ENSGACG00000001586 | scaffold_27 | 4643827 | 4660572 | wnk2 | 3 | 0 | 0 | 0 | 1 | 2 |
| ENSGACG00000003140 | ChrX | 3938771 | 3955496 | TGFBR2 | 3 | 0 | 0 | 0 | 2 | 1 |
| ENSGACG00000003570 | ChrXVI | 7947036 | 8001612 | myo16 | 3 | 0 | 0 | 0 | 3 | 0 |
| ENSGACG00000004301 | ChrVIII | 2431974 | 2441330 | brdt | 3 | 0 | 0 | 0 | 3 | 0 |
| ENSGACG00000004534 | ChrX | 6955694 | 6967472 | ef01a | 3 | 0 | 0 | 0 | 3 | 0 |
| ENSGACG00000004850 | ChrXVII | 2873238 | 2885900 | slc2a1b | 3 | 0 | 0 | 0 | 3 | 0 |
| ENSGACG00000006314 | ChrXVI | 12033164 | 12044262 | | 3 | 2 | 2 | 0 | 0 | 1 |
| ENSGACG00000007093 | ChrI | 4607682 | 4609548 | tiparp | 3 | 0 | 0 | 0 | 3 | 0 |
| ENSGACG00000007383 | ChrXII | 9777368 | 9778303 | | 3 | 0 | 0 | 0 | 3 | 0 |
| ENSGACG00000007722 | ChrXVIII | 6627166 | 6639844 | rab15 | 3 | 0 | 0 | 0 | 0 | 3 |
| ENSGACG00000007746 | ChrXV | 4848926 | 4850768 | tmed8 | 3 | 0 | 0 | 0 | 0 | 3 |
| ENSGACG00000009731 | ChrX | 14983618 | 14987412 | mhc1zea | 3 | 0 | 0 | 0 | 3 | 0 |
| ENSGACG00000010537 | ChrXVII | 10768218 | 10769153 | inka1b | 3 | 0 | 0 | 0 | 3 | 0 |
| ENSGACG00000010563 | scaffold_210 | 56747 | 62418 | si:dkey-190l8.2 | 3 | 1 | 0 | 1 | 1 | 1 |
| ENSGACG00000010565 | ChrXVIII | 10469304 | 10475665 | | 3 | 0 | 0 | 0 | 2 | 1 |
| ENSGACG00000011263 | ChrXIII | 12814874 | 12817907 | | 3 | 0 | 0 | 0 | 2 | 1 |
| ENSGACG00000012318 | ChrXVIII | 13638248 | 13642046 | clic5b | 3 | 3 | 3 | 0 | 0 | 0 |
| ENSGACG00000013121 | ChrXVIII | 15160892 | 15161878 | | 3 | 0 | 0 | 0 | 2 | 1 |
| ENSGACG00000013254 | ChrXVIII | 15436938 | 15444996 | tpd52l1 | 3 | 0 | 0 | 0 | 3 | 0 |
| ENSGACG00000014260 | ChrXIII | 18434585 | 18440058 | hspb8 | 3 | 0 | 0 | 0 | 3 | 0 |

| | | | | | | | | | | |
|---|---|---|---|---|---|---|---|---|---|---|
| ENSGACG00000014962 | scaffold_74 | 309652 | 319392 | | 3 | 0 | 0 | 0 | 3 | 0 |
| ENSGACG00000016010 | ChrIII | 9180647 | 9183385 | | 3 | 0 | 0 | 0 | 0 | 3 |
| ENSGACG00000016619 | ChrXIV | 5212300 | 5279453 | rgs3b | 3 | 0 | 0 | 0 | 3 | 0 |
| ENSGACG00000017087 | ChrII | 19884513 | 19943913 | 0v2a | 3 | 0 | 0 | 0 | 3 | 0 |
| ENSGACG00000018250 | ChrIV | 12158019 | 12160272 | irg1l (1 of many) | 3 | 0 | 0 | 0 | 3 | 0 |
| ENSGACG00000018330 | ChrIX | 10996164 | 11001903 | KCNIP2 | 3 | 0 | 0 | 0 | 0 | 3 |
| ENSGACG00000019294 | ChrVII | 3216567 | 3220967 | CCKAR | 3 | 0 | 0 | 0 | 0 | 3 |
| ENSGACG00000019897 | ChrIV | 31023275 | 31026361 | ada2b | 3 | 0 | 0 | 0 | 1 | 2 |
| ENSGACG00000020654 | ChrVII | 21884931 | 21941223 | nrg2a | 3 | 0 | 0 | 0 | 2 | 1 |
| ENSGACG00000022752 | scaffold_90 | 72943 | 73071 | RF00263 | 3 | 0 | 0 | 0 | 3 | 0 |
| ENSGACG00000000086 | scaffold_99 | 339701 | 347704 | brpf1 | 2 | 0 | 0 | 0 | 2 | 0 |
| ENSGACG00000000307 | scaffold_27 | 48405 | 52111 | | 2 | 0 | 0 | 0 | 2 | 0 |
| ENSGACG00000000972 | scaffold_37 | 1733384 | 1743429 | lancl2 | 2 | 0 | 0 | 0 | 2 | 0 |
| ENSGACG00000000969 | scaffold_47 | 1687223 | 1699812 | zcchc4 | 2 | 0 | 0 | 0 | 0 | 2 |
| ENSGACG00000001304 | scaffold_156 | 64555 | 69127 | desi2 | 2 | 0 | 0 | 0 | 1 | 1 |
| ENSGACG00000001438 | ChrXVI | 125393 | 139423 | rrp1 | 2 | 0 | 0 | 0 | 2 | 0 |
| ENSGACG00000002344 | ChrX | 1751605 | 1763088 | nedd9 | 2 | 0 | 0 | 0 | 2 | 0 |
| ENSGACG00000002370 | ChrV | 793362 | 803990 | SPOCK2 | 2 | 0 | 0 | 0 | 2 | 0 |
| ENSGACG00000003413 | ChrXVII | 569558 | 572420 | | 2 | 0 | 0 | 0 | 1 | 1 |
| ENSGACG00000003522 | ChrXVII | 805182 | 813396 | ywhaba | 2 | 0 | 0 | 0 | 2 | 0 |
| ENSGACG00000003564 | ChrXVI | 7915270 | 7918611 | irs2b | 2 | 0 | 0 | 0 | 2 | 0 |
| ENSGACG00000003851 | ChrXVI | 8803555 | 8808879 | gpr143 | 2 | 2 | 1 | 1 | 0 | 0 |
| ENSGACG00000003865 | ChrXXI | 8925778 | 8930097 | insig1 | 2 | 0 | 0 | 0 | 2 | 0 |
| ENSGACG00000004450 | ChrV | 5466348 | 5479796 | ubald1b | 2 | 0 | 0 | 0 | 2 | 0 |
| ENSGACG00000004657 | ChrXVIII | 1241770 | 1243679 | | 2 | 0 | 0 | 0 | 0 | 2 |
| ENSGACG00000004780 | ChrXII | 4743053 | 4896679 | plx01a | 2 | 0 | 0 | 0 | 1 | 1 |
| ENSGACG00000005146 | ChrXXI | 11006245 | 11014630 | gad2 | 2 | 0 | 0 | 0 | 2 | 0 |
| ENSGACG00000005222 | ChrXVII | 3437445 | 3438929 | kc02a | 2 | 1 | 1 | 0 | 0 | 1 |
| ENSGACG00000005316 | ChrXII | 5810740 | 5816656 | fam131c | 2 | 0 | 0 | 0 | 1 | 1 |

| | | | | | | | | | | |
|---|---|---|---|---|---|---|---|---|---|---|
| ENSGACG00000005369 | ChrXV | 1628452 | 1643542 | galnt16 | 2 | 0 | 0 | 0 | 2 | 0 |
| ENSGACG00000005488 | ChrXXI | 11359656 | 11367398 | phex | 2 | 2 | 2 | 0 | 0 | 0 |
| ENSGACG00000005778 | ChrXVI | 11091546 | 11093393 | dap1b | 2 | 1 | 1 | 0 | 1 | 0 |
| ENSGACG00000006075 | ChrXVII | 4589002 | 4596182 | cbfa2t2 | 2 | 0 | 0 | 0 | 2 | 0 |
| ENSGACG00000006148 | ChrI | 2319386 | 2323033 | | 2 | 0 | 0 | 0 | 1 | 1 |
| ENSGACG00000006187 | ChrXVII | 4907492 | 4911313 | timm17a | 2 | 0 | 0 | 0 | 2 | 0 |
| ENSGACG00000006193 | ChrVI | 7611306 | 7689548 | DST | 2 | 0 | 0 | 0 | 2 | 0 |
| ENSGACG00000006321 | ChrI | 3060604 | 3065797 | p2rx8 | 2 | 0 | 0 | 0 | 2 | 0 |
| ENSGACG00000006570 | ChrXVIII | 4669062 | 4680200 | amd1 | 2 | 0 | 0 | 0 | 2 | 0 |
| ENSGACG00000007232 | ChrXV | 4137556 | 4147913 | lck | 2 | 0 | 0 | 0 | 2 | 0 |
| ENSGACG00000007241 | ChrXV | 4150772 | 4153424 | fam167b | 2 | 0 | 0 | 0 | 0 | 2 |
| ENSGACG00000007668 | ChrXX | 8962606 | 8987449 | aplp1 | 2 | 0 | 0 | 0 | 2 | 0 |
| ENSGACG00000007915 | ChrXV | 5068567 | 5089769 | tmem260 | 2 | 1 | 0 | 1 | 0 | 1 |
| ENSGACG00000008836 | ChrXV | 6070712 | 6073477 | klf11b | 2 | 0 | 0 | 0 | 2 | 0 |
| ENSGACG00000008991 | ChrXII | 11144529 | 11145968 | sp5l | 2 | 0 | 0 | 0 | 2 | 0 |
| ENSGACG00000009263 | ChrXII | 11345449 | 11369829 | fmnl3 | 2 | 0 | 0 | 0 | 2 | 0 |
| ENSGACG00000009558 | ChrXI | 6851020 | 6922054 | asic2 | 2 | 2 | 2 | 0 | 0 | 0 |
| ENSGACG00000009925 | ChrXVIII | 9577490 | 9600773 | lpgat1 | 2 | 0 | 0 | 0 | 2 | 0 |
| ENSGACG00000010040 | ChrVIII | 12761712 | 12774504 | ccdc50 | 2 | 0 | 0 | 0 | 0 | 2 |
| ENSGACG00000010381 | ChrX | 15527868 | 15528362 | | 2 | 0 | 0 | 0 | 0 | 2 |
| ENSGACG00000010475 | ChrVIII | 13174745 | 13231603 | insrb | 2 | 0 | 0 | 0 | 2 | 0 |
| ENSGACG00000010791 | ChrXVIII | 10774342 | 10785642 | fndc4a | 2 | 1 | 1 | 0 | 1 | 0 |
| ENSGACG00000011302 | ChrXII | 15080122 | 15082735 | mul1b | 2 | 0 | 0 | 0 | 2 | 0 |
| ENSGACG00000011602 | ChrXVII | 12652011 | 12662065 | LAMB3 | 2 | 0 | 0 | 0 | 2 | 0 |
| ENSGACG00000011732 | ChrXVIII | 12338480 | 12366063 | myo6a | 2 | 0 | 0 | 0 | 2 | 0 |
| ENSGACG00000011811 | ChrXVIII | 12406545 | 12408354 | COX7A2 | 2 | 0 | 0 | 0 | 2 | 0 |
| ENSGACG00000012139 | ChrI | 14774869 | 14796093 | yap1 | 2 | 0 | 0 | 0 | 2 | 0 |
| ENSGACG00000012773 | ChrXV | 13276935 | 13293594 | pdss2 | 2 | 1 | 1 | 0 | 0 | 1 |
| ENSGACG00000013429 | ChrXI | 13025988 | 13026614 | | 2 | 1 | 0 | 1 | 1 | 0 |

| | | | | | | | | | | |
|---|---|---|---|---|---|---|---|---|---|---|
| ENSGACG00000013428 | ChrVIII | 17395611 | 17400555 | trmt13 | 2 | 0 | 0 | 0 | 2 | 0 |
| ENSGACG00000013595 | ChrXIII | 16645839 | 16651528 | vamp8 | 2 | 0 | 0 | 0 | 2 | 0 |
| ENSGACG00000013744 | ChrVIII | 17762827 | 17768426 | | 2 | 0 | 0 | 0 | 2 | 0 |
| ENSGACG00000013743 | ChrIII | 1852860 | 1900173 | tnr | 2 | 0 | 0 | 0 | 2 | 0 |
| ENSGACG00000014053 | ChrI | 20793942 | 20794313 | nhlh2 | 2 | 0 | 0 | 0 | 2 | 0 |
| ENSGACG00000014454 | ChrXI | 15389422 | 15396323 | elac2 | 2 | 0 | 0 | 0 | 1 | 1 |
| ENSGACG00000014902 | ChrII | 5427953 | 5521857 | itfg1 | 2 | 0 | 0 | 0 | 1 | 1 |
| ENSGACG00000016477 | ChrXIV | 3999374 | 4006176 | agxt2 (1 of many) | 2 | 0 | 0 | 0 | 2 | 0 |
| ENSGACG00000016694 | ChrII | 17411270 | 17445836 | chd9 | 2 | 0 | 0 | 0 | 1 | 1 |
| ENSGACG00000017537 | ChrXIV | 8099548 | 8156144 | bcr | 2 | 0 | 0 | 0 | 2 | 0 |
| ENSGACG00000018722 | scaffold_208 | 83156 | 90702 | iffo1a | 2 | 0 | 0 | 0 | 2 | 0 |
| ENSGACG00000019481 | ChrIV | 25424476 | 25433284 | krr1 | 2 | 1 | 1 | 0 | 1 | 0 |
| ENSGACG00000020034 | ChrIV | 31999298 | 32000748 | tnnt2c | 2 | 0 | 0 | 0 | 1 | 1 |
| ENSGACG00000020040 | ChrIV | 32012155 | 32084033 | btbd11a | 2 | 0 | 0 | 0 | 2 | 0 |
| ENSGACG00000020073 | ChrIV | 32350582 | 32355303 | dclre1c | 2 | 0 | 0 | 0 | 2 | 0 |
| ENSGACG00000020706 | ChrVII | 22666912 | 22672421 | | 2 | 0 | 0 | 0 | 1 | 1 |
| ENSGACG00000020808 | ChrVII | 25496687 | 25519233 | grk6 | 2 | 0 | 0 | 0 | 2 | 0 |
| ENSGACG00000022198 | ChrII | 15100637 | 15100747 | | 2 | 0 | 0 | 0 | 2 | 0 |
| ENSGACG00000000048 | scaffold_757 | 6140 | 6811 | | 1 | 0 | 0 | 0 | 1 | 0 |
| ENSGACG00000000162 | scaffold_216 | 55071 | 61253 | ACTR2 | 1 | 0 | 0 | 0 | 0 | 1 |
| ENSGACG00000000381 | scaffold_131 | 232303 | 234144 | rbm46 | 1 | 1 | 1 | 0 | 0 | 0 |
| ENSGACG00000000429 | scaffold_80 | 140050 | 141102 | | 1 | 0 | 0 | 0 | 1 | 0 |
| ENSGACG00000000434 | scaffold_80 | 184463 | 204053 | | 1 | 0 | 0 | 0 | 1 | 0 |
| ENSGACG00000000504 | scaffold_139 | 130345 | 136698 | DRC7 | 1 | 0 | 0 | 0 | 1 | 0 |
| ENSGACG00000000525 | scaffold_397 | 1558 | 15029 | | 1 | 0 | 0 | 0 | 1 | 0 |
| ENSGACG00000000634 | scaffold_90 | 226420 | 229275 | cradd | 1 | 1 | 0 | 1 | 0 | 0 |
| ENSGACG00000000795 | scaffold_141 | 12636 | 18456 | fgd1 | 1 | 0 | 0 | 0 | 0 | 1 |
| ENSGACG00000000935 | scaffold_1182 | 4446 | 4961 | | 1 | 1 | 0 | 1 | 0 | 0 |
| ENSGACG00000000950 | scaffold_157 | 146047 | 149073 | si:dkey-201i6.2 | 1 | 0 | 0 | 0 | 1 | 0 |

| | | | | | | | | | | |
|---|---|---|---|---|---|---|---|---|---|---|
| ENSGACG00000001006 | scaffold_37 | 2177494 | 2216214 | arhgap12b | 1 | 0 | 0 | 0 | 1 | 0 |
| ENSGACG00000001021 | scaffold_37 | 2247366 | 2268896 | si:ch211-276f18.2 | 1 | 0 | 0 | 0 | 1 | 0 |
| ENSGACG00000001050 | scaffold_37 | 2338551 | 2342806 | aoc1 | 1 | 0 | 0 | 0 | 1 | 0 |
| ENSGACG00000001087 | scaffold_150 | 77433 | 80873 | | 1 | 0 | 0 | 0 | 1 | 0 |
| ENSGACG00000001219 | scaffold_27 | 3491387 | 3517587 | skib | 1 | 0 | 0 | 0 | 1 | 0 |
| ENSGACG00000001233 | scaffold_181 | 61484 | 71960 | adcy5 | 1 | 0 | 0 | 0 | 1 | 0 |
| ENSGACG00000001282 | scaffold_249 | 6147 | 18792 | EHD4 | 1 | 0 | 0 | 0 | 1 | 0 |
| ENSGACG00000001284 | scaffold_130 | 39785 | 63580 | sema4ab | 1 | 0 | 0 | 0 | 1 | 0 |
| ENSGACG00000001458 | scaffold_273 | 15193 | 18488 | | 1 | 0 | 0 | 0 | 1 | 0 |
| ENSGACG00000001467 | scaffold_273 | 23860 | 30175 | sec24a | 1 | 1 | 1 | 0 | 0 | 0 |
| ENSGACG00000001644 | scaffold_27 | 4804545 | 4808600 | tmcc1b | 1 | 1 | 1 | 0 | 0 | 0 |
| ENSGACG00000001842 | scaffold_600 | 385 | 1591 | | 1 | 0 | 0 | 0 | 0 | 1 |
| ENSGACG00000001883 | ChrXVI | 2628609 | 2645144 | klf12a | 1 | 0 | 0 | 0 | 1 | 0 |
| ENSGACG00000001886 | ChrXVI | 2686951 | 2764612 | dachd | 1 | 0 | 0 | 0 | 1 | 0 |
| ENSGACG00000001897 | scaffold_326 | 46372 | 48199 | mrps24 | 1 | 1 | 0 | 1 | 0 | 0 |
| ENSGACG00000001979 | ChrX | 1148457 | 1148636 | | 1 | 0 | 0 | 0 | 1 | 0 |
| ENSGACG00000001980 | scaffold_149 | 22071 | 23010 | | 1 | 0 | 0 | 0 | 0 | 1 |
| ENSGACG00000002013 | scaffold_56 | 24216 | 25922 | | 1 | 0 | 0 | 0 | 1 | 0 |
| ENSGACG00000002128 | ChrXXI | 2737163 | 2755917 | cngb3.1 | 1 | 0 | 0 | 0 | 1 | 0 |
| ENSGACG00000002216 | ChrXVI | 4428375 | 4457157 | ADARB1 | 1 | 0 | 0 | 0 | 1 | 0 |
| ENSGACG00000002270 | ChrX | 1627559 | 1629451 | tekt1 | 1 | 0 | 0 | 0 | 0 | 1 |
| ENSGACG00000002326 | ChrXXI | 3947748 | 3952816 | slc35b3 | 1 | 0 | 0 | 0 | 1 | 0 |
| ENSGACG00000002374 | ChrXXI | 4343544 | 4350834 | bco1 | 1 | 0 | 0 | 0 | 0 | 1 |
| ENSGACG00000002418 | ChrXXI | 4917563 | 4971561 | zfhx4 | 1 | 0 | 0 | 0 | 1 | 0 |
| ENSGACG00000002439 | ChrVI | 624617 | 631220 | degs1 | 1 | 0 | 0 | 0 | 1 | 0 |
| ENSGACG00000002469 | ChrXXI | 5440575 | 5443026 | | 1 | 0 | 0 | 0 | 0 | 1 |
| ENSGACG00000002501 | ChrXVI | 5863099 | 5885871 | slc4a3 | 1 | 0 | 0 | 0 | 1 | 0 |
| ENSGACG00000002545 | ChrX | 2269766 | 2289402 | nfatc1 | 1 | 0 | 0 | 0 | 1 | 0 |
| ENSGACG00000002663 | ChrV | 1626313 | 1664933 | ptprea | 1 | 0 | 0 | 0 | 1 | 0 |

| | | | | | | | | | | |
|---|---|---|---|---|---|---|---|---|---|---|
| ENSGACG00000002670 | ChrXXI | 6065576 | 6128487 | sema5a | 1 | 0 | 0 | 0 | 0 | 1 |
| ENSGACG00000002757 | ChrXXI | 6579026 | 6642145 | stau2 | 1 | 0 | 0 | 0 | 0 | 1 |
| ENSGACG00000002788 | ChrXXI | 6671169 | 6674187 | SBSPON | 1 | 0 | 0 | 0 | 1 | 0 |
| ENSGACG00000002861 | ChrV | 1990736 | 1995983 | | 1 | 0 | 0 | 0 | 0 | 1 |
| ENSGACG00000002882 | ChrXVI | 7062666 | 7077323 | ATP11A (1 of many) | 1 | 0 | 0 | 0 | 1 | 0 |
| ENSGACG00000002912 | ChrXVI | 7222380 | 7225324 | txndc9 | 1 | 0 | 0 | 0 | 1 | 0 |
| ENSGACG00000002982 | ChrXII | 1118299 | 1128392 | TMCC1 (1 of many) | 1 | 0 | 0 | 0 | 0 | 1 |
| ENSGACG00000003032 | ChrVI | 2464760 | 2466700 | fignl1 | 1 | 0 | 0 | 0 | 0 | 1 |
| ENSGACG00000003031 | ChrXVI | 7391481 | 7394255 | ecrg4a | 1 | 0 | 0 | 0 | 1 | 0 |
| ENSGACG00000003212 | ChrVI | 2759500 | 2773283 | fam204a | 1 | 0 | 0 | 0 | 1 | 0 |
| ENSGACG00000003235 | ChrXX | 629982 | 640598 | tmem145 | 1 | 0 | 0 | 0 | 1 | 0 |
| ENSGACG00000003244 | ChrX | 4431116 | 4431815 | | 1 | 1 | 0 | 1 | 0 | 0 |
| ENSGACG00000003270 | ChrV | 2667010 | 2669113 | si:ch73-127m5.2 | 1 | 0 | 0 | 0 | 1 | 0 |
| ENSGACG00000003327 | ChrX | 4521465 | 4540223 | ash1l | 1 | 0 | 0 | 0 | 1 | 0 |
| ENSGACG00000003323 | ChrXII | 1813168 | 1898170 | | 1 | 0 | 0 | 0 | 1 | 0 |
| ENSGACG00000003383 | ChrVIII | 1540683 | 1540880 | | 1 | 0 | 0 | 0 | 1 | 0 |
| ENSGACG00000003395 | ChrXVI | 7711922 | 7714172 | asb11 | 1 | 0 | 0 | 0 | 1 | 0 |
| ENSGACG00000003425 | ChrXII | 2053482 | 2071894 | mitfb | 1 | 0 | 0 | 0 | 1 | 0 |
| ENSGACG00000003495 | ChrXVII | 699787 | 719603 | GRIP2 | 1 | 0 | 0 | 0 | 1 | 0 |
| ENSGACG00000003515 | ChrXXI | 8073271 | 8088883 | | 1 | 0 | 0 | 0 | 0 | 1 |
| ENSGACG00000003552 | ChrX | 5093819 | 5117753 | pard6gb | 1 | 0 | 0 | 0 | 1 | 0 |
| ENSGACG00000003582 | ChrXVI | 8023067 | 8024372 | tnfsf13b | 1 | 0 | 0 | 0 | 1 | 0 |
| ENSGACG00000003600 | ChrXXI | 8136415 | 8143194 | | 1 | 1 | 1 | 0 | 0 | 0 |
| ENSGACG00000003640 | ChrXVII | 1072273 | 1073132 | | 1 | 0 | 0 | 0 | 1 | 0 |
| ENSGACG00000003722 | ChrV | 4090150 | 4096264 | meox1 | 1 | 0 | 0 | 0 | 1 | 0 |
| ENSGACG00000003781 | ChrXX | 1859763 | 1890241 | | 1 | 0 | 0 | 0 | 1 | 0 |
| ENSGACG00000003807 | ChrV | 4374700 | 4425195 | si:ch211-94l19.4 | 1 | 0 | 0 | 0 | 1 | 0 |
| ENSGACG00000003932 | ChrX | 5372498 | 5374253 | | 1 | 0 | 0 | 0 | 1 | 0 |

| | | | | | | | | | | |
|---|---|---|---|---|---|---|---|---|---|---|
| ENSGACG00000003937 | ChrX | 5419651 | 5422992 | tshz1 | 1 | 1 | 1 | 0 | 0 | 0 |
| ENSGACG00000003958 | ChrVI | 4091981 | 4093294 | sertad2b | 1 | 0 | 0 | 0 | 1 | 0 |
| ENSGACG00000003974 | ChrVI | 4172491 | 4201696 | b3gat2 | 1 | 0 | 0 | 0 | 1 | 0 |
| ENSGACG00000004014 | ChrXVI | 8929582 | 8950765 | tlr7 | 1 | 0 | 0 | 0 | 0 | 1 |
| ENSGACG00000004100 | ChrXVIII | 313994 | 349188 | e0h | 1 | 0 | 0 | 0 | 0 | 1 |
| ENSGACG00000004101 | ChrXVI | 9224446 | 9228563 | dct | 1 | 0 | 0 | 0 | 0 | 1 |
| ENSGACG00000004118 | ChrVI | 4433273 | 4465659 | | 1 | 0 | 0 | 0 | 0 | 1 |
| ENSGACG00000004149 | ChrVI | 4667811 | 4669850 | ntpcr | 1 | 0 | 0 | 0 | 1 | 0 |
| ENSGACG00000004339 | ChrVIII | 2537139 | 2541472 | | 1 | 1 | 1 | 0 | 0 | 0 |
| ENSGACG00000004294 | ChrXX | 2209827 | 2216716 | sacm1lb | 1 | 1 | 1 | 0 | 0 | 0 |
| ENSGACG00000004352 | ChrXVII | 1895568 | 1903148 | | 1 | 1 | 1 | 0 | 0 | 0 |
| ENSGACG00000004352 | ChrXVII | 1895568 | 1903148 | | 1 | 0 | 0 | 0 | 1 | 0 |
| ENSGACG00000004383 | ChrVIII | 2891959 | 2898703 | lmo4b | 1 | 0 | 0 | 0 | 1 | 0 |
| ENSGACG00000004402 | ChrV | 5177690 | 5178772 | ccr10 | 1 | 0 | 0 | 0 | 1 | 0 |
| ENSGACG00000004412 | ChrXVII | 1957372 | 2014039 | arhgef10la | 1 | 0 | 0 | 0 | 1 | 0 |
| ENSGACG00000004431 | ChrXII | 3958137 | 3978532 | ntsr1 | 1 | 1 | 0 | 1 | 0 | 0 |
| ENSGACG00000004474 | ChrV | 5578449 | 5580362 | | 1 | 0 | 0 | 0 | 1 | 0 |
| ENSGACG00000004508 | ChrXVIII | 1155689 | 1171739 | emilin1a | 1 | 0 | 0 | 0 | 1 | 0 |
| ENSGACG00000004559 | ChrV | 5619682 | 5624540 | tmem184ba | 1 | 0 | 0 | 0 | 1 | 0 |
| ENSGACG00000004636 | ChrI | 372834 | 374001 | | 1 | 1 | 1 | 0 | 0 | 0 |
| ENSGACG00000004738 | ChrXVII | 2595419 | 2644876 | | 1 | 0 | 0 | 0 | 1 | 0 |
| ENSGACG00000004770 | ChrXVI | 10045160 | 10049909 | SP3 | 1 | 0 | 0 | 0 | 1 | 0 |
| ENSGACG00000004870 | ChrXXI | 10440079 | 10446151 | dsela | 1 | 1 | 1 | 0 | 0 | 0 |
| ENSGACG00000004873 | ChrXXI | 10500632 | 10511464 | CDH19 | 1 | 1 | 1 | 0 | 0 | 0 |
| ENSGACG00000004924 | ChrXII | 5500786 | 5508872 | brpf3b | 1 | 0 | 0 | 0 | 1 | 0 |
| ENSGACG00000004937 | ChrV | 5987158 | 5991150 | zdhhc16b | 1 | 0 | 0 | 0 | 1 | 0 |
| ENSGACG00000004954 | ChrXVII | 3053327 | 3129303 | grm4 | 1 | 0 | 0 | 0 | 1 | 0 |
| ENSGACG00000005016 | ChrVIII | 3619176 | 3623161 | | 1 | 0 | 0 | 0 | 1 | 0 |
| ENSGACG00000005087 | ChrXXI | 10930916 | 10935699 | mastl | 1 | 0 | 0 | 0 | 0 | 1 |

| | | | | | | | | | | |
|---|---|---|---|---|---|---|---|---|---|---|
| ENSGACG00000005081 | ChrXV | 1425353 | 1432246 | | 1 | 0 | 0 | 0 | 0 | 1 |
| ENSGACG00000005129 | ChrVIII | 3952798 | 3985032 | si:ch211-242b18.1 | 1 | 0 | 0 | 0 | 0 | 1 |
| ENSGACG00000005157 | ChrXVI | 10427406 | 10429718 | ccdc173 | 1 | 0 | 0 | 0 | 0 | 1 |
| ENSGACG00000005173 | ChrI | 931955 | 937054 | hmbsa | 1 | 0 | 0 | 0 | 0 | 1 |
| ENSGACG00000005197 | ChrX | 7842276 | 7843919 | ggcta | 1 | 0 | 0 | 0 | 0 | 1 |
| ENSGACG00000005264 | ChrXVI | 10507462 | 10514195 | rdh1 | 1 | 0 | 0 | 0 | 1 | 0 |
| ENSGACG00000005270 | ChrXI | 1017948 | 1036084 | | 1 | 0 | 0 | 0 | 1 | 0 |
| ENSGACG00000005280 | ChrXVIII | 2211718 | 2416099 | nrxn3b | 1 | 0 | 0 | 0 | 1 | 0 |
| ENSGACG00000005319 | ChrXIII | 3168194 | 3174770 | | 1 | 0 | 0 | 0 | 1 | 0 |
| ENSGACG00000005284 | ChrXVII | 3503858 | 3508516 | slc5a8l | 1 | 0 | 0 | 0 | 1 | 0 |
| ENSGACG00000005332 | ChrVIII | 4566579 | 4571023 | atpaf1 | 1 | 0 | 0 | 0 | 1 | 0 |
| ENSGACG00000005353 | ChrVIII | 4667337 | 4693951 | | 1 | 0 | 0 | 0 | 1 | 0 |
| ENSGACG00000005493 | ChrXV | 1689391 | 1695853 | metap1 | 1 | 1 | 1 | 0 | 0 | 0 |
| ENSGACG00000005584 | ChrI | 1585298 | 1585992 | | 1 | 0 | 0 | 0 | 1 | 0 |
| ENSGACG00000005603 | ChrXX | 4857164 | 4860534 | has2 | 1 | 0 | 0 | 0 | 1 | 0 |
| ENSGACG00000005647 | ChrXV | 1810555 | 1848494 | sntg2 | 1 | 0 | 0 | 0 | 1 | 0 |
| ENSGACG00000005688 | ChrVIII | 5057122 | 5059646 | | 1 | 0 | 0 | 0 | 1 | 0 |
| ENSGACG00000005678 | ChrXVII | 3951822 | 3964914 | cers5 | 1 | 0 | 0 | 0 | 0 | 1 |
| ENSGACG00000005740 | ChrV | 7636119 | 7639426 | | 1 | 0 | 0 | 0 | 0 | 1 |
| ENSGACG00000005765 | ChrXVII | 3986584 | 4098651 | si:dkey-57c15.1 | 1 | 0 | 0 | 0 | 1 | 0 |
| ENSGACG00000005753 | ChrXII | 6969334 | 7035500 | celsr3 | 1 | 0 | 0 | 0 | 0 | 1 |
| ENSGACG00000005814 | ChrXV | 2022468 | 2079686 | LTBP2 | 1 | 0 | 0 | 0 | 1 | 0 |
| ENSGACG00000005851 | ChrXVI | 11375726 | 11391780 | kcnj3a | 1 | 0 | 0 | 0 | 1 | 0 |
| ENSGACG00000005869 | ChrI | 2075764 | 2085083 | numbl | 1 | 1 | 1 | 0 | 0 | 0 |
| ENSGACG00000005912 | ChrXVII | 4162499 | 4184565 | eya2 | 1 | 0 | 0 | 0 | 1 | 0 |
| ENSGACG00000005959 | ChrXIII | 4441375 | 4446248 | CKMT2 | 1 | 0 | 0 | 0 | 0 | 1 |
| ENSGACG00000006022 | ChrI | 2206772 | 2213401 | bace1 | 1 | 0 | 0 | 0 | 1 | 0 |
| ENSGACG00000006044 | ChrVIII | 6015397 | 6017317 | | 1 | 0 | 0 | 0 | 1 | 0 |
| ENSGACG00000006136 | ChrI | 2281697 | 2283639 | | 1 | 0 | 0 | 0 | 1 | 0 |

| | | | | | | | | | | |
|---|---|---|---|---|---|---|---|---|---|---|
| ENSGACG00000006168 | ChrXVIII | 3331871 | 3333721 | col10a1b | 1 | 1 | 1 | 0 | 0 | 0 |
| ENSGACG00000006183 | ChrXV | 2631748 | 2632214 | gchfr | 1 | 0 | 0 | 0 | 1 | 0 |
| ENSGACG00000006180 | ChrVIII | 6394455 | 6397463 | | 1 | 1 | 1 | 0 | 0 | 0 |
| ENSGACG00000006245 | ChrXVIII | 3473616 | 3494661 | bach2b | 1 | 0 | 0 | 0 | 0 | 1 |
| ENSGACG00000006309 | ChrXV | 2723196 | 2735169 | cep170b | 1 | 0 | 0 | 0 | 1 | 0 |
| ENSGACG00000006345 | ChrXX | 6003650 | 6006502 | eomesb | 1 | 0 | 0 | 0 | 1 | 0 |
| ENSGACG00000006380 | ChrXIII | 5034648 | 5038273 | | 1 | 0 | 0 | 0 | 0 | 1 |
| ENSGACG00000006398 | ChrXV | 2806798 | 2812667 | pak6a | 1 | 0 | 0 | 0 | 1 | 0 |
| ENSGACG00000006421 | ChrX | 9120079 | 9120495 | ferd3l | 1 | 0 | 0 | 0 | 1 | 0 |
| ENSGACG00000006443 | ChrXII | 7864313 | 7874314 | ppih | 1 | 0 | 0 | 0 | 1 | 0 |
| ENSGACG00000006458 | ChrXVII | 5292055 | 5300667 | gli1 | 1 | 0 | 0 | 0 | 1 | 0 |
| ENSGACG00000006465 | ChrXVI | 12261048 | 12264385 | pofut2 | 1 | 0 | 0 | 0 | 0 | 1 |
| ENSGACG00000006528 | ChrXVI | 12761910 | 12797556 | cps1 | 1 | 1 | 1 | 0 | 0 | 0 |
| ENSGACG00000006533 | ChrXV | 2891400 | 2894701 | | 1 | 0 | 0 | 0 | 1 | 0 |
| ENSGACG00000006537 | ChrVI | 7898542 | 7919293 | mcph1 | 1 | 0 | 0 | 0 | 1 | 0 |
| ENSGACG00000006563 | ChrV | 8413714 | 8419499 | dclre1a | 1 | 0 | 0 | 0 | 1 | 0 |
| ENSGACG00000006591 | ChrI | 3486252 | 3490154 | | 1 | 0 | 0 | 0 | 1 | 0 |
| ENSGACG00000006594 | ChrXVI | 12865337 | 12867816 | | 1 | 0 | 0 | 0 | 0 | 1 |
| ENSGACG00000006597 | ChrXVI | 12870903 | 12877502 | | 1 | 0 | 0 | 0 | 1 | 0 |
| ENSGACG00000006604 | ChrVIII | 6966044 | 6979032 | WDR63 | 1 | 0 | 0 | 0 | 1 | 0 |
| ENSGACG00000006606 | ChrXI | 3172985 | 3193999 | cpt1cb | 1 | 0 | 0 | 0 | 1 | 0 |
| ENSGACG00000006693 | ChrXV | 2973149 | 2977521 | | 1 | 0 | 0 | 0 | 1 | 0 |
| ENSGACG00000006736 | ChrXII | 8388213 | 8475879 | KAZN (1 of many) | 1 | 0 | 0 | 0 | 0 | 1 |
| ENSGACG00000006837 | ChrV | 9060530 | 9065297 | nmt1b | 1 | 0 | 0 | 0 | 1 | 0 |
| ENSGACG00000006800 | ChrXV | 3469650 | 3516066 | col12a1a | 1 | 0 | 0 | 0 | 1 | 0 |
| ENSGACG00000006884 | ChrXV | 3547262 | 3568975 | tfb1m | 1 | 0 | 0 | 0 | 1 | 0 |
| ENSGACG00000006896 | ChrXI | 3652039 | 3673572 | grb2b | 1 | 1 | 1 | 0 | 0 | 0 |
| ENSGACG00000006931 | ChrXV | 3674888 | 3699659 | | 1 | 0 | 0 | 0 | 1 | 0 |
| ENSGACG00000006980 | ChrXV | 3714812 | 3741586 | | 1 | 1 | 1 | 0 | 0 | 0 |

| | | | | | | | | | | |
|---|---|---|---|---|---|---|---|---|---|---|
| ENSGACG00000007218 | ChrXIII | 6430504 | 6435884 | plk2b | 1 | 0 | 0 | 0 | 1 | 0 |
| ENSGACG00000007276 | ChrI | 5465852 | 5488747 | mecom | 1 | 0 | 0 | 0 | 0 | 1 |
| ENSGACG00000007280 | ChrXV | 4171868 | 4184504 | clic4 | 1 | 1 | 1 | 0 | 0 | 0 |
| ENSGACG00000007293 | ChrX | 10477197 | 10572975 | adgrb2 | 1 | 0 | 0 | 0 | 1 | 0 |
| ENSGACG00000007320 | ChrI | 5512892 | 5515273 | | 1 | 0 | 0 | 0 | 1 | 0 |
| ENSGACG00000007342 | ChrXIII | 6570975 | 6572197 | atoh1a | 1 | 0 | 0 | 0 | 1 | 0 |
| ENSGACG00000007372 | ChrX | 10807926 | 10961030 | csmd2 | 1 | 1 | 0 | 1 | 0 | 0 |
| ENSGACG00000007372 | ChrX | 10807926 | 10961030 | csmd2 | 1 | 0 | 0 | 0 | 1 | 0 |
| ENSGACG00000007422 | ChrXX | 8591407 | 8593694 | prss1 (1 of many) | 1 | 0 | 0 | 0 | 0 | 1 |
| ENSGACG00000007449 | ChrXI | 4378241 | 4381177 | | 1 | 1 | 1 | 0 | 0 | 0 |
| ENSGACG00000007492 | ChrX | 11267031 | 11270381 | heyl | 1 | 0 | 0 | 0 | 1 | 0 |
| ENSGACG00000007529 | ChrVI | 8795607 | 8802574 | CNNM1 (1 of many) | 1 | 0 | 0 | 0 | 1 | 0 |
| ENSGACG00000007503 | ChrXVI | 14554631 | 14687685 | lrp1bb | 1 | 0 | 0 | 0 | 1 | 0 |
| ENSGACG00000007578 | ChrXVI | 14827472 | 14849263 | myo1b | 1 | 0 | 0 | 0 | 1 | 0 |
| ENSGACG00000007610 | ChrX | 11394452 | 11398811 | slc6a18 | 1 | 0 | 0 | 0 | 0 | 1 |
| ENSGACG00000007653 | ChrXIII | 7397696 | 7405305 | 0a35 | 1 | 0 | 0 | 0 | 1 | 0 |
| ENSGACG00000007695 | ChrI | 5944178 | 5955498 | porb | 1 | 0 | 0 | 0 | 0 | 1 |
| ENSGACG00000007724 | ChrXI | 4748362 | 4752125 | cbx7a | 1 | 0 | 0 | 0 | 1 | 0 |
| ENSGACG00000007745 | ChrXIII | 7495399 | 7509368 | EGFLAM | 1 | 1 | 1 | 0 | 0 | 0 |
| ENSGACG00000007773 | ChrX | 11796739 | 11880195 | ptprua | 1 | 0 | 0 | 0 | 1 | 0 |
| ENSGACG00000007826 | ChrX | 12126255 | 12133705 | lin28a | 1 | 0 | 0 | 0 | 1 | 0 |
| ENSGACG00000007825 | ChrI | 6236909 | 6237643 | a0pc15 | 1 | 0 | 0 | 0 | 1 | 0 |
| ENSGACG00000007869 | ChrXV | 4989079 | 4992230 | ap5m1 | 1 | 0 | 0 | 0 | 1 | 0 |
| ENSGACG00000007893 | ChrXX | 9249276 | 9251026 | | 1 | 1 | 0 | 1 | 0 | 0 |
| ENSGACG00000007889 | ChrXVI | 15735794 | 15745166 | mpp4a | 1 | 1 | 1 | 0 | 0 | 0 |
| ENSGACG00000008000 | ChrXIII | 8084061 | 8086494 | rfk | 1 | 0 | 0 | 0 | 1 | 0 |
| ENSGACG00000008041 | ChrVIII | 9282717 | 9296560 | | 1 | 0 | 0 | 0 | 0 | 1 |
| ENSGACG00000008073 | ChrV | 9942067 | 9963447 | kcnc3a | 1 | 0 | 0 | 0 | 1 | 0 |

| | | | | | | | | | | |
|---|---|---|---|---|---|---|---|---|---|---|
| ENSGACG00000008081 | ChrV | 9974960 | 9977703 | | 1 | 0 | 0 | 0 | 1 | 0 |
| ENSGACG00000008079 | ChrXII | 10283054 | 10285128 | dclre1b | 1 | 0 | 0 | 0 | 1 | 0 |
| ENSGACG00000008092 | ChrX | 12446407 | 12460395 | eepd1 | 1 | 0 | 0 | 0 | 1 | 0 |
| ENSGACG00000008121 | ChrI | 6554830 | 6569989 | pdgfd | 1 | 0 | 0 | 0 | 0 | 1 |
| ENSGACG00000008082 | ChrXII | 10316175 | 10365141 | cpne5a | 1 | 0 | 0 | 0 | 0 | 1 |
| ENSGACG00000008174 | ChrX | 12578667 | 12611207 | TRIO | 1 | 0 | 0 | 0 | 1 | 0 |
| ENSGACG00000008194 | ChrVIII | 9448685 | 9451926 | si:dkey-197i20.6 | 1 | 0 | 0 | 0 | 1 | 0 |
| ENSGACG00000008187 | ChrV | 10034443 | 10040862 | | 1 | 0 | 0 | 0 | 1 | 0 |
| ENSGACG00000008213 | ChrV | 10078945 | 10079847 | | 1 | 0 | 0 | 0 | 0 | 1 |
| ENSGACG00000008282 | ChrXV | 5566137 | 5574768 | FGFRL1 (1 of many) | 1 | 0 | 0 | 0 | 1 | 0 |
| ENSGACG00000008305 | ChrXVI | 16460421 | 16461727 | | 1 | 0 | 0 | 0 | 1 | 0 |
| ENSGACG00000008509 | ChrXX | 9883924 | 9885147 | NHLRC1 | 1 | 0 | 0 | 0 | 0 | 1 |
| ENSGACG00000008542 | ChrXV | 5811952 | 5817096 | tpp1 | 1 | 0 | 0 | 0 | 1 | 0 |
| ENSGACG00000008613 | ChrXV | 5827640 | 5855519 | evla | 1 | 0 | 0 | 0 | 1 | 0 |
| ENSGACG00000008642 | ChrXVI | 17305099 | 17306341 | | 1 | 0 | 0 | 0 | 1 | 0 |
| ENSGACG00000008710 | ChrXVI | 17405163 | 17405973 | | 1 | 0 | 0 | 0 | 1 | 0 |
| ENSGACG00000008728 | ChrV | 10795397 | 10797367 | znf668 | 1 | 0 | 0 | 0 | 0 | 1 |
| ENSGACG00000008764 | ChrVI | 9973371 | 9977148 | ppifa | 1 | 0 | 0 | 0 | 0 | 1 |
| ENSGACG00000008862 | ChrX | 13440280 | 13453314 | | 1 | 0 | 0 | 0 | 0 | 1 |
| ENSGACG00000008937 | ChrXI | 6189394 | 6191153 | zgc:91968 | 1 | 0 | 0 | 0 | 0 | 1 |
| ENSGACG00000009012 | ChrX | 13753836 | 13755884 | | 1 | 0 | 0 | 0 | 1 | 0 |
| ENSGACG00000009025 | ChrV | 10893482 | 10895023 | zgc:112496 | 1 | 0 | 0 | 0 | 1 | 0 |
| ENSGACG00000009033 | ChrV | 10926848 | 10927459 | h1f0 | 1 | 0 | 0 | 0 | 0 | 1 |
| ENSGACG00000009037 | ChrXVI | 17622563 | 17645255 | satb2 | 1 | 0 | 0 | 0 | 0 | 1 |
| ENSGACG00000009043 | ChrVI | 10538527 | 10559377 | pcsk2 | 1 | 0 | 0 | 0 | 0 | 1 |
| ENSGACG00000009105 | ChrV | 10961013 | 10969417 | nlrc3 | 1 | 0 | 0 | 0 | 1 | 0 |
| ENSGACG00000009086 | ChrVIII | 11312464 | 11328558 | rubcn | 1 | 0 | 0 | 0 | 1 | 0 |
| ENSGACG00000009135 | ChrVI | 10670867 | 10727803 | pcdh15a | 1 | 0 | 0 | 0 | 0 | 1 |

| | | | | | | | | | | |
|---|---|---|---|---|---|---|---|---|---|---|
| ENSGACG00000009211 | ChrXVIII | 8793962 | 8796629 | si:dkey-65b13.13 | 1 | 0 | 0 | 0 | 1 | 0 |
| ENSGACG00000009292 | ChrXVII | 8863605 | 8869133 | slc2a9l1 (1 of many) | 1 | 0 | 0 | 0 | 1 | 0 |
| ENSGACG00000009317 | ChrXVII | 8948722 | 8981993 | adamts9 | 1 | 0 | 0 | 0 | 1 | 0 |
| ENSGACG00000009482 | ChrXV | 7346370 | 7363708 | rcan3 | 1 | 0 | 0 | 0 | 0 | 1 |
| ENSGACG00000009483 | ChrVIII | 11917528 | 11996766 | negr1 | 1 | 0 | 0 | 0 | 1 | 0 |
| ENSGACG00000009569 | ChrXII | 11864095 | 11913150 | | 1 | 0 | 0 | 0 | 1 | 0 |
| ENSGACG00000009658 | ChrVI | 11660215 | 11671041 | ZRANB1 (1 of many) | 1 | 0 | 0 | 0 | 1 | 0 |
| ENSGACG00000009650 | ChrXV | 7669664 | 7725769 | qkia | 1 | 1 | 1 | 0 | 0 | 0 |
| ENSGACG00000009678 | ChrXV | 7921527 | 7922252 | | 1 | 0 | 0 | 0 | 1 | 0 |
| ENSGACG00000009836 | ChrXII | 12234412 | 12236332 | | 1 | 0 | 0 | 0 | 1 | 0 |
| ENSGACG00000009889 | ChrX | 15258511 | 15263294 | psmb4 | 1 | 0 | 0 | 0 | 0 | 1 |
| ENSGACG00000009945 | ChrXVII | 9984936 | 9992528 | BMP7 (1 of many) | 1 | 0 | 0 | 0 | 1 | 0 |
| ENSGACG00000009991 | ChrXVIII | 9665614 | 9671172 | tagapb | 1 | 0 | 0 | 0 | 0 | 1 |
| ENSGACG00000009996 | ChrXVII | 10262746 | 10278516 | dido1 | 1 | 0 | 0 | 0 | 1 | 0 |
| ENSGACG00000010079 | ChrXVII | 10406369 | 10422223 | zgc:92107 | 1 | 0 | 0 | 0 | 1 | 0 |
| ENSGACG00000010337 | ChrI | 10680875 | 10691269 | v2ra18 | 1 | 0 | 0 | 0 | 1 | 0 |
| ENSGACG00000010320 | ChrXVII | 10630815 | 10635514 | gnl3l | 1 | 0 | 0 | 0 | 1 | 0 |
| ENSGACG00000010352 | ChrV | 12242543 | 12244965 | | 1 | 0 | 0 | 0 | 1 | 0 |
| ENSGACG00000010363 | ChrVI | 12867904 | 12868960 | sf3b5 | 1 | 0 | 0 | 0 | 1 | 0 |
| ENSGACG00000010489 | ChrVIII | 13247354 | 13252601 | use1 | 1 | 0 | 0 | 0 | 1 | 0 |
| ENSGACG00000010572 | ChrXX | 12085828 | 12089631 | foxj2 | 1 | 0 | 0 | 0 | 1 | 0 |
| ENSGACG00000010616 | ChrVIII | 13406915 | 13410357 | loxl5a | 1 | 0 | 0 | 0 | 1 | 0 |
| ENSGACG00000010598 | ChrXI | 8347830 | 8352415 | rsad1 | 1 | 1 | 0 | 1 | 0 | 0 |
| ENSGACG00000010688 | ChrXV | 9729005 | 9736264 | zgc:101744 | 1 | 0 | 0 | 0 | 1 | 0 |
| ENSGACG00000010643 | ChrXVII | 10920133 | 10925973 | | 1 | 1 | 1 | 0 | 0 | 0 |
| ENSGACG00000010716 | ChrXIII | 11838527 | 11847542 | gda | 1 | 0 | 0 | 0 | 0 | 1 |
| ENSGACG00000010874 | ChrXVIII | 10910812 | 10916238 | ccm2 | 1 | 0 | 0 | 0 | 1 | 0 |
| ENSGACG00000010882 | ChrXVIII | 10919991 | 10923357 | fam167ab | 1 | 0 | 0 | 0 | 1 | 0 |

| | | | | | | | | | | |
|---|---|---|---|---|---|---|---|---|---|---|
| ENSGACG00000010897 | ChrXIII | 12056151 | 12080196 | bmp1a | 1 | 0 | 0 | 0 | 1 | 0 |
| ENSGACG00000010930 | ChrXV | 9957811 | 9977956 | ppp2r3a | 1 | 0 | 0 | 0 | 1 | 0 |
| ENSGACG00000011145 | ChrVIII | 13948367 | 13950370 | dmrta2 | 1 | 0 | 0 | 0 | 1 | 0 |
| ENSGACG00000011152 | ChrVIII | 13997640 | 14027578 | elavl4 | 1 | 1 | 1 | 0 | 0 | 0 |
| ENSGACG00000011194 | ChrXII | 14916639 | 14926699 | syt6a | 1 | 1 | 0 | 1 | 0 | 0 |
| ENSGACG00000011202 | ChrVI | 14449274 | 14452562 | | 1 | 0 | 0 | 0 | 1 | 0 |
| ENSGACG00000011285 | ChrVI | 14524837 | 14528348 | | 1 | 0 | 0 | 0 | 1 | 0 |
| ENSGACG00000011353 | ChrXII | 15159858 | 15171195 | mical1 | 1 | 0 | 0 | 0 | 0 | 1 |
| ENSGACG00000011386 | ChrXV | 11043345 | 11061560 | dtnbb | 1 | 0 | 0 | 0 | 1 | 0 |
| ENSGACG00000011404 | ChrXIII | 12942202 | 12948571 | | 1 | 0 | 0 | 0 | 1 | 0 |
| ENSGACG00000011408 | ChrXI | 10141661 | 10156859 | pdgfab | 1 | 0 | 0 | 0 | 1 | 0 |
| ENSGACG00000011418 | ChrXVII | 12394671 | 12396051 | mrps33 | 1 | 1 | 1 | 0 | 0 | 0 |
| ENSGACG00000011472 | ChrVIII | 14527394 | 14529615 | | 1 | 1 | 0 | 1 | 0 | 0 |
| ENSGACG00000011496 | ChrI | 13367609 | 13368755 | | 1 | 0 | 0 | 0 | 1 | 0 |
| ENSGACG00000011517 | ChrXVIII | 11751680 | 11752669 | gpr6 | 1 | 1 | 1 | 0 | 0 | 0 |
| ENSGACG00000011596 | ChrI | 13626289 | 13631703 | spartb | 1 | 0 | 0 | 0 | 1 | 0 |
| ENSGACG00000011621 | ChrXX | 12578998 | 12584703 | | 1 | 0 | 0 | 0 | 1 | 0 |
| ENSGACG00000011657 | ChrXX | 12630689 | 12639183 | cyp11c1 | 1 | 1 | 0 | 1 | 0 | 0 |
| ENSGACG00000011694 | ChrXI | 10642924 | 10653747 | axin2 | 1 | 0 | 0 | 0 | 1 | 0 |
| ENSGACG00000011737 | ChrXI | 10793354 | 10795700 | cbx8a | 1 | 0 | 0 | 0 | 1 | 0 |
| ENSGACG00000011809 | ChrXVII | 13822213 | 13835193 | rybpa | 1 | 0 | 0 | 0 | 1 | 0 |
| ENSGACG00000011892 | ChrXVII | 14100397 | 14121542 | frmd4ba | 1 | 0 | 0 | 0 | 1 | 0 |
| ENSGACG00000011900 | ChrXVIII | 12613115 | 12627157 | PRKD3 | 1 | 0 | 0 | 0 | 1 | 0 |
| ENSGACG00000011935 | ChrXVIII | 12659540 | 12662849 | r0seh1 | 1 | 0 | 0 | 0 | 0 | 1 |
| ENSGACG00000012038 | ChrXX | 13088418 | 13095119 | ef03b | 1 | 0 | 0 | 0 | 1 | 0 |
| ENSGACG00000012043 | ChrXX | 13166502 | 13166753 | si:dkey-246i14.3 | 1 | 0 | 0 | 0 | 0 | 1 |
| ENSGACG00000012063 | ChrXV | 11951816 | 11991325 | actn1 | 1 | 0 | 0 | 0 | 0 | 1 |
| ENSGACG00000012160 | ChrVI | 16702488 | 16708055 | abhd12 | 1 | 0 | 0 | 0 | 1 | 0 |
| ENSGACG00000012162 | ChrI | 14817967 | 14822383 | pgr | 1 | 0 | 0 | 0 | 0 | 1 |

| | | | | | | | | | | |
|---|---|---|---|---|---|---|---|---|---|---|
| ENSGACG00000012118 | ChrVIII | 15437418 | 15442478 | elovl8b (1 of many) | 1 | 0 | 0 | 0 | 1 | 0 |
| ENSGACG00000012320 | ChrXI | 11499498 | 11522052 | tnrc6c1 | 1 | 0 | 0 | 0 | 1 | 0 |
| ENSGACG00000012326 | ChrVIII | 15624530 | 15629529 | abhd17aa | 1 | 0 | 0 | 0 | 1 | 0 |
| ENSGACG00000012401 | ChrXVIII | 13839871 | 13882401 | klhl29 | 1 | 0 | 0 | 0 | 1 | 0 |
| ENSGACG00000012413 | ChrXX | 13626490 | 13629746 | si:dkey-199f5.8 | 1 | 0 | 0 | 0 | 1 | 0 |
| ENSGACG00000012443 | ChrXX | 13677919 | 13684983 | ago1 | 1 | 0 | 0 | 0 | 1 | 0 |
| ENSGACG00000012473 | ChrXX | 13875776 | 13876264 | hamp (1 of many) | 1 | 0 | 0 | 0 | 1 | 0 |
| ENSGACG00000012649 | ChrVIII | 15997577 | 15998999 | | 1 | 0 | 0 | 0 | 1 | 0 |
| ENSGACG00000012757 | ChrXI | 12294325 | 12298638 | 0CC1 | 1 | 0 | 0 | 0 | 1 | 0 |
| ENSGACG00000012760 | ChrVIII | 16214608 | 16225148 | si:dkey-110c1.10 | 1 | 0 | 0 | 0 | 1 | 0 |
| ENSGACG00000012754 | ChrXX | 14531859 | 14563311 | agmo | 1 | 0 | 0 | 0 | 0 | 1 |
| ENSGACG00000012786 | ChrXX | 14572594 | 14617822 | dgkb | 1 | 0 | 0 | 0 | 1 | 0 |
| ENSGACG00000012863 | ChrIII | 320213 | 322935 | CXorf38 | 1 | 0 | 0 | 0 | 1 | 0 |
| ENSGACG00000012876 | ChrVIII | 16469522 | 16475075 | rgmd | 1 | 0 | 0 | 0 | 1 | 0 |
| ENSGACG00000012938 | ChrXVIII | 14983710 | 14985734 | chac1 | 1 | 0 | 0 | 0 | 1 | 0 |
| ENSGACG00000012944 | ChrIII | 448107 | 459938 | | 1 | 0 | 0 | 0 | 1 | 0 |
| ENSGACG00000013008 | ChrXX | 15116693 | 15118431 | | 1 | 0 | 0 | 0 | 1 | 0 |
| ENSGACG00000013036 | ChrVIII | 16820744 | 16827630 | cilp2 | 1 | 0 | 0 | 0 | 1 | 0 |
| ENSGACG00000013069 | ChrIII | 565595 | 593939 | adcy1b | 1 | 0 | 0 | 0 | 1 | 0 |
| ENSGACG00000013071 | ChrVIII | 16932446 | 16936340 | tmem161a | 1 | 0 | 0 | 0 | 1 | 0 |
| ENSGACG00000013153 | ChrXII | 18049614 | 18054108 | | 1 | 0 | 0 | 0 | 1 | 0 |
| ENSGACG00000013156 | ChrXV | 15523603 | 15528254 | commd8 | 1 | 0 | 0 | 0 | 1 | 0 |
| ENSGACG00000013168 | ChrIII | 815615 | 833164 | CDH2 (1 of many) | 1 | 0 | 0 | 0 | 0 | 1 |
| ENSGACG00000013215 | ChrXIII | 16093221 | 16101732 | mxd1 | 1 | 1 | 1 | 0 | 0 | 0 |
| ENSGACG00000013228 | ChrI | 17134277 | 17138229 | | 1 | 1 | 0 | 1 | 0 | 0 |
| ENSGACG00000013248 | ChrXVIII | 15399767 | 15421191 | nkain2 | 1 | 1 | 1 | 0 | 0 | 0 |
| ENSGACG00000013291 | ChrXIII | 16174644 | 16177810 | znf703 | 1 | 0 | 0 | 0 | 1 | 0 |
| ENSGACG00000013297 | ChrVIII | 17242910 | 17245367 | rab11ba | 1 | 0 | 0 | 0 | 0 | 1 |

| | | | | | | | | | | |
|---|---|---|---|---|---|---|---|---|---|---|
| ENSGACG00000013330 | ChrXX | 16121368 | 16124233 | tspan13a | 1 | 0 | 0 | 0 | 1 | 0 |
| ENSGACG00000013346 | ChrXI | 12863133 | 12864817 | | 1 | 0 | 0 | 0 | 1 | 0 |
| ENSGACG00000013389 | ChrXVIII | 15662633 | 15665757 | 0ga | 1 | 0 | 0 | 0 | 1 | 0 |
| ENSGACG00000013413 | ChrXVIII | 15692426 | 15698913 | si:ch211-57i17.1 | 1 | 1 | 0 | 1 | 0 | 0 |
| ENSGACG00000013478 | scaffold_196 | 78142 | 81595 | | 1 | 0 | 0 | 0 | 1 | 0 |
| ENSGACG00000013492 | ChrIII | 1536538 | 1543454 | | 1 | 1 | 0 | 1 | 0 | 0 |
| ENSGACG00000013537 | ChrVIII | 17456886 | 17459393 | s1pr1 | 1 | 0 | 0 | 0 | 1 | 0 |
| ENSGACG00000013691 | ChrXX | 17223600 | 17228134 | rspo1 | 1 | 0 | 0 | 0 | 0 | 1 |
| ENSGACG00000013737 | ChrXX | 17284737 | 17296628 | | 1 | 0 | 0 | 0 | 1 | 0 |
| ENSGACG00000013866 | ChrXX | 17717428 | 17731718 | inpp5b | 1 | 0 | 0 | 0 | 1 | 0 |
| ENSGACG00000013905 | ChrXIII | 17465408 | 17466387 | | 1 | 0 | 0 | 0 | 0 | 1 |
| ENSGACG00000013933 | ChrI | 20491288 | 20509048 | xirp2a | 1 | 0 | 0 | 0 | 1 | 0 |
| ENSGACG00000013943 | ChrIII | 2351398 | 2375649 | | 1 | 0 | 0 | 0 | 1 | 0 |
| ENSGACG00000013973 | ChrXIII | 17562120 | 17593351 | notch1b | 1 | 0 | 0 | 0 | 0 | 1 |
| ENSGACG00000013977 | ChrVIII | 18160785 | 18167767 | SCML2 (1 of many) | 1 | 1 | 1 | 0 | 0 | 0 |
| ENSGACG00000014007 | ChrXIII | 17704841 | 17747886 | col27a1b | 1 | 0 | 0 | 0 | 0 | 1 |
| ENSGACG00000014040 | ChrVIII | 18275004 | 18275726 | | 1 | 0 | 0 | 0 | 1 | 0 |
| ENSGACG00000014173 | ChrXIII | 18210034 | 18254621 | ksr2 | 1 | 0 | 0 | 0 | 1 | 0 |
| ENSGACG00000014216 | ChrI | 21155272 | 21181371 | farp1 | 1 | 0 | 0 | 0 | 0 | 1 |
| ENSGACG00000014252 | ChrXIII | 18343004 | 18348481 | suds3 | 1 | 1 | 1 | 0 | 0 | 0 |
| ENSGACG00000014284 | ChrXIII | 18582295 | 18583087 | | 1 | 0 | 0 | 0 | 1 | 0 |
| ENSGACG00000014303 | ChrIII | 3571387 | 3812973 | astn1 | 1 | 1 | 1 | 0 | 0 | 0 |
| ENSGACG00000014314 | scaffold_112 | 27482 | 36338 | HTRA1 | 1 | 0 | 0 | 0 | 0 | 1 |
| ENSGACG00000014334 | ChrXIII | 18869454 | 18873205 | tmem175 | 1 | 0 | 0 | 0 | 1 | 0 |
| ENSGACG00000014343 | ChrVIII | 18848370 | 18849326 | | 1 | 0 | 0 | 0 | 1 | 0 |
| ENSGACG00000014350 | ChrXIII | 18889876 | 18895161 | ascc2 | 1 | 1 | 1 | 0 | 0 | 0 |
| ENSGACG00000014353 | ChrI | 21861214 | 21861551 | | 1 | 0 | 0 | 0 | 1 | 0 |
| ENSGACG00000014471 | ChrXIII | 19003720 | 19035311 | slc4a4a | 1 | 0 | 0 | 0 | 1 | 0 |

| | | | | | | | | | | |
|---|---|---|---|---|---|---|---|---|---|---|
| ENSGACG00000014505 | scaffold_112 | 343720 | 345162 | aqp8a,2 | 1 | 0 | 0 | 0 | 0 | 1 |
| ENSGACG00000014620 | scaffold_76 | 287090 | 304975 | pitx3 | 1 | 0 | 0 | 0 | 1 | 0 |
| ENSGACG00000014736 | ChrXIII | 19874116 | 19881489 | | 1 | 0 | 0 | 0 | 0 | 1 |
| ENSGACG00000014814 | ChrII | 5076191 | 5081585 | | 1 | 0 | 0 | 0 | 1 | 0 |
| ENSGACG00000014832 | ChrII | 5125390 | 5127551 | gnrhr2 | 1 | 0 | 0 | 0 | 1 | 0 |
| ENSGACG00000014892 | ChrIII | 6063746 | 6077226 | | 1 | 0 | 0 | 0 | 0 | 1 |
| ENSGACG00000014899 | scaffold_74 | 165635 | 168241 | gpd1c | 1 | 0 | 0 | 0 | 1 | 0 |
| ENSGACG00000014963 | ChrI | 23565974 | 23566795 | | 1 | 0 | 0 | 0 | 1 | 0 |
| ENSGACG00000014987 | ChrXI | 16469732 | 16473523 | tsen54 | 1 | 0 | 0 | 0 | 0 | 1 |
| ENSGACG00000015056 | ChrI | 24829395 | 24935529 | | 1 | 0 | 0 | 0 | 1 | 0 |
| ENSGACG00000015086 | scaffold_68 | 309394 | 354193 | CDH4 (1 of many) | 1 | 0 | 0 | 0 | 1 | 0 |
| ENSGACG00000015165 | ChrIII | 7054385 | 7057639 | PRKAB2 | 1 | 0 | 0 | 0 | 1 | 0 |
| ENSGACG00000015177 | ChrI | 25571569 | 25595585 | map4k4 | 1 | 0 | 0 | 0 | 1 | 0 |
| ENSGACG00000015260 | ChrII | 6665264 | 6668296 | pcdh9 | 1 | 0 | 0 | 0 | 1 | 0 |
| ENSGACG00000015262 | ChrII | 7099598 | 7102887 | PCDH20 | 1 | 0 | 0 | 0 | 1 | 0 |
| ENSGACG00000015264 | ChrII | 7115287 | 7128545 | tdrd3 | 1 | 0 | 0 | 0 | 1 | 0 |
| ENSGACG00000015273 | ChrIII | 7416267 | 7419629 | map2k2b | 1 | 0 | 0 | 0 | 1 | 0 |
| ENSGACG00000015281 | ChrII | 7632856 | 7635853 | trmt10c | 1 | 1 | 1 | 0 | 0 | 0 |
| ENSGACG00000015302 | ChrII | 7675747 | 7702944 | rasa3 | 1 | 0 | 0 | 0 | 1 | 0 |
| ENSGACG00000015374 | ChrII | 7942950 | 8012640 | | 1 | 0 | 0 | 0 | 1 | 0 |
| ENSGACG00000015377 | ChrIII | 7710663 | 7721459 | | 1 | 0 | 0 | 0 | 1 | 0 |
| ENSGACG00000015381 | ChrIII | 7730160 | 7749741 | epha4a | 1 | 0 | 0 | 0 | 1 | 0 |
| ENSGACG00000015490 | ChrXIV | 174415 | 177418 | ufc1 | 1 | 0 | 0 | 0 | 1 | 0 |
| ENSGACG00000015516 | ChrI | 27641497 | 27646007 | | 1 | 0 | 0 | 0 | 0 | 1 |
| ENSGACG00000015581 | ChrI | 27801691 | 27803793 | sumo1 | 1 | 0 | 0 | 0 | 1 | 0 |
| ENSGACG00000015617 | ChrI | 27906006 | 27925118 | ahr2 | 1 | 0 | 0 | 0 | 1 | 0 |
| ENSGACG00000015651 | ChrXIV | 608441 | 613228 | mat2al | 1 | 1 | 0 | 1 | 0 | 0 |
| ENSGACG00000015736 | ChrXIV | 723869 | 725546 | mzt2b | 1 | 0 | 0 | 0 | 0 | 1 |
| ENSGACG00000015775 | ChrXIV | 1031417 | 1038996 | ARSB | 1 | 1 | 0 | 1 | 0 | 0 |

| | | | | | | | | | | |
|---|---|---|---|---|---|---|---|---|---|---|
| ENSGACG00000015791 | ChrII | 11127315 | 11148839 | PHLPP2 | 1 | 0 | 0 | 0 | 1 | 0 |
| ENSGACG00000015831 | ChrII | 11422567 | 11478619 | tcf12 | 1 | 0 | 0 | 0 | 1 | 0 |
| ENSGACG00000015862 | ChrXIV | 1681568 | 1684183 | npffr2b | 1 | 1 | 1 | 0 | 0 | 0 |
| ENSGACG00000015884 | ChrXIV | 1751442 | 1766482 | erbin | 1 | 0 | 0 | 0 | 1 | 0 |
| ENSGACG00000015902 | ChrXIV | 1944466 | 1968159 | fnbp1b | 1 | 1 | 0 | 1 | 0 | 0 |
| ENSGACG00000015915 | ChrXIV | 2174485 | 2181252 | ttll11 | 1 | 0 | 0 | 0 | 1 | 0 |
| ENSGACG00000015949 | ChrXIV | 2711343 | 2720198 | usp20 | 1 | 0 | 0 | 0 | 0 | 1 |
| ENSGACG00000015999 | ChrIX | 791090 | 795086 | | 1 | 0 | 0 | 0 | 1 | 0 |
| ENSGACG00000016065 | ChrIII | 9230992 | 9232131 | b3gnt5b | 1 | 0 | 0 | 0 | 1 | 0 |
| ENSGACG00000016073 | ChrIX | 1334639 | 1357056 | | 1 | 0 | 0 | 0 | 0 | 1 |
| ENSGACG00000016089 | ChrXIV | 3138466 | 3139400 | rpl28 | 1 | 0 | 0 | 0 | 0 | 1 |
| ENSGACG00000016100 | ChrIX | 1429086 | 1441309 | polq | 1 | 0 | 0 | 0 | 1 | 0 |
| ENSGACG00000016150 | ChrII | 13614114 | 13615977 | psmc3 | 1 | 0 | 0 | 0 | 1 | 0 |
| ENSGACG00000016142 | ChrXIV | 3217562 | 3221176 | atp5fa1 (1 of many) | 1 | 1 | 1 | 0 | 0 | 0 |
| ENSGACG00000016163 | ChrIII | 9641348 | 9647872 | gpbp1l1 | 1 | 0 | 0 | 0 | 1 | 0 |
| ENSGACG00000016169 | ChrXIV | 3251935 | 3256350 | | 1 | 0 | 0 | 0 | 0 | 1 |
| ENSGACG00000016193 | ChrIX | 1782128 | 1785348 | chr09 | 1 | 0 | 0 | 0 | 1 | 0 |
| ENSGACG00000016222 | ChrXIV | 3365899 | 3371406 | kcmf1 | 1 | 0 | 0 | 0 | 1 | 0 |
| ENSGACG00000016347 | ChrIX | 3127102 | 3143261 | p2rx3b | 1 | 0 | 0 | 0 | 1 | 0 |
| ENSGACG00000016360 | ChrIV | 472586 | 477561 | pcdh1b | 1 | 0 | 0 | 0 | 0 | 1 |
| ENSGACG00000016426 | ChrIV | 1302365 | 1304893 | fgfrl1a | 1 | 1 | 1 | 0 | 0 | 0 |
| ENSGACG00000016451 | ChrII | 15348070 | 15353934 | CTSH | 1 | 0 | 0 | 0 | 1 | 0 |
| ENSGACG00000016473 | ChrXIV | 3993506 | 3998272 | prlra | 1 | 0 | 0 | 0 | 1 | 0 |
| ENSGACG00000016521 | ChrII | 15654499 | 15662194 | fam214a | 1 | 0 | 0 | 0 | 0 | 1 |
| ENSGACG00000016606 | ChrIX | 4603846 | 4604358 | | 1 | 0 | 0 | 0 | 1 | 0 |
| ENSGACG00000016616 | ChrIX | 4624437 | 4630228 | mief2 | 1 | 0 | 0 | 0 | 1 | 0 |
| ENSGACG00000016624 | ChrIV | 2464056 | 2509522 | rapgef2 | 1 | 0 | 0 | 0 | 1 | 0 |
| ENSGACG00000016708 | ChrIX | 4885702 | 4887717 | | 1 | 0 | 0 | 0 | 1 | 0 |

| | | | | | | | | | | |
|---|---|---|---|---|---|---|---|---|---|---|
| ENSGACG00000016723 | ChrIX | 5098704 | 5108431 | g012a | 1 | 0 | 0 | 0 | 1 | 0 |
| ENSGACG00000016746 | ChrII | 18045594 | 18048728 | gldn | 1 | 1 | 1 | 0 | 0 | 0 |
| ENSGACG00000016846 | ChrIX | 5456262 | 5476760 | | 1 | 1 | 1 | 0 | 0 | 0 |
| ENSGACG00000016916 | ChrXIV | 6417172 | 6426532 | sgsm1b | 1 | 0 | 0 | 0 | 1 | 0 |
| ENSGACG00000016918 | ChrIII | 12322870 | 12341345 | | 1 | 1 | 1 | 0 | 0 | 0 |
| ENSGACG00000016952 | ChrII | 19109323 | 19110312 | rxfp3,3a2 | 1 | 0 | 0 | 0 | 0 | 1 |
| ENSGACG00000016957 | ChrII | 19149179 | 19152927 | calml4a | 1 | 1 | 0 | 1 | 0 | 0 |
| ENSGACG00000016959 | ChrXIV | 6533608 | 6537690 | svopb | 1 | 0 | 0 | 0 | 0 | 1 |
| ENSGACG00000016992 | ChrII | 19281893 | 19298785 | znf609b | 1 | 0 | 0 | 0 | 1 | 0 |
| ENSGACG00000017000 | ChrII | 19349048 | 19401226 | myo9aa | 1 | 0 | 0 | 0 | 1 | 0 |
| ENSGACG00000017063 | ChrII | 19650101 | 19656297 | | 1 | 0 | 0 | 0 | 1 | 0 |
| ENSGACG00000017074 | ChrIV | 5485973 | 5494737 | efemp2a | 1 | 0 | 0 | 0 | 1 | 0 |
| ENSGACG00000017077 | ChrIV | 5517696 | 5539310 | mrpl11 | 1 | 0 | 0 | 0 | 1 | 0 |
| ENSGACG00000017096 | ChrII | 20027174 | 20041230 | slc6a5 | 1 | 0 | 0 | 0 | 1 | 0 |
| ENSGACG00000017124 | ChrIX | 6068561 | 6084066 | | 1 | 0 | 0 | 0 | 1 | 0 |
| ENSGACG00000017160 | ChrII | 21080854 | 21084821 | | 1 | 0 | 0 | 0 | 0 | 1 |
| ENSGACG00000017209 | ChrXIV | 6969603 | 6972066 | pcyox1 | 1 | 0 | 0 | 0 | 1 | 0 |
| ENSGACG00000017255 | ChrIX | 7204599 | 7294154 | inpp4b | 1 | 0 | 0 | 0 | 1 | 0 |
| ENSGACG00000017313 | ChrII | 21709930 | 21734771 | MEGF11 (1 of many) | 1 | 1 | 0 | 1 | 0 | 0 |
| ENSGACG00000017319 | ChrII | 21769057 | 21771819 | socs4 | 1 | 1 | 1 | 0 | 0 | 0 |
| ENSGACG00000017384 | ChrII | 22035294 | 22037750 | ndufs3 | 1 | 0 | 0 | 0 | 1 | 0 |
| ENSGACG00000017389 | ChrXIV | 7665108 | 7673135 | urm1 | 1 | 0 | 0 | 0 | 1 | 0 |
| ENSGACG00000017407 | ChrXIV | 7749442 | 7751978 | fam163b | 1 | 0 | 0 | 0 | 1 | 0 |
| ENSGACG00000017464 | ChrII | 22240099 | 22241218 | | 1 | 0 | 0 | 0 | 1 | 0 |
| ENSGACG00000017476 | ChrII | 22301954 | 22308632 | cpeb1b | 1 | 0 | 0 | 0 | 0 | 1 |
| ENSGACG00000017506 | ChrII | 22373022 | 22375772 | nqo1 (1 of many) | 1 | 0 | 0 | 0 | 1 | 0 |
| ENSGACG00000017504 | ChrIV | 7553451 | 7640413 | enox2 | 1 | 0 | 0 | 0 | 1 | 0 |
| ENSGACG00000017542 | ChrII | 22570349 | 22571189 | | 1 | 0 | 0 | 0 | 1 | 0 |

| | | | | | | | | | | |
|---|---|---|---|---|---|---|---|---|---|---|
| ENSGACG00000017563 | ChrIV | 8075659 | 8076747 | | 1 | 0 | 0 | 0 | 1 | 0 |
| ENSGACG00000017568 | ChrIII | 14440090 | 14454692 | PFKP (1 of many) | 1 | 0 | 0 | 0 | 0 | 1 |
| ENSGACG00000017613 | ChrII | 22859068 | 22883630 | ppfia1 | 1 | 0 | 0 | 0 | 1 | 0 |
| ENSGACG00000017647 | ChrXIV | 8597952 | 8633437 | sfswap | 1 | 0 | 0 | 0 | 0 | 1 |
| ENSGACG00000017689 | ChrIV | 8520482 | 8545266 | si:ch211-220e11.3 | 1 | 0 | 0 | 0 | 1 | 0 |
| ENSGACG00000017699 | ChrXIV | 8996130 | 9029956 | ank1b | 1 | 0 | 0 | 0 | 0 | 1 |
| ENSGACG00000017713 | ChrIII | 15794470 | 15804479 | | 1 | 0 | 0 | 0 | 1 | 0 |
| ENSGACG00000017773 | ChrIV | 9228627 | 9233379 | rad9a | 1 | 0 | 0 | 0 | 1 | 0 |
| ENSGACG00000017893 | ChrIII | 16251876 | 16277818 | cac01eb | 1 | 0 | 0 | 0 | 1 | 0 |
| ENSGACG00000017924 | ChrXIV | 10388550 | 10398180 | ghrb | 1 | 0 | 0 | 0 | 1 | 0 |
| ENSGACG00000017934 | ChrIV | 10439115 | 10505618 | si:dkey-201i24.6 | 1 | 1 | 1 | 0 | 0 | 0 |
| ENSGACG00000017983 | ChrIII | 16717673 | 16723180 | atg9b | 1 | 1 | 1 | 0 | 0 | 0 |
| ENSGACG00000017998 | ChrIV | 10730374 | 10732632 | arsia | 1 | 0 | 0 | 0 | 1 | 0 |
| ENSGACG00000018028 | ChrIX | 9645774 | 9651002 | spock3 | 1 | 1 | 1 | 0 | 0 | 0 |
| ENSGACG00000018121 | ChrXIV | 11304434 | 11333562 | grin1a | 1 | 0 | 0 | 0 | 1 | 0 |
| ENSGACG00000018145 | ChrXIV | 11471663 | 11476209 | barhl1b | 1 | 0 | 0 | 0 | 1 | 0 |
| ENSGACG00000018172 | ChrXIV | 11597815 | 11602791 | adra1ab | 1 | 0 | 0 | 0 | 0 | 1 |
| ENSGACG00000018179 | ChrXIV | 11850201 | 11893984 | vav2 | 1 | 0 | 0 | 0 | 1 | 0 |
| ENSGACG00000018233 | ChrIX | 10713754 | 10725152 | pdcd11 | 1 | 1 | 1 | 0 | 0 | 0 |
| ENSGACG00000018231 | ChrIV | 12013740 | 12034920 | abcb7 | 1 | 0 | 0 | 0 | 1 | 0 |
| ENSGACG00000018322 | ChrIV | 12941555 | 12943706 | | 1 | 0 | 0 | 0 | 1 | 0 |
| ENSGACG00000018323 | ChrXIV | 13873722 | 13900598 | lmx1ba | 1 | 0 | 0 | 0 | 0 | 1 |
| ENSGACG00000018335 | ChrXIV | 13922425 | 13930656 | mapkap1 | 1 | 0 | 0 | 0 | 1 | 0 |
| ENSGACG00000018341 | ChrIX | 11059538 | 11064667 | ogal | 1 | 1 | 1 | 0 | 0 | 0 |
| ENSGACG00000018360 | ChrIX | 11194695 | 11200674 | lrpap1 | 1 | 0 | 0 | 0 | 1 | 0 |
| ENSGACG00000018375 | ChrXIV | 14135402 | 14160309 | | 1 | 1 | 0 | 1 | 0 | 0 |
| ENSGACG00000018409 | ChrIX | 11666673 | 11682015 | pcdh10a | 1 | 0 | 0 | 0 | 1 | 0 |
| ENSGACG00000018461 | ChrIV | 14219527 | 14248769 | ablim3 | 1 | 0 | 0 | 0 | 1 | 0 |
| ENSGACG00000018600 | ChrVII | 242336 | 245534 | | 1 | 1 | 1 | 0 | 0 | 0 |

| | | | | | | | | | | |
|---|---|---|---|---|---|---|---|---|---|---|
| ENSGACG00000018633 | ChrVII | 365776 | 372621 | | 1 | 0 | 0 | 0 | 1 | 0 |
| ENSGACG00000018652 | ChrVII | 494053 | 497077 | | 1 | 0 | 0 | 0 | 1 | 0 |
| ENSGACG00000018663 | ChrIV | 16118651 | 16126776 | tmem173 | 1 | 0 | 0 | 0 | 1 | 0 |
| ENSGACG00000018759 | ChrVII | 835005 | 836654 | | 1 | 0 | 0 | 0 | 0 | 1 |
| ENSGACG00000018788 | ChrVII | 1108957 | 1111978 | tmem185 | 1 | 1 | 0 | 1 | 0 | 0 |
| ENSGACG00000018802 | ChrVII | 1133367 | 1137045 | | 1 | 0 | 0 | 0 | 1 | 0 |
| ENSGACG00000018836 | ChrIV | 18419217 | 18424359 | sypl1 | 1 | 1 | 1 | 0 | 0 | 0 |
| ENSGACG00000018841 | ChrIV | 18438511 | 18458268 | gsap | 1 | 0 | 0 | 0 | 1 | 0 |
| ENSGACG00000018867 | ChrVII | 1452630 | 1459173 | khnyn | 1 | 0 | 0 | 0 | 0 | 1 |
| ENSGACG00000018876 | ChrVII | 1481900 | 1484089 | emc4 | 1 | 0 | 0 | 0 | 0 | 1 |
| ENSGACG00000018929 | ChrIV | 19626362 | 19645224 | | 1 | 0 | 0 | 0 | 1 | 0 |
| ENSGACG00000018991 | ChrVII | 2259206 | 2267772 | zgc:55262 | 1 | 0 | 0 | 0 | 0 | 1 |
| ENSGACG00000019026 | ChrIV | 20833178 | 20835191 | alg10 | 1 | 0 | 0 | 0 | 0 | 1 |
| ENSGACG00000019034 | ChrIV | 20945694 | 20956175 | abcd2 | 1 | 0 | 0 | 0 | 1 | 0 |
| ENSGACG00000019043 | ChrIX | 15241216 | 15246931 | pycr1b | 1 | 1 | 1 | 0 | 0 | 0 |
| ENSGACG00000019101 | ChrVII | 2678788 | 2682285 | rgp1 | 1 | 0 | 0 | 0 | 1 | 0 |
| ENSGACG00000019117 | ChrVII | 2701856 | 2708450 | si:dkeyp-75h12.2 | 1 | 0 | 0 | 0 | 1 | 0 |
| ENSGACG00000019125 | ChrIV | 21818906 | 21878888 | SYN3 | 1 | 0 | 0 | 0 | 0 | 1 |
| ENSGACG00000019138 | ChrVII | 2808940 | 2812270 | | 1 | 0 | 0 | 0 | 1 | 0 |
| ENSGACG00000019176 | ChrIV | 22147684 | 22153447 | pfkfb3 | 1 | 0 | 0 | 0 | 1 | 0 |
| ENSGACG00000019227 | ChrIV | 22899577 | 22913555 | mkrn1 | 1 | 0 | 0 | 0 | 0 | 1 |
| ENSGACG00000019253 | ChrIV | 23209606 | 23250489 | | 1 | 1 | 1 | 0 | 0 | 0 |
| ENSGACG00000019286 | ChrVII | 3160049 | 3162174 | si:dkey-19b23.7 | 1 | 0 | 0 | 0 | 1 | 0 |
| ENSGACG00000019353 | ChrIX | 16630442 | 16684711 | MGAT5B (1 of many) | 1 | 0 | 0 | 0 | 1 | 0 |
| ENSGACG00000019362 | ChrVII | 4248635 | 4260779 | neurl4 | 1 | 0 | 0 | 0 | 1 | 0 |
| ENSGACG00000019440 | ChrIV | 24711763 | 24712875 | | 1 | 0 | 0 | 0 | 1 | 0 |
| ENSGACG00000019460 | ChrIX | 17631571 | 17632626 | dand5 | 1 | 0 | 0 | 0 | 1 | 0 |
| ENSGACG00000019472 | ChrIV | 25318371 | 25416178 | 0v3 | 1 | 0 | 0 | 0 | 1 | 0 |

| | | | | | | | | | | |
|---|---|---|---|---|---|---|---|---|---|---|
| ENSGACG00000019498 | ChrVII | 5325617 | 5432198 | ADGRL3 | 1 | 0 | 0 | 0 | 0 | 1 |
| ENSGACG00000019674 | ChrIV | 28658720 | 28716045 | wnk1b | 1 | 0 | 0 | 0 | 1 | 0 |
| ENSGACG00000019679 | ChrIV | 28725325 | 28735256 | | 1 | 0 | 0 | 0 | 1 | 0 |
| ENSGACG00000019805 | ChrIV | 30481780 | 30482190 | btg1 | 1 | 0 | 0 | 0 | 1 | 0 |
| ENSGACG00000019945 | ChrIV | 31129826 | 31165134 | ptn | 1 | 0 | 0 | 0 | 1 | 0 |
| ENSGACG00000019970 | ChrIV | 31495299 | 31507946 | si:dkey-97m3.1 | 1 | 0 | 0 | 0 | 1 | 0 |
| ENSGACG00000020022 | ChrVII | 9404415 | 9405080 | zgc:113305 | 1 | 0 | 0 | 0 | 1 | 0 |
| ENSGACG00000020033 | ChrVII | 10133090 | 10158972 | si:cabz01078036.1 | 1 | 0 | 0 | 0 | 1 | 0 |
| ENSGACG00000020045 | ChrIV | 32117996 | 32125903 | parpbp | 1 | 0 | 0 | 0 | 1 | 0 |
| ENSGACG00000020050 | ChrIV | 32208749 | 32220861 | nrip2 | 1 | 0 | 0 | 0 | 1 | 0 |
| ENSGACG00000020051 | ChrIV | 32222974 | 32224772 | klhl42 | 1 | 0 | 0 | 0 | 1 | 0 |
| ENSGACG00000020174 | ChrVII | 12750327 | 12755673 | capns1a | 1 | 0 | 0 | 0 | 1 | 0 |
| ENSGACG00000020311 | ChrVII | 16430416 | 16478058 | | 1 | 0 | 0 | 0 | 0 | 1 |
| ENSGACG00000020325 | ChrVII | 17024309 | 17031165 | | 1 | 0 | 0 | 0 | 1 | 0 |
| ENSGACG00000020369 | ChrVII | 17649875 | 17653814 | | 1 | 0 | 0 | 0 | 1 | 0 |
| ENSGACG00000020401 | ChrVII | 17997085 | 18001531 | si:ch211-137i24.10 | 1 | 1 | 0 | 1 | 0 | 0 |
| ENSGACG00000020426 | ChrVII | 18408108 | 18413324 | taf6 | 1 | 0 | 0 | 0 | 0 | 1 |
| ENSGACG00000020448 | ChrVII | 18690919 | 18691724 | | 1 | 1 | 0 | 1 | 0 | 0 |
| ENSGACG00000020556 | ChrVII | 20054776 | 20129917 | nbeaa | 1 | 0 | 0 | 0 | 1 | 0 |
| ENSGACG00000020577 | ChrVII | 20565349 | 20569644 | eva1c | 1 | 0 | 0 | 0 | 1 | 0 |
| ENSGACG00000020658 | ChrVII | 22028152 | 22042272 | ppp3ca | 1 | 0 | 0 | 0 | 1 | 0 |
| ENSGACG00000020675 | ChrVII | 22206068 | 22209023 | | 1 | 0 | 0 | 0 | 0 | 1 |
| ENSGACG00000020726 | ChrVII | 23208162 | 23253092 | ARHGAP26 | 1 | 0 | 0 | 0 | 1 | 0 |
| ENSGACG00000020747 | ChrVII | 23962472 | 23988220 | dpysl3 | 1 | 0 | 0 | 0 | 1 | 0 |
| ENSGACG00000020803 | ChrVII | 25380828 | 25384895 | | 1 | 0 | 0 | 0 | 1 | 0 |
| ENSGACG00000020805 | ChrVII | 25418091 | 25465731 | dbn1 | 1 | 1 | 1 | 0 | 0 | 0 |
| ENSGACG00000020811 | ChrVII | 25541259 | 25547799 | | 1 | 0 | 0 | 0 | 1 | 0 |
| ENSGACG00000020824 | ChrVII | 26193351 | 26195752 | cnot8 | 1 | 1 | 0 | 1 | 0 | 0 |

| | | | | | | | | | | |
|---|---|---|---|---|---|---|---|---|---|---|
| ENSGACG00000020842 | ChrVII | 26490578 | 26493742 | rbm41 | 1 | 0 | 0 | 0 | 1 | 0 |
| ENSGACG00000020848 | ChrVII | 26791004 | 26845216 | doc2b | 1 | 0 | 0 | 0 | 0 | 1 |
| ENSGACG00000020849 | ChrVII | 26855161 | 26860699 | gatsl2 | 1 | 0 | 0 | 0 | 1 | 0 |
| ENSGACG00000021308 | ChrIII | 9959732 | 9959817 | | 1 | 0 | 0 | 0 | 1 | 0 |
| ENSGACG00000021339 | ChrXIII | 17724932 | 17725023 | | 1 | 0 | 0 | 0 | 1 | 0 |
| ENSGACG00000021543 | ChrVIII | 10666396 | 10666480 | | 1 | 0 | 0 | 0 | 0 | 1 |
| ENSGACG00000022289 | ChrVII | 19110984 | 19111069 | | 1 | 0 | 0 | 0 | 1 | 0 |
| ENSGACG00000022499 | ChrX | 9755442 | 9755535 | | 1 | 0 | 0 | 0 | 1 | 0 |
| ENSGACG00000022573 | ChrI | 4299459 | 4299517 | | 1 | 0 | 0 | 0 | 0 | 1 |
| ENSGACG00000022581 | ChrX | 9041304 | 9041363 | | 1 | 1 | 1 | 0 | 0 | 0 |
| ENSGACG00000022669 | ChrVIII | 10778399 | 10778538 | RF00191 | 1 | 0 | 0 | 0 | 1 | 0 |
| ENSGACG00000022764 | ChrIX | 5186427 | 5186513 | RF00271 | 1 | 0 | 0 | 0 | 1 | 0 |

**Table S2B:** Differentially methylated genes between populations from Kiel (20 PSU) and Sylt (33 PSU): For genes associated with DMS, Ensembl gene ID and gene name as well as the position on the chromosome are listed. The numbers refer to the numbers of DMS in the population comparison (wild), these DMS were classified into ‘inducible’, ‘inconclusive’ and ‘stable’ sites according to their behavior in a two-generation salinity acclimation experiment with laboratory bred sticklebacks from Kiel (20 PSU) exposed to experimental salinity increase (33 PSU) (see Methods for details). Further, inducible sites were distinguished whether they matched methylation levels of the locally adapted population (‘*expected*’) or not (‘*opposite*’).

| Ensembl gene ID | chromosome | start position | end position | gene name | wild | inducible | expected inducible | opposite inducible | stable | inconclusive |
|---|---|---|---|---|---|---|---|---|---|---|
| ENSGACG00000020323 | ChrVII | 17010160 | 17011176 | | 23 | 0 | 0 | 0 | 22 | 1 |
| ENSGACG00000013229 | ChrXVIII | 15327717 | 15352321 | | 15 | 10 | 10 | 0 | 1 | 4 |
| ENSGACG00000013359 | ChrXI | 12960883 | 12968110 | sec14l1 | 15 | 0 | 0 | 0 | 12 | 3 |
| ENSGACG00000019416 | ChrVII | 4451892 | 4453656 | HMX1 orthologue | 15 | 3 | 3 | 0 | 5 | 7 |
| ENSGACG00000002948 | ChrVIII | 218240 | 221355 | ddx10 | 14 | 0 | 0 | 0 | 6 | 8 |
| ENSGACG00000016350 | ChrXIV | 3603545 | 3604923 | | 14 | 1 | 0 | 1 | 7 | 6 |
| ENSGACG00000006636 | ChrXVIII | 4780893 | 4786820 | ZC3H12D | 13 | 0 | 0 | 0 | 3 | 10 |
| ENSGACG00000004667 | ChrXII | 4273498 | 4286193 | tti1 | 12 | 0 | 0 | 0 | 12 | 0 |
| ENSGACG00000015566 | ChrII | 9043062 | 9051779 | casc4 | 10 | 0 | 0 | 0 | 10 | 0 |
| ENSGACG00000004433 | ChrXVII | 2127457 | 2211376 | igsf21a | 9 | 9 | 9 | 0 | 0 | 0 |
| ENSGACG00000006772 | ChrXV | 3284481 | 3316132 | dlgap2a | 9 | 0 | 0 | 0 | 7 | 2 |
| ENSGACG00000008805 | ChrX | 13142212 | 13171897 | ST3GAL1 (1 of many) | 9 | 0 | 0 | 0 | 8 | 1 |
| ENSGACG00000008919 | ChrX | 13502303 | 13521318 | kcnk9 | 8 | 0 | 0 | 0 | 0 | 8 |
| ENSGACG00000020706 | ChrVII | 22666912 | 22672421 | | 8 | 0 | 0 | 0 | 8 | 0 |
| ENSGACG00000004770 | ChrXVI | 10045160 | 10049909 | SP3 | 7 | 0 | 0 | 0 | 5 | 2 |
| ENSGACG00000014483 | ChrXI | 15438933 | 15454835 | gas7a | 7 | 0 | 0 | 0 | 7 | 0 |

| | | | | | | | | | | |
|---|---|---|---|---|---|---|---|---|---|---|
| ENSGACG00000015862 | ChrXIV | 1681568 | 1684183 | npffr2b | 7 | 0 | 0 | 0 | 7 | 0 |
| ENSGACG00000016341 | ChrIV | 229120 | 240408 | si:dkeyp-110e4.11 | 7 | 0 | 0 | 0 | 0 | 7 |
| ENSGACG00000019897 | ChrIV | 31023275 | 31026361 | ada2b | 7 | 0 | 0 | 0 | 0 | 7 |
| ENSGACG00000006040 | ChrXV | 2368176 | 2420656 | RYR3 (1 of many) | 6 | 3 | 0 | 3 | 0 | 3 |
| ENSGACG00000012193 | ChrXVIII | 13344808 | 13352902 | | 6 | 0 | 0 | 0 | 6 | 0 |
| ENSGACG00000015057 | scaffold_74 | 485843 | 502285 | pkp4 | 6 | 0 | 0 | 0 | 6 | 0 |
| ENSGACG00000017845 | ChrIX | 8726948 | 8730779 | lgi2b | 6 | 0 | 0 | 0 | 0 | 6 |
| ENSGACG00000018110 | ChrIX | 10035146 | 10041439 | krt222 | 6 | 0 | 0 | 0 | 6 | 0 |
| ENSGACG00000018576 | ChrIX | 13131695 | 13177741 | tusc3 | 6 | 0 | 0 | 0 | 0 | 6 |
| ENSGACG00000001675 | scaffold_120 | 284160 | 295979 | LAMC2 | 5 | 0 | 0 | 0 | 2 | 3 |
| ENSGACG00000002963 | ChrXX | 130883 | 136797 | | 5 | 2 | 2 | 0 | 0 | 3 |
| ENSGACG00000005061 | ChrXVI | 10329406 | 10372640 | myo3b | 5 | 0 | 0 | 0 | 0 | 5 |
| ENSGACG00000009482 | ChrXV | 7346370 | 7363708 | rcan3 | 5 | 0 | 0 | 0 | 3 | 2 |
| ENSGACG00000010846 | ChrVIII | 13598498 | 13605960 | lhx4 | 5 | 1 | 1 | 0 | 1 | 3 |
| ENSGACG00000011118 | ChrI | 12219197 | 12219822 | | 5 | 0 | 0 | 0 | 2 | 3 |
| ENSGACG00000011657 | ChrXX | 12630689 | 12639183 | cyp11c1 | 5 | 3 | 0 | 3 | 1 | 1 |
| ENSGACG00000011995 | ChrXV | 11740267 | 11769069 | trim9 | 5 | 0 | 0 | 0 | 2 | 3 |
| ENSGACG00000020047 | ChrIV | 32137226 | 32150243 | th2 | 5 | 0 | 0 | 0 | 5 | 0 |
| ENSGACG00000001000 | scaffold_108 | 236983 | 240230 | hvcn1 | 4 | 0 | 0 | 0 | 4 | 0 |
| ENSGACG00000002150 | scaffold_56 | 991932 | 1000901 | b4galt6 | 4 | 0 | 0 | 0 | 0 | 4 |
| ENSGACG00000003831 | ChrXVII | 1268650 | 1270064 | | 4 | 1 | 1 | 0 | 1 | 2 |
| ENSGACG00000003932 | ChrX | 5372498 | 5374253 | | 4 | 0 | 0 | 0 | 4 | 0 |
| ENSGACG00000008558 | ChrVIII | 9695227 | 9702494 | si:ch211-43f4.1 | 4 | 0 | 0 | 0 | 4 | 0 |
| ENSGACG00000008986 | ChrX | 13667723 | 13673241 | ube2ql1 | 4 | 0 | 0 | 0 | 3 | 1 |
| ENSGACG00000011019 | ChrXI | 9080011 | 9104314 | cyth1a | 4 | 0 | 0 | 0 | 4 | 0 |
| ENSGACG00000017534 | ChrXIV | 8089235 | 8092718 | | 4 | 0 | 0 | 0 | 3 | 1 |
| ENSGACG00000001150 | scaffold_237 | 44323 | 48391 | | 3 | 1 | 1 | 0 | 1 | 1 |
| ENSGACG00000002270 | ChrX | 1627559 | 1629451 | tekt1 | 3 | 0 | 0 | 0 | 1 | 2 |
| ENSGACG00000002707 | ChrXII | 560982 | 576138 | grip2a | 3 | 0 | 0 | 0 | 3 | 0 |

| | | | | | | | | | | |
|---|---|---|---|---|---|---|---|---|---|---|
| ENSGACG00000002865 | ChrXII | 847025 | 855389 | prickle2a | 3 | 0 | 0 | 0 | 2 | 1 |
| ENSGACG00000005778 | ChrXVI | 11091546 | 11093393 | dap1b | 3 | 0 | 0 | 0 | 3 | 0 |
| ENSGACG00000007036 | ChrXIII | 6072435 | 6095546 | | 3 | 0 | 0 | 0 | 3 | 0 |
| ENSGACG00000009367 | ChrV | 11265977 | 11279015 | baiap2l1a | 3 | 0 | 0 | 0 | 2 | 1 |
| ENSGACG00000010367 | ChrXVIII | 10300965 | 10314744 | acss1 | 3 | 0 | 0 | 0 | 3 | 0 |
| ENSGACG00000010998 | ChrXVII | 11632756 | 11638904 | cldn19 | 3 | 0 | 0 | 0 | 0 | 3 |
| ENSGACG00000012122 | ChrXV | 12090680 | 12101068 | map3k9 | 3 | 0 | 0 | 0 | 3 | 0 |
| ENSGACG00000014293 | ChrVIII | 18757229 | 18761036 | tmem165 | 3 | 0 | 0 | 0 | 3 | 0 |
| ENSGACG00000015715 | ChrII | 10395592 | 10421700 | tead1b | 3 | 0 | 0 | 0 | 3 | 0 |
| ENSGACG00000016957 | ChrII | 19149179 | 19152927 | calml4a | 3 | 2 | 0 | 2 | 0 | 1 |
| ENSGACG00000017210 | ChrIII | 13216739 | 13218719 | zgc:113531 | 3 | 0 | 0 | 0 | 1 | 2 |
| ENSGACG00000017236 | ChrII | 21381632 | 21396647 | | 3 | 0 | 0 | 0 | 3 | 0 |
| ENSGACG00000018330 | ChrIX | 10996164 | 11001903 | KCNIP2 | 3 | 0 | 0 | 0 | 1 | 2 |
| ENSGACG00000020808 | ChrVII | 25496687 | 25519233 | grk6 | 3 | 0 | 0 | 0 | 3 | 0 |
| ENSGACG00000021523 | scaffold_80 | 401870 | 401961 | | 3 | 0 | 0 | 0 | 3 | 0 |
| ENSGACG00000000253 | scaffold_114 | 74933 | 109669 | itpr3 | 2 | 0 | 0 | 0 | 2 | 0 |
| ENSGACG00000000935 | scaffold_1182 | 4446 | 4961 | | 2 | 1 | 0 | 1 | 1 | 0 |
| ENSGACG00000000972 | scaffold_37 | 1733384 | 1743429 | lancl2 | 2 | 0 | 0 | 0 | 2 | 0 |
| ENSGACG00000001842 | scaffold_600 | 385 | 1591 | | 2 | 0 | 0 | 0 | 0 | 2 |
| ENSGACG00000002193 | ChrV | 298074 | 298954 | DUSP13 | 2 | 0 | 0 | 0 | 2 | 0 |
| ENSGACG00000002344 | ChrX | 1751605 | 1763088 | nedd9 | 2 | 0 | 0 | 0 | 2 | 0 |
| ENSGACG00000003215 | ChrV | 2545786 | 2550169 | prph2a | 2 | 0 | 0 | 0 | 2 | 0 |
| ENSGACG00000003249 | ChrXII | 1592910 | 1631790 | scube3 | 2 | 0 | 0 | 0 | 1 | 1 |
| ENSGACG00000003868 | ChrXV | 24 | 4126 | | 2 | 0 | 0 | 0 | 2 | 0 |
| ENSGACG00000003922 | ChrVI | 4019342 | 4043269 | | 2 | 0 | 0 | 0 | 2 | 0 |
| ENSGACG00000004040 | ChrXII | 3116439 | 3118541 | | 2 | 0 | 0 | 0 | 0 | 2 |
| ENSGACG00000004212 | ChrXIII | 1654417 | 1666382 | si:dkey-178e17.1 | 2 | 0 | 0 | 0 | 0 | 2 |
| ENSGACG00000004339 | ChrVIII | 2537139 | 2541472 | | 2 | 0 | 0 | 0 | 1 | 1 |
| ENSGACG00000004526 | ChrX | 6917318 | 6925218 | ef03a | 2 | 0 | 0 | 0 | 2 | 0 |

| | | | | | | | | | | |
|---|---|---|---|---|---|---|---|---|---|---|
| ENSGACG00000004657 | ChrXVIII | 1241770 | 1243679 | | 2 | 0 | 0 | 0 | 2 | 0 |
| ENSGACG00000005316 | ChrXII | 5810740 | 5816656 | fam131c | 2 | 0 | 0 | 0 | 2 | 0 |
| ENSGACG00000005631 | ChrXI | 1709563 | 1713578 | hoxb3a | 2 | 0 | 0 | 0 | 2 | 0 |
| ENSGACG00000006038 | ChrXVII | 4531155 | 4544248 | znfx1 | 2 | 0 | 0 | 0 | 1 | 1 |
| ENSGACG00000006245 | ChrXVIII | 3473616 | 3494661 | bach2b | 2 | 0 | 0 | 0 | 0 | 2 |
| ENSGACG00000006328 | ChrXX | 5969700 | 5976203 | | 2 | 0 | 0 | 0 | 2 | 0 |
| ENSGACG00000006387 | ChrXX | 6166089 | 6172879 | entpd3 | 2 | 1 | 0 | 1 | 0 | 1 |
| ENSGACG00000006456 | ChrXVII | 5282637 | 5335165 | | 2 | 0 | 0 | 0 | 2 | 0 |
| ENSGACG00000006638 | ChrI | 3984198 | 4032562 | gramd1bb | 2 | 0 | 0 | 0 | 2 | 0 |
| ENSGACG00000007503 | ChrXVI | 14554631 | 14687685 | lrp1bb | 2 | 0 | 0 | 0 | 2 | 0 |
| ENSGACG00000007773 | ChrX | 11796739 | 11880195 | ptprua | 2 | 1 | 0 | 1 | 1 | 0 |
| ENSGACG00000008092 | ChrX | 12446407 | 12460395 | eepd1 | 2 | 0 | 0 | 0 | 2 | 0 |
| ENSGACG00000008187 | ChrV | 10034443 | 10040862 | | 2 | 0 | 0 | 0 | 2 | 0 |
| ENSGACG00000008829 | ChrVI | 10053208 | 10156710 | kcnma1a | 2 | 0 | 0 | 0 | 2 | 0 |
| ENSGACG00000009156 | ChrXVI | 17703064 | 17704594 | | 2 | 2 | 2 | 0 | 0 | 0 |
| ENSGACG00000010158 | ChrXII | 12648568 | 12734883 | ERC2 (1 of many) | 2 | 2 | 2 | 0 | 0 | 0 |
| ENSGACG00000010489 | ChrVIII | 13247354 | 13252601 | use1 | 2 | 0 | 0 | 0 | 2 | 0 |
| ENSGACG00000010661 | ChrXI | 8377436 | 8379317 | znf750 | 2 | 0 | 0 | 0 | 2 | 0 |
| ENSGACG00000011608 | ChrXVIII | 12075826 | 12137714 | stxbp5a | 2 | 0 | 0 | 0 | 1 | 1 |
| ENSGACG00000011935 | ChrXVIII | 12659540 | 12662849 | r0seh1 | 2 | 0 | 0 | 0 | 2 | 0 |
| ENSGACG00000014285 | ChrII | 2430586 | 2434229 | slc7a6os | 2 | 0 | 0 | 0 | 2 | 0 |
| ENSGACG00000014667 | ChrXIII | 19641934 | 19644791 | coq5 | 2 | 0 | 0 | 0 | 2 | 0 |
| ENSGACG00000016544 | ChrIV | 2203221 | 2209876 | neil3 | 2 | 0 | 0 | 0 | 2 | 0 |
| ENSGACG00000016953 | ChrII | 19113447 | 19146003 | | 2 | 0 | 0 | 0 | 1 | 1 |
| ENSGACG00000017000 | ChrII | 19349048 | 19401226 | myo9aa | 2 | 2 | 0 | 2 | 0 | 0 |
| ENSGACG00000017077 | ChrIV | 5517696 | 5539310 | mrpl11 | 2 | 1 | 1 | 0 | 1 | 0 |
| ENSGACG00000017327 | ChrIII | 13672383 | 13693975 | rock1 | 2 | 0 | 0 | 0 | 1 | 1 |
| ENSGACG00000017644 | ChrIII | 14929551 | 14958963 | | 2 | 0 | 0 | 0 | 2 | 0 |
| ENSGACG00000018082 | ChrXIV | 11064738 | 11154619 | whr0 | 2 | 0 | 0 | 0 | 2 | 0 |

| | | | | | | | | | | |
|---|---|---|---|---|---|---|---|---|---|---|
| ENSGACG00000019224 | ChrIX | 16048022 | 16056648 | cep170ab | 2 | 2 | 0 | 2 | 0 | 0 |
| ENSGACG00000019481 | ChrIV | 25424476 | 25433284 | krr1 | 2 | 0 | 0 | 0 | 1 | 1 |
| ENSGACG00000000075 | scaffold_99 | 201535 | 229070 | | 1 | 0 | 0 | 0 | 1 | 0 |
| ENSGACG00000000307 | scaffold_27 | 48405 | 52111 | | 1 | 0 | 0 | 0 | 1 | 0 |
| ENSGACG00000000381 | scaffold_131 | 232303 | 234144 | rbm46 | 1 | 0 | 0 | 0 | 1 | 0 |
| ENSGACG00000000554 | scaffold_67 | 141528 | 172215 | DPP6 | 1 | 1 | 1 | 0 | 0 | 0 |
| ENSGACG00000000557 | scaffold_27 | 1192061 | 1196661 | | 1 | 0 | 0 | 0 | 0 | 1 |
| ENSGACG00000000582 | scaffold_67 | 364680 | 406885 | CDH18 | 1 | 0 | 0 | 0 | 1 | 0 |
| ENSGACG00000000634 | scaffold_90 | 226420 | 229275 | cradd | 1 | 0 | 0 | 0 | 1 | 0 |
| ENSGACG00000000950 | scaffold_157 | 146047 | 149073 | si:dkey-201i6.2 | 1 | 0 | 0 | 0 | 1 | 0 |
| ENSGACG00000001063 | scaffold_27 | 3053896 | 3067281 | dstyk | 1 | 0 | 0 | 0 | 0 | 1 |
| ENSGACG00000001165 | scaffold_129 | 135238 | 139395 | tfr2 | 1 | 0 | 0 | 0 | 1 | 0 |
| ENSGACG00000001382 | scaffold_27 | 4081612 | 4132724 | cadpsb | 1 | 0 | 0 | 0 | 1 | 0 |
| ENSGACG00000001468 | scaffold_111 | 190733 | 194077 | mgat1b | 1 | 0 | 0 | 0 | 0 | 1 |
| ENSGACG00000001581 | scaffold_120 | 126130 | 128131 | si:ch1073-184j22.2 | 1 | 0 | 0 | 0 | 0 | 1 |
| ENSGACG00000001580 | scaffold_169 | 57700 | 104630 | nrp1b | 1 | 0 | 0 | 0 | 1 | 0 |
| ENSGACG00000001633 | scaffold_120 | 253516 | 254412 | | 1 | 0 | 0 | 0 | 0 | 1 |
| ENSGACG00000001637 | scaffold_106 | 301947 | 326476 | aco1 | 1 | 1 | 0 | 1 | 0 | 0 |
| ENSGACG00000001734 | ChrXVI | 2405778 | 2407622 | kbtbd7 | 1 | 0 | 0 | 0 | 1 | 0 |
| ENSGACG00000001745 | scaffold_122 | 182930 | 185817 | tlr18 | 1 | 0 | 0 | 0 | 0 | 1 |
| ENSGACG00000001883 | ChrXVI | 2628609 | 2645144 | klf12a | 1 | 0 | 0 | 0 | 1 | 0 |
| ENSGACG00000001892 | ChrXVI | 2776645 | 2777646 | gpr18 | 1 | 1 | 0 | 1 | 0 | 0 |
| ENSGACG00000001980 | scaffold_149 | 22071 | 23010 | | 1 | 1 | 0 | 1 | 0 | 0 |
| ENSGACG00000001990 | ChrX | 1252132 | 1253742 | zgc:91910 | 1 | 0 | 0 | 0 | 0 | 1 |
| ENSGACG00000001974 | ChrXXI | 2122593 | 2149047 | prkdc | 1 | 0 | 0 | 0 | 1 | 0 |
| ENSGACG00000002007 | ChrXVI | 3402359 | 3407495 | | 1 | 0 | 0 | 0 | 0 | 1 |
| ENSGACG00000002044 | scaffold_214 | 65170 | 78666 | fam69c | 1 | 0 | 0 | 0 | 1 | 0 |
| ENSGACG00000002216 | ChrXVI | 4428375 | 4457157 | ADARB1 | 1 | 0 | 0 | 0 | 1 | 0 |
| ENSGACG00000002328 | ChrXVI | 5502519 | 5557728 | d0h7 | 1 | 0 | 0 | 0 | 1 | 0 |

| | | | | | | | | | | |
|---|---|---|---|---|---|---|---|---|---|---|
| ENSGACG00000002439 | ChrVI | 624617 | 631220 | degs1 | 1 | 0 | 0 | 0 | 0 | 1 |
| ENSGACG00000002542 | ChrV | 1182211 | 1207003 | CHST15 | 1 | 1 | 1 | 0 | 0 | 0 |
| ENSGACG00000002640 | ChrV | 1488301 | 1595530 | dock1 | 1 | 0 | 0 | 0 | 1 | 0 |
| ENSGACG00000002655 | ChrXII | 409174 | 434561 | | 1 | 0 | 0 | 0 | 1 | 0 |
| ENSGACG00000002755 | ChrXVI | 6491791 | 6504029 | gpr39 | 1 | 0 | 0 | 0 | 1 | 0 |
| ENSGACG00000002766 | ChrXVI | 6578389 | 6579905 | | 1 | 0 | 0 | 0 | 1 | 0 |
| ENSGACG00000002798 | ChrXVI | 6698167 | 6721328 | cmss1 | 1 | 0 | 0 | 0 | 1 | 0 |
| ENSGACG00000002794 | ChrXXI | 6761590 | 6775206 | trpa1b | 1 | 0 | 0 | 0 | 1 | 0 |
| ENSGACG00000002797 | ChrXII | 657639 | 663452 | rbm39b | 1 | 0 | 0 | 0 | 1 | 0 |
| ENSGACG00000002805 | ChrXII | 664631 | 666046 | rh50 | 1 | 0 | 0 | 0 | 1 | 0 |
| ENSGACG00000002820 | ChrVI | 1604348 | 1655313 | INPP5A | 1 | 0 | 0 | 0 | 1 | 0 |
| ENSGACG00000002821 | ChrXVI | 6947968 | 6950857 | r0seh2b | 1 | 0 | 0 | 0 | 1 | 0 |
| ENSGACG00000002889 | ChrXVI | 7108006 | 7109040 | sox1a | 1 | 0 | 0 | 0 | 1 | 0 |
| ENSGACG00000002982 | ChrXII | 1118299 | 1128392 | TMCC1 (1 of many) | 1 | 0 | 0 | 0 | 0 | 1 |
| ENSGACG00000003024 | ChrXII | 1350285 | 1355534 | ddx23 | 1 | 0 | 0 | 0 | 1 | 0 |
| ENSGACG00000003039 | ChrXVII | 106908 | 109258 | | 1 | 0 | 0 | 0 | 1 | 0 |
| ENSGACG00000003210 | ChrXVII | 474804 | 485236 | | 1 | 0 | 0 | 0 | 1 | 0 |
| ENSGACG00000003216 | ChrVI | 2793166 | 2802270 | pik3ap1 | 1 | 0 | 0 | 0 | 1 | 0 |
| ENSGACG00000003244 | ChrX | 4431116 | 4431815 | | 1 | 0 | 0 | 0 | 0 | 1 |
| ENSGACG00000003280 | ChrX | 4476813 | 4485782 | rprd2a | 1 | 0 | 0 | 0 | 1 | 0 |
| ENSGACG00000003349 | ChrXII | 1948134 | 1950471 | | 1 | 0 | 0 | 0 | 0 | 1 |
| ENSGACG00000003467 | ChrXX | 759236 | 760746 | apoa2 | 1 | 0 | 0 | 0 | 0 | 1 |
| ENSGACG00000003513 | ChrXVII | 772640 | 787599 | rims4 | 1 | 1 | 1 | 0 | 0 | 0 |
| ENSGACG00000003533 | ChrX | 4941823 | 4965131 | jarid2b | 1 | 0 | 0 | 0 | 1 | 0 |
| ENSGACG00000003552 | ChrX | 5093819 | 5117753 | pard6gb | 1 | 0 | 0 | 0 | 1 | 0 |
| ENSGACG00000003560 | ChrXIII | 768298 | 822763 | NR6A1 | 1 | 0 | 0 | 0 | 1 | 0 |
| ENSGACG00000003616 | ChrXII | 2350366 | 2351344 | pdyn | 1 | 0 | 0 | 0 | 1 | 0 |
| ENSGACG00000003621 | ChrXXI | 8263488 | 8269613 | gyg1b | 1 | 0 | 0 | 0 | 1 | 0 |

| | | | | | | | | | | |
|---|---|---|---|---|---|---|---|---|---|---|
| ENSGACG00000003642 | ChrXIII | 971041 | 971448 | mrpl41 | 1 | 0 | 0 | 0 | 1 | 0 |
| ENSGACG00000003641 | ChrXVII | 1100095 | 1107153 | ahcy | 1 | 0 | 0 | 0 | 0 | 1 |
| ENSGACG00000003673 | ChrVIII | 1742873 | 1762035 | crocc2 | 1 | 0 | 0 | 0 | 1 | 0 |
| ENSGACG00000003793 | ChrXVII | 1220142 | 1250579 | slmapa | 1 | 0 | 0 | 0 | 1 | 0 |
| ENSGACG00000003830 | ChrXVI | 8743067 | 8776300 | ANOS1 | 1 | 0 | 0 | 0 | 1 | 0 |
| ENSGACG00000003851 | ChrXVI | 8803555 | 8808879 | gpr143 | 1 | 0 | 0 | 0 | 1 | 0 |
| ENSGACG00000003974 | ChrVI | 4172491 | 4201696 | b3gat2 | 1 | 0 | 0 | 0 | 1 | 0 |
| ENSGACG00000004154 | ChrVI | 4697513 | 4739277 | sipa1l2 | 1 | 0 | 0 | 0 | 1 | 0 |
| ENSGACG00000004161 | ChrX | 6444050 | 6447473 | cyp51 | 1 | 0 | 0 | 0 | 0 | 1 |
| ENSGACG00000004237 | ChrXVI | 9502200 | 9533558 | DOCK9 (1 of many) | 1 | 0 | 0 | 0 | 0 | 1 |
| ENSGACG00000004387 | ChrX | 6719206 | 6725533 | znf687b | 1 | 0 | 0 | 0 | 1 | 0 |
| ENSGACG00000004411 | ChrV | 5205599 | 5280399 | snx29 | 1 | 0 | 0 | 0 | 1 | 0 |
| ENSGACG00000004431 | ChrXII | 3958137 | 3978532 | ntsr1 | 1 | 1 | 0 | 1 | 0 | 0 |
| ENSGACG00000004453 | ChrXVIII | 1071040 | 1077095 | si:ch211-63o20.7 | 1 | 0 | 0 | 0 | 1 | 0 |
| ENSGACG00000004448 | ChrXVII | 2225924 | 2233495 | dhrs3b | 1 | 0 | 0 | 0 | 1 | 0 |
| ENSGACG00000004636 | ChrI | 372834 | 374001 | | 1 | 0 | 0 | 0 | 1 | 0 |
| ENSGACG00000004635 | ChrXIII | 2105329 | 2109391 | si:dkey-193c22.1 | 1 | 0 | 0 | 0 | 1 | 0 |
| ENSGACG00000004738 | ChrXVII | 2595419 | 2644876 | | 1 | 0 | 0 | 0 | 1 | 0 |
| ENSGACG00000004752 | ChrXVII | 2653110 | 2664476 | si:ch211-286o17.1 | 1 | 0 | 0 | 0 | 1 | 0 |
| ENSGACG00000004716 | ChrX | 7178557 | 7197519 | si:dkeyp-120h9.1 | 1 | 0 | 0 | 0 | 0 | 1 |
| ENSGACG00000004743 | ChrXX | 2589138 | 2594554 | sqlea | 1 | 0 | 0 | 0 | 1 | 0 |
| ENSGACG00000004766 | ChrXX | 2597737 | 2628658 | oxr1b | 1 | 1 | 1 | 0 | 0 | 0 |
| ENSGACG00000004773 | ChrXII | 4621502 | 4622140 | | 1 | 0 | 0 | 0 | 1 | 0 |
| ENSGACG00000004780 | ChrXII | 4743053 | 4896679 | plx01a | 1 | 1 | 1 | 0 | 0 | 0 |
| ENSGACG00000004808 | ChrXVI | 10104986 | 10129378 | | 1 | 0 | 0 | 0 | 1 | 0 |
| ENSGACG00000004873 | ChrXXI | 10500632 | 10511464 | CDH19 | 1 | 1 | 1 | 0 | 0 | 0 |
| ENSGACG00000004940 | ChrXVIII | 1504942 | 1509134 | serpi01l | 1 | 0 | 0 | 0 | 0 | 1 |
| ENSGACG00000005016 | ChrVIII | 3619176 | 3623161 | | 1 | 0 | 0 | 0 | 1 | 0 |

| | | | | | | | | | | |
|---|---|---|---|---|---|---|---|---|---|---|
| ENSGACG00000005034 | ChrXVII | 3261730 | 3295915 | 0v1b | 1 | 0 | 0 | 0 | 0 | 1 |
| ENSGACG00000005087 | ChrXXI | 10930916 | 10935699 | mastl | 1 | 0 | 0 | 0 | 0 | 1 |
| ENSGACG00000005138 | ChrXI | 818949 | 822342 | | 1 | 0 | 0 | 0 | 1 | 0 |
| ENSGACG00000005222 | ChrXVII | 3437445 | 3438929 | kc02a | 1 | 0 | 0 | 0 | 1 | 0 |
| ENSGACG00000005224 | ChrXVIII | 2150130 | 2155377 | gtf2a1 | 1 | 0 | 0 | 0 | 1 | 0 |
| ENSGACG00000005280 | ChrXVIII | 2211718 | 2416099 | nrxn3b | 1 | 1 | 1 | 0 | 0 | 0 |
| ENSGACG00000005395 | ChrXIII | 3541942 | 3609276 | unc5db | 1 | 0 | 0 | 0 | 1 | 0 |
| ENSGACG00000005473 | ChrXVII | 3704985 | 3708788 | ncoa5 | 1 | 0 | 0 | 0 | 0 | 1 |
| ENSGACG00000005498 | ChrXII | 6282953 | 6382409 | | 1 | 0 | 0 | 0 | 1 | 0 |
| ENSGACG00000005579 | ChrXII | 6456672 | 6459151 | casp9 | 1 | 1 | 1 | 0 | 0 | 0 |
| ENSGACG00000005585 | ChrI | 1594281 | 1664168 | si:ch211-276c2.2 | 1 | 0 | 0 | 0 | 0 | 1 |
| ENSGACG00000005642 | ChrXII | 6730343 | 6739230 | | 1 | 0 | 0 | 0 | 0 | 1 |
| ENSGACG00000005690 | ChrVIII | 5061230 | 5065590 | fetub | 1 | 0 | 0 | 0 | 0 | 1 |
| ENSGACG00000005826 | ChrXV | 2081586 | 2088261 | pomt2 | 1 | 0 | 0 | 0 | 1 | 0 |
| ENSGACG00000005871 | ChrXI | 2407492 | 2415100 | 0t15 | 1 | 1 | 1 | 0 | 0 | 0 |
| ENSGACG00000005912 | ChrXVII | 4162499 | 4184565 | eya2 | 1 | 0 | 0 | 0 | 1 | 0 |
| ENSGACG00000005959 | ChrXIII | 4441375 | 4446248 | CKMT2 | 1 | 0 | 0 | 0 | 1 | 0 |
| ENSGACG00000005959 | ChrXIII | 4441375 | 4446248 | CKMT2 | 1 | 0 | 0 | 0 | 0 | 1 |
| ENSGACG00000006073 | ChrXVI | 11690439 | 11697554 | SMARCAL1 | 1 | 0 | 0 | 0 | 1 | 0 |
| ENSGACG00000006136 | ChrI | 2281697 | 2283639 | | 1 | 0 | 0 | 0 | 1 | 0 |
| ENSGACG00000006109 | ChrXV | 2485491 | 2488657 | ACTA1 (1 of many) | 1 | 0 | 0 | 0 | 1 | 0 |
| ENSGACG00000006168 | ChrXVIII | 3331871 | 3333721 | col10a1b | 1 | 1 | 1 | 0 | 0 | 0 |
| ENSGACG00000006159 | ChrXII | 7573896 | 7590952 | | 1 | 0 | 0 | 0 | 1 | 0 |
| ENSGACG00000006162 | ChrV | 8116939 | 8120965 | | 1 | 0 | 0 | 0 | 0 | 1 |
| ENSGACG00000006227 | ChrI | 2824077 | 2830150 | rap1gap2a | 1 | 0 | 0 | 0 | 1 | 0 |
| ENSGACG00000006220 | ChrXIII | 4822900 | 4865643 | DMXL1 | 1 | 0 | 0 | 0 | 1 | 0 |
| ENSGACG00000006268 | ChrXVIII | 3894826 | 3965753 | epha7 | 1 | 1 | 1 | 0 | 0 | 0 |
| ENSGACG00000006305 | ChrXIII | 4975851 | 4979063 | mapkapk5 | 1 | 0 | 0 | 0 | 1 | 0 |

| | | | | | | | | | | |
|---|---|---|---|---|---|---|---|---|---|---|
| ENSGACG00000006314 | ChrXVI | 12033164 | 12044262 | | 1 | 1 | 1 | 0 | 0 | 0 |
| ENSGACG00000006321 | ChrI | 3060604 | 3065797 | p2rx8 | 1 | 0 | 0 | 0 | 1 | 0 |
| ENSGACG00000006409 | ChrXI | 2907304 | 2924089 | plppr2a | 1 | 1 | 1 | 0 | 0 | 0 |
| ENSGACG00000006410 | ChrXVII | 5112878 | 5123495 | shmt2 | 1 | 0 | 0 | 0 | 1 | 0 |
| ENSGACG00000006443 | ChrXII | 7864313 | 7874314 | ppih | 1 | 0 | 0 | 0 | 1 | 0 |
| ENSGACG00000006475 | ChrXVIII | 4611333 | 4622904 | | 1 | 0 | 0 | 0 | 0 | 1 |
| ENSGACG00000006571 | ChrX | 9279918 | 9287935 | casd1 | 1 | 1 | 0 | 1 | 0 | 0 |
| ENSGACG00000006656 | ChrXV | 2935681 | 2938985 | | 1 | 0 | 0 | 0 | 1 | 0 |
| ENSGACG00000006652 | ChrX | 9301880 | 9334968 | ppp1r9a | 1 | 0 | 0 | 0 | 1 | 0 |
| ENSGACG00000006740 | ChrXI | 3465800 | 3467727 | si:dkeyp-69e1.8 | 1 | 0 | 0 | 0 | 0 | 1 |
| ENSGACG00000006849 | ChrX | 9509713 | 9515132 | mpp6b | 1 | 0 | 0 | 0 | 0 | 1 |
| ENSGACG00000006931 | ChrXV | 3674888 | 3699659 | | 1 | 0 | 0 | 0 | 1 | 0 |
| ENSGACG00000006954 | ChrX | 9600905 | 9605308 | | 1 | 0 | 0 | 0 | 1 | 0 |
| ENSGACG00000007068 | ChrI | 4543685 | 4582641 | KC0B1 | 1 | 0 | 0 | 0 | 1 | 0 |
| ENSGACG00000007130 | ChrXX | 8083957 | 8088889 | cdcp1a | 1 | 0 | 0 | 0 | 1 | 0 |
| ENSGACG00000007220 | ChrV | 9430229 | 9431432 | zgc:113090 | 1 | 0 | 0 | 0 | 0 | 1 |
| ENSGACG00000007241 | ChrXV | 4150772 | 4153424 | fam167b | 1 | 0 | 0 | 0 | 1 | 0 |
| ENSGACG00000007260 | ChrXV | 4158259 | 4161687 | tdh2 | 1 | 0 | 0 | 0 | 1 | 0 |
| ENSGACG00000007146 | ChrXVI | 14092977 | 14114066 | dars | 1 | 0 | 0 | 0 | 1 | 0 |
| ENSGACG00000007270 | ChrVIII | 8208428 | 8335563 | | 1 | 0 | 0 | 0 | 1 | 0 |
| ENSGACG00000007280 | ChrXV | 4171868 | 4184504 | clic4 | 1 | 0 | 0 | 0 | 0 | 1 |
| ENSGACG00000007343 | ChrX | 10666995 | 10679875 | col9a2 | 1 | 0 | 0 | 0 | 0 | 1 |
| ENSGACG00000007372 | ChrX | 10807926 | 10961030 | csmd2 | 1 | 0 | 0 | 0 | 0 | 1 |
| ENSGACG00000007442 | ChrXVII | 6294895 | 6297552 | | 1 | 0 | 0 | 0 | 0 | 1 |
| ENSGACG00000007613 | ChrI | 5854274 | 5864978 | baz1b | 1 | 0 | 0 | 0 | 1 | 0 |
| ENSGACG00000007693 | ChrI | 5937570 | 5939340 | zgc:103681 | 1 | 0 | 0 | 0 | 1 | 0 |
| ENSGACG00000007711 | ChrXX | 9099195 | 9126875 | nphs1 | 1 | 0 | 0 | 0 | 1 | 0 |
| ENSGACG00000007746 | ChrXV | 4848926 | 4850768 | tmed8 | 1 | 0 | 0 | 0 | 0 | 1 |
| ENSGACG00000007745 | ChrXIII | 7495399 | 7509368 | EGFLAM | 1 | 0 | 0 | 0 | 0 | 1 |

| | | | | | | | | | | |
|---|---|---|---|---|---|---|---|---|---|---|
| ENSGACG00000007819 | ChrX | 12069833 | 12099979 | foxo6b | 1 | 1 | 1 | 0 | 0 | 0 |
| ENSGACG00000007825 | ChrI | 6236909 | 6237643 | a0pc15 | 1 | 0 | 0 | 0 | 1 | 0 |
| ENSGACG00000007835 | ChrV | 9774074 | 9779340 | | 1 | 0 | 0 | 0 | 1 | 0 |
| ENSGACG00000007863 | ChrVIII | 8897020 | 8952064 | nek7 | 1 | 0 | 0 | 0 | 1 | 0 |
| ENSGACG00000007953 | ChrI | 6283524 | 6297921 | NOX4 | 1 | 0 | 0 | 0 | 1 | 0 |
| ENSGACG00000008143 | ChrXVI | 15980825 | 15981403 | arl4cb | 1 | 0 | 0 | 0 | 1 | 0 |
| ENSGACG00000008201 | ChrVIII | 9458245 | 9463856 | | 1 | 0 | 0 | 0 | 1 | 0 |
| ENSGACG00000008207 | ChrI | 6847350 | 6887280 | si:ch211-248g20.5 | 1 | 0 | 0 | 0 | 1 | 0 |
| ENSGACG00000008217 | ChrVIII | 9502643 | 9506528 | | 1 | 0 | 0 | 0 | 0 | 1 |
| ENSGACG00000008225 | ChrX | 12619123 | 12643081 | grb10a | 1 | 1 | 1 | 0 | 0 | 0 |
| ENSGACG00000008252 | ChrX | 12695029 | 12695737 | | 1 | 0 | 0 | 0 | 0 | 1 |
| ENSGACG00000008350 | ChrXIII | 8375895 | 8401616 | smtnb | 1 | 0 | 0 | 0 | 1 | 0 |
| ENSGACG00000008540 | ChrXVI | 16966117 | 16967188 | | 1 | 0 | 0 | 0 | 0 | 1 |
| ENSGACG00000008628 | ChrXVII | 7986609 | 8018375 | | 1 | 1 | 1 | 0 | 0 | 0 |
| ENSGACG00000008689 | ChrV | 10760247 | 10779914 | | 1 | 1 | 1 | 0 | 0 | 0 |
| ENSGACG00000008798 | ChrX | 13132413 | 13138599 | si:ch1073-296d18.1 | 1 | 0 | 0 | 0 | 0 | 1 |
| ENSGACG00000009135 | ChrVI | 10670867 | 10727803 | pcdh15a | 1 | 0 | 0 | 0 | 0 | 1 |
| ENSGACG00000009146 | ChrXIII | 9082014 | 9083453 | fam222a | 1 | 1 | 1 | 0 | 0 | 0 |
| ENSGACG00000009211 | ChrXVIII | 8793962 | 8796629 | si:dkey-65b13.13 | 1 | 0 | 0 | 0 | 1 | 0 |
| ENSGACG00000009251 | ChrVIII | 11476798 | 11480609 | htr2b | 1 | 0 | 0 | 0 | 1 | 0 |
| ENSGACG00000009248 | ChrXVIII | 8940535 | 8942987 | | 1 | 0 | 0 | 0 | 1 | 0 |
| ENSGACG00000009346 | ChrVIII | 11764463 | 11771444 | srsf11 | 1 | 1 | 1 | 0 | 0 | 0 |
| ENSGACG00000009393 | ChrXVII | 9412190 | 9495385 | FAM19A1 | 1 | 0 | 0 | 0 | 1 | 0 |
| ENSGACG00000009627 | ChrXX | 11403589 | 11405982 | si:ch73-54f23.4 | 1 | 0 | 0 | 0 | 1 | 0 |
| ENSGACG00000009653 | ChrX | 14747980 | 14771097 | znf976 | 1 | 1 | 0 | 1 | 0 | 0 |
| ENSGACG00000009650 | ChrXV | 7669664 | 7725769 | qkia | 1 | 0 | 0 | 0 | 1 | 0 |
| ENSGACG00000009733 | ChrXIII | 9474478 | 9479229 | D0JB5 | 1 | 1 | 0 | 1 | 0 | 0 |
| ENSGACG00000009746 | ChrX | 14996770 | 15014430 | | 1 | 0 | 0 | 0 | 1 | 0 |

| | | | | | | | | | | |
|---|---|---|---|---|---|---|---|---|---|---|
| ENSGACG00000009796 | ChrXVIII | 9441961 | 9443603 | kcns3b | 1 | 0 | 0 | 0 | 1 | 0 |
| ENSGACG00000009870 | ChrXIII | 9641184 | 9654500 | serinc5 | 1 | 0 | 0 | 0 | 1 | 0 |
| ENSGACG00000009920 | ChrXV | 8067468 | 8068457 | | 1 | 0 | 0 | 0 | 0 | 1 |
| ENSGACG00000009923 | ChrX | 15274191 | 15282234 | | 1 | 0 | 0 | 0 | 1 | 0 |
| ENSGACG00000009968 | ChrXV | 8210188 | 8222680 | | 1 | 0 | 0 | 0 | 1 | 0 |
| ENSGACG00000009983 | ChrXVII | 10033214 | 10065567 | cdh4 | 1 | 0 | 0 | 0 | 0 | 1 |
| ENSGACG00000009991 | ChrXVIII | 9665614 | 9671172 | tagapb | 1 | 0 | 0 | 0 | 0 | 1 |
| ENSGACG00000010108 | ChrVI | 12267753 | 12268331 | grem2a | 1 | 0 | 0 | 0 | 1 | 0 |
| ENSGACG00000010176 | ChrXIII | 9941933 | 9943709 | | 1 | 0 | 0 | 0 | 1 | 0 |
| ENSGACG00000010260 | ChrXV | 8712213 | 8715682 | manea | 1 | 0 | 0 | 0 | 1 | 0 |
| ENSGACG00000010297 | ChrVI | 12537633 | 12538976 | | 1 | 0 | 0 | 0 | 0 | 1 |
| ENSGACG00000010299 | ChrXVIII | 10233449 | 10270176 | daam1b | 1 | 0 | 0 | 0 | 1 | 0 |
| ENSGACG00000010494 | ChrVIII | 13255573 | 13285202 | si:zfos-588f8.1 | 1 | 0 | 0 | 0 | 1 | 0 |
| ENSGACG00000010507 | ChrXV | 9224970 | 9273325 | zgc:154061 | 1 | 0 | 0 | 0 | 1 | 0 |
| ENSGACG00000010532 | ChrXV | 9468781 | 9481444 | spred1 | 1 | 0 | 0 | 0 | 1 | 0 |
| ENSGACG00000010542 | ChrXV | 9490435 | 9501192 | ptpn21 | 1 | 0 | 0 | 0 | 1 | 0 |
| ENSGACG00000010528 | ChrVIII | 13335579 | 13342295 | ncln | 1 | 0 | 0 | 0 | 1 | 0 |
| ENSGACG00000010643 | ChrXVII | 10920133 | 10925973 | | 1 | 0 | 0 | 0 | 0 | 1 |
| ENSGACG00000010835 | ChrXV | 9862819 | 9867105 | slc35g2a | 1 | 0 | 0 | 0 | 0 | 1 |
| ENSGACG00000010886 | ChrXIII | 12045266 | 12049108 | si:dkey-32e23.6 | 1 | 0 | 0 | 0 | 1 | 0 |
| ENSGACG00000010951 | ChrI | 12038391 | 12056617 | si:dkey-243i1.1 | 1 | 0 | 0 | 0 | 1 | 0 |
| ENSGACG00000011003 | ChrVI | 14099672 | 14126488 | kif20ba | 1 | 0 | 0 | 0 | 1 | 0 |
| ENSGACG00000011145 | ChrVIII | 13948367 | 13950370 | dmrta2 | 1 | 1 | 1 | 0 | 0 | 0 |
| ENSGACG00000011191 | ChrVI | 14358268 | 14389292 | slc30a6 | 1 | 0 | 0 | 0 | 1 | 0 |
| ENSGACG00000011235 | ChrXV | 10608343 | 10624532 | crip2 | 1 | 0 | 0 | 0 | 1 | 0 |
| ENSGACG00000011269 | ChrXV | 10677665 | 10678117 | tmem229b | 1 | 1 | 1 | 0 | 0 | 0 |
| ENSGACG00000011225 | ChrVI | 14477471 | 14520782 | SPTBN1 | 1 | 0 | 0 | 0 | 1 | 0 |
| ENSGACG00000011313 | ChrXI | 9901388 | 9903421 | vasnb | 1 | 0 | 0 | 0 | 1 | 0 |
| ENSGACG00000011355 | ChrXVIII | 11589919 | 11605386 | nhsl1b | 1 | 0 | 0 | 0 | 1 | 0 |

| | | | | | | | | | | |
|---|---|---|---|---|---|---|---|---|---|---|
| ENSGACG00000011353 | ChrXII | 15159858 | 15171195 | mical1 | 1 | 0 | 0 | 0 | 0 | 1 |
| ENSGACG00000011470 | ChrVIII | 14490182 | 14491442 | | 1 | 0 | 0 | 0 | 1 | 0 |
| ENSGACG00000011556 | ChrXIII | 13107695 | 13111326 | rorb | 1 | 1 | 1 | 0 | 0 | 0 |
| ENSGACG00000011602 | ChrXVII | 12652011 | 12662065 | LAMB3 | 1 | 0 | 0 | 0 | 1 | 0 |
| ENSGACG00000011628 | ChrVI | 15521491 | 15543739 | nckap1 | 1 | 0 | 0 | 0 | 1 | 0 |
| ENSGACG00000011643 | ChrXI | 10486001 | 10490341 | cacng5b | 1 | 0 | 0 | 0 | 1 | 0 |
| ENSGACG00000011670 | ChrXX | 12649677 | 12655617 | REC8 | 1 | 1 | 1 | 0 | 0 | 0 |
| ENSGACG00000011679 | ChrI | 13725256 | 13739469 | trim3a | 1 | 0 | 0 | 0 | 0 | 1 |
| ENSGACG00000011694 | ChrXI | 10642924 | 10653747 | axin2 | 1 | 0 | 0 | 0 | 1 | 0 |
| ENSGACG00000011697 | ChrI | 13774012 | 13834603 | dchs1a | 1 | 0 | 0 | 0 | 1 | 0 |
| ENSGACG00000011710 | ChrXV | 11490059 | 11493449 | | 1 | 0 | 0 | 0 | 0 | 1 |
| ENSGACG00000011711 | ChrVIII | 15092507 | 15103135 | olfml2ba | 1 | 0 | 0 | 0 | 0 | 1 |
| ENSGACG00000011770 | ChrXVII | 13695917 | 13762426 | pdzrn3b | 1 | 0 | 0 | 0 | 0 | 1 |
| ENSGACG00000011933 | ChrVI | 16265135 | 16268160 | tmem41ab | 1 | 0 | 0 | 0 | 0 | 1 |
| ENSGACG00000012010 | ChrXVIII | 12857870 | 12861754 | | 1 | 0 | 0 | 0 | 0 | 1 |
| ENSGACG00000012063 | ChrXV | 11951816 | 11991325 | actn1 | 1 | 0 | 0 | 0 | 1 | 0 |
| ENSGACG00000012128 | ChrI | 14736944 | 14767457 | | 1 | 0 | 0 | 0 | 1 | 0 |
| ENSGACG00000012153 | ChrXIII | 13909513 | 13913900 | asb6 | 1 | 0 | 0 | 0 | 1 | 0 |
| ENSGACG00000012241 | ChrXVIII | 13379496 | 13382944 | rad51 | 1 | 0 | 0 | 0 | 0 | 1 |
| ENSGACG00000012270 | ChrXVIII | 13405473 | 13406295 | | 1 | 0 | 0 | 0 | 1 | 0 |
| ENSGACG00000012283 | ChrI | 15424209 | 15453953 | cdon | 1 | 0 | 0 | 0 | 1 | 0 |
| ENSGACG00000012322 | ChrXVIII | 13649915 | 13680123 | runx2b | 1 | 1 | 0 | 1 | 0 | 0 |
| ENSGACG00000012330 | ChrVIII | 15635360 | 15643570 | rgl1 | 1 | 0 | 0 | 0 | 0 | 1 |
| ENSGACG00000012385 | ChrXVIII | 13772879 | 13784829 | mfsd2b | 1 | 0 | 0 | 0 | 1 | 0 |
| ENSGACG00000012455 | ChrXI | 11660812 | 11665052 | | 1 | 0 | 0 | 0 | 1 | 0 |
| ENSGACG00000012442 | ChrVIII | 15749226 | 15764692 | notch2 | 1 | 0 | 0 | 0 | 1 | 0 |
| ENSGACG00000012512 | ChrXIII | 14598285 | 14599898 | lrrtm4l2 | 1 | 1 | 0 | 1 | 0 | 0 |
| ENSGACG00000012503 | ChrXVIII | 14203391 | 14223968 | PLCB4 | 1 | 0 | 0 | 0 | 1 | 0 |
| ENSGACG00000012587 | ChrXI | 11922124 | 11924367 | wfikkn2a | 1 | 0 | 0 | 0 | 1 | 0 |

| | | | | | | | | | | |
|---|---|---|---|---|---|---|---|---|---|---|
| ENSGACG00000012609 | ChrXI | 12065873 | 12068920 | | 1 | 0 | 0 | 0 | 1 | 0 |
| ENSGACG00000012641 | ChrXX | 14272032 | 14305104 | | 1 | 0 | 0 | 0 | 0 | 1 |
| ENSGACG00000012641 | ChrXX | 14272032 | 14305104 | | 1 | 0 | 0 | 0 | 1 | 0 |
| ENSGACG00000012655 | scaffold_98 | 371516 | 372691 | | 1 | 0 | 0 | 0 | 1 | 0 |
| ENSGACG00000012760 | ChrVIII | 16214608 | 16225148 | si:dkey-110c1.10 | 1 | 1 | 1 | 0 | 0 | 0 |
| ENSGACG00000012876 | ChrVIII | 16469522 | 16475075 | rgmd | 1 | 0 | 0 | 0 | 1 | 0 |
| ENSGACG00000012938 | ChrXVIII | 14983710 | 14985734 | chac1 | 1 | 0 | 0 | 0 | 0 | 1 |
| ENSGACG00000012995 | ChrVIII | 16649364 | 16651755 | fstl3 | 1 | 0 | 0 | 0 | 1 | 0 |
| ENSGACG00000012997 | ChrXII | 17297684 | 17327439 | plch2a | 1 | 0 | 0 | 0 | 1 | 0 |
| ENSGACG00000013040 | ChrXVIII | 15062591 | 15066857 | stx7l | 1 | 0 | 0 | 0 | 0 | 1 |
| ENSGACG00000013071 | ChrVIII | 16932446 | 16936340 | tmem161a | 1 | 0 | 0 | 0 | 1 | 0 |
| ENSGACG00000013118 | ChrXV | 15401988 | 15432525 | BCL11B | 1 | 0 | 0 | 0 | 1 | 0 |
| ENSGACG00000013228 | ChrI | 17134277 | 17138229 | | 1 | 0 | 0 | 0 | 0 | 1 |
| ENSGACG00000013248 | ChrXVIII | 15399767 | 15421191 | nkain2 | 1 | 0 | 0 | 0 | 1 | 0 |
| ENSGACG00000013252 | ChrXVIII | 15429123 | 15433946 | rnf217 | 1 | 1 | 0 | 1 | 0 | 0 |
| ENSGACG00000013255 | ChrXV | 15759319 | 15781086 | MYT1L | 1 | 0 | 0 | 0 | 1 | 0 |
| ENSGACG00000013218 | ChrXI | 12798318 | 12805251 | | 1 | 0 | 0 | 0 | 1 | 0 |
| ENSGACG00000013273 | ChrXIII | 16131688 | 16140819 | arhgap25 | 1 | 0 | 0 | 0 | 1 | 0 |
| ENSGACG00000013297 | ChrVIII | 17242910 | 17245367 | rab11ba | 1 | 1 | 1 | 0 | 0 | 0 |
| ENSGACG00000013422 | scaffold_61 | 545965 | 563244 | fancl | 1 | 0 | 0 | 0 | 1 | 0 |
| ENSGACG00000013463 | ChrXIII | 16423554 | 16426950 | rad9b | 1 | 0 | 0 | 0 | 1 | 0 |
| ENSGACG00000013478 | scaffold_196 | 78142 | 81595 | | 1 | 0 | 0 | 0 | 0 | 1 |
| ENSGACG00000013488 | ChrXI | 13112136 | 13116786 | lrrc4ba | 1 | 0 | 0 | 0 | 0 | 1 |
| ENSGACG00000013504 | ChrXX | 16295857 | 16300154 | hormad1 | 1 | 1 | 1 | 0 | 0 | 0 |
| ENSGACG00000013506 | ChrXI | 13157415 | 13172421 | syt3 | 1 | 0 | 0 | 0 | 1 | 0 |
| ENSGACG00000013523 | ChrIII | 1578911 | 1598618 | | 1 | 1 | 1 | 0 | 0 | 0 |
| ENSGACG00000013636 | ChrVIII | 17679353 | 17685427 | zbtb11 | 1 | 0 | 0 | 0 | 1 | 0 |
| ENSGACG00000013639 | ChrXX | 16926113 | 16927065 | fabp10a | 1 | 0 | 0 | 0 | 0 | 1 |
| ENSGACG00000013787 | ChrXI | 13506470 | 13516007 | tyk2 | 1 | 0 | 0 | 0 | 0 | 1 |

| | | | | | | | | | | |
|---|---|---|---|---|---|---|---|---|---|---|
| ENSGACG00000013869 | ChrXI | 13640461 | 13641149 | | 1 | 0 | 0 | 0 | 0 | 1 |
| ENSGACG00000013977 | ChrVIII | 18160785 | 18167767 | SCML2 (1 of many) | 1 | 0 | 0 | 0 | 0 | 1 |
| ENSGACG00000014073 | ChrXIII | 18076604 | 18087798 | nf2a | 1 | 0 | 0 | 0 | 1 | 0 |
| ENSGACG00000014104 | ChrVIII | 18357770 | 18360435 | cbr4 | 1 | 0 | 0 | 0 | 1 | 0 |
| ENSGACG00000014252 | ChrXIII | 18343004 | 18348481 | suds3 | 1 | 1 | 1 | 0 | 0 | 0 |
| ENSGACG00000014384 | ChrXI | 14979751 | 14982832 | | 1 | 0 | 0 | 0 | 1 | 0 |
| ENSGACG00000014435 | ChrIII | 4386571 | 4396709 | tgm1l1 | 1 | 0 | 0 | 0 | 1 | 0 |
| ENSGACG00000014548 | scaffold_240 | 819 | 11444 | dirc2 | 1 | 1 | 1 | 0 | 0 | 0 |
| ENSGACG00000014576 | ChrI | 22242655 | 22244658 | | 1 | 0 | 0 | 0 | 1 | 0 |
| ENSGACG00000014580 | ChrI | 22269225 | 22284819 | GPM6B (1 of many) | 1 | 0 | 0 | 0 | 1 | 0 |
| ENSGACG00000014627 | ChrI | 22403687 | 22410614 | cbsa | 1 | 0 | 0 | 0 | 1 | 0 |
| ENSGACG00000014832 | ChrII | 5125390 | 5127551 | gnrhr2 | 1 | 0 | 0 | 0 | 1 | 0 |
| ENSGACG00000014892 | ChrIII | 6063746 | 6077226 | | 1 | 0 | 0 | 0 | 1 | 0 |
| ENSGACG00000014902 | ChrII | 5427953 | 5521857 | itfg1 | 1 | 0 | 0 | 0 | 1 | 0 |
| ENSGACG00000014926 | ChrI | 23288291 | 23341962 | zranb3 | 1 | 0 | 0 | 0 | 1 | 0 |
| ENSGACG00000014935 | ChrXI | 16311211 | 16319300 | | 1 | 1 | 0 | 1 | 0 | 0 |
| ENSGACG00000015001 | ChrXI | 16612089 | 16615925 | sgf29 | 1 | 0 | 0 | 0 | 0 | 1 |
| ENSGACG00000015011 | ChrXI | 16647398 | 16653948 | | 1 | 0 | 0 | 0 | 0 | 1 |
| ENSGACG00000015054 | ChrI | 24715179 | 24716868 | | 1 | 0 | 0 | 0 | 1 | 0 |
| ENSGACG00000015079 | ChrI | 25173862 | 25177412 | cyyr1 | 1 | 0 | 0 | 0 | 0 | 1 |
| ENSGACG00000015135 | ChrI | 25487311 | 25491104 | prkag3a | 1 | 1 | 1 | 0 | 0 | 0 |
| ENSGACG00000015240 | ChrI | 26613808 | 26622627 | adarb1a | 1 | 0 | 0 | 0 | 1 | 0 |
| ENSGACG00000015269 | scaffold_48 | 760666 | 799826 | myo15b | 1 | 0 | 0 | 0 | 1 | 0 |
| ENSGACG00000015412 | scaffold_48 | 1674248 | 1685118 | ipmkb | 1 | 0 | 0 | 0 | 1 | 0 |
| ENSGACG00000015490 | ChrXIV | 174415 | 177418 | ufc1 | 1 | 0 | 0 | 0 | 1 | 0 |
| ENSGACG00000015736 | ChrXIV | 723869 | 725546 | mzt2b | 1 | 0 | 0 | 0 | 0 | 1 |
| ENSGACG00000015791 | ChrII | 11127315 | 11148839 | PHLPP2 | 1 | 1 | 1 | 0 | 0 | 0 |
| ENSGACG00000015813 | ChrXIV | 1195794 | 1214842 | | 1 | 0 | 0 | 0 | 0 | 1 |

| | | | | | | | | | | |
|---|---|---|---|---|---|---|---|---|---|---|
| ENSGACG00000015831 | ChrII | 11422567 | 11478619 | tcf12 | 1 | 0 | 0 | 0 | 1 | 0 |
| ENSGACG00000015862 | ChrXIV | 1681568 | 1684183 | npffr2b | 1 | 0 | 0 | 0 | 0 | 1 |
| ENSGACG00000015920 | ChrXIV | 2455047 | 2525326 | lamc3 | 1 | 0 | 0 | 0 | 1 | 0 |
| ENSGACG00000016065 | ChrIII | 9230992 | 9232131 | b3gnt5b | 1 | 0 | 0 | 0 | 0 | 1 |
| ENSGACG00000016073 | ChrIX | 1334639 | 1357056 | | 1 | 1 | 1 | 0 | 0 | 0 |
| ENSGACG00000016204 | ChrIX | 1923297 | 1930876 | ncapg | 1 | 0 | 0 | 0 | 1 | 0 |
| ENSGACG00000016262 | ChrII | 14069329 | 14225725 | ush2a | 1 | 0 | 0 | 0 | 1 | 0 |
| ENSGACG00000016328 | ChrIX | 3038147 | 3051468 | | 1 | 0 | 0 | 0 | 1 | 0 |
| ENSGACG00000016451 | ChrII | 15348070 | 15353934 | CTSH | 1 | 0 | 0 | 0 | 0 | 1 |
| ENSGACG00000016473 | ChrXIV | 3993506 | 3998272 | prlra | 1 | 0 | 0 | 0 | 1 | 0 |
| ENSGACG00000016484 | ChrII | 15471688 | 15483464 | | 1 | 0 | 0 | 0 | 1 | 0 |
| ENSGACG00000016513 | ChrIV | 2044920 | 2048363 | tdo2a | 1 | 0 | 0 | 0 | 1 | 0 |
| ENSGACG00000016529 | ChrIII | 10691525 | 10715730 | st6gal0c3 | 1 | 0 | 0 | 0 | 0 | 1 |
| ENSGACG00000016530 | ChrIX | 4476469 | 4478563 | rdh8b | 1 | 1 | 0 | 1 | 0 | 0 |
| ENSGACG00000016539 | ChrII | 15944730 | 15950252 | scamp5b | 1 | 0 | 0 | 0 | 1 | 0 |
| ENSGACG00000016553 | ChrII | 16198253 | 16219272 | SHF | 1 | 0 | 0 | 0 | 1 | 0 |
| ENSGACG00000016589 | ChrIX | 4587599 | 4591165 | | 1 | 0 | 0 | 0 | 0 | 1 |
| ENSGACG00000016634 | ChrIV | 2582164 | 2682609 | FSTL5 | 1 | 0 | 0 | 0 | 1 | 0 |
| ENSGACG00000016731 | ChrIV | 3340848 | 3411728 | SPATA5 | 1 | 0 | 0 | 0 | 0 | 1 |
| ENSGACG00000016800 | ChrIX | 5319576 | 5323725 | pdia2 | 1 | 0 | 0 | 0 | 1 | 0 |
| ENSGACG00000016846 | ChrIX | 5456262 | 5476760 | | 1 | 1 | 1 | 0 | 0 | 0 |
| ENSGACG00000016849 | ChrXIV | 6012778 | 6022500 | | 1 | 0 | 0 | 0 | 0 | 1 |
| ENSGACG00000016943 | ChrXIV | 6450832 | 6461836 | cabp7b | 1 | 0 | 0 | 0 | 1 | 0 |
| ENSGACG00000016997 | ChrII | 19330861 | 19331520 | senp8 | 1 | 0 | 0 | 0 | 1 | 0 |
| ENSGACG00000017036 | ChrXIV | 6661932 | 6668110 | rnf34b | 1 | 0 | 0 | 0 | 1 | 0 |
| ENSGACG00000017075 | ChrII | 19771601 | 19780222 | zdhhc13 | 1 | 0 | 0 | 0 | 1 | 0 |
| ENSGACG00000017087 | ChrII | 19884513 | 19943913 | 0v2a | 1 | 0 | 0 | 0 | 1 | 0 |
| ENSGACG00000017093 | ChrXIV | 6818762 | 6821900 | trim69 | 1 | 0 | 0 | 0 | 0 | 1 |
| ENSGACG00000017160 | ChrII | 21080854 | 21084821 | | 1 | 0 | 0 | 0 | 1 | 0 |

| | | | | | | | | | | |
|---|---|---|---|---|---|---|---|---|---|---|
| ENSGACG00000017179 | ChrIII | 13106071 | 13116555 | fyco1b | 1 | 0 | 0 | 0 | 1 | 0 |
| ENSGACG00000017190 | ChrIV | 5872021 | 5884796 | mcf2a | 1 | 0 | 0 | 0 | 1 | 0 |
| ENSGACG00000017209 | ChrXIV | 6969603 | 6972066 | pcyox1 | 1 | 0 | 0 | 0 | 1 | 0 |
| ENSGACG00000017250 | ChrIX | 7131393 | 7145303 | gab1 | 1 | 0 | 0 | 0 | 1 | 0 |
| ENSGACG00000017287 | ChrIII | 13454527 | 13465167 | mmp16b | 1 | 1 | 1 | 0 | 0 | 0 |
| ENSGACG00000017291 | ChrIX | 7339426 | 7343930 | elmod2 | 1 | 0 | 0 | 0 | 1 | 0 |
| ENSGACG00000017300 | ChrIII | 13558657 | 13562089 | gata6 | 1 | 0 | 0 | 0 | 1 | 0 |
| ENSGACG00000017313 | ChrII | 21709930 | 21734771 | MEGF11 (1 of many) | 1 | 1 | 1 | 0 | 0 | 0 |
| ENSGACG00000017331 | ChrII | 21831393 | 21842872 | samd4a | 1 | 0 | 0 | 0 | 1 | 0 |
| ENSGACG00000017352 | ChrII | 21934736 | 21940961 | | 1 | 0 | 0 | 0 | 1 | 0 |
| ENSGACG00000017402 | ChrIII | 13870170 | 13884783 | pth1ra | 1 | 0 | 0 | 0 | 1 | 0 |
| ENSGACG00000017480 | ChrIII | 14138063 | 14147391 | WDR37 (1 of many) | 1 | 0 | 0 | 0 | 1 | 0 |
| ENSGACG00000017537 | ChrXIV | 8099548 | 8156144 | bcr | 1 | 0 | 0 | 0 | 1 | 0 |
| ENSGACG00000017598 | ChrXIV | 8297103 | 8367291 | si:dkey-112m2.1 | 1 | 0 | 0 | 0 | 1 | 0 |
| ENSGACG00000017645 | ChrII | 23237993 | 23278096 | vps13c | 1 | 0 | 0 | 0 | 1 | 0 |
| ENSGACG00000017680 | ChrXIV | 8907355 | 8911332 | stx2a | 1 | 0 | 0 | 0 | 1 | 0 |
| ENSGACG00000017696 | ChrIII | 15339421 | 15358805 | | 1 | 0 | 0 | 0 | 1 | 0 |
| ENSGACG00000017662 | ChrIX | 8582121 | 8588508 | acsl1a | 1 | 0 | 0 | 0 | 1 | 0 |
| ENSGACG00000017748 | scaffold_137 | 157404 | 162580 | plekhf2 | 1 | 0 | 0 | 0 | 1 | 0 |
| ENSGACG00000017791 | ChrXIV | 9332963 | 9357146 | | 1 | 0 | 0 | 0 | 1 | 0 |
| ENSGACG00000017788 | ChrIV | 9240071 | 9244174 | p2rx3a | 1 | 0 | 0 | 0 | 1 | 0 |
| ENSGACG00000017924 | ChrXIV | 10388550 | 10398180 | ghrb | 1 | 0 | 0 | 0 | 1 | 0 |
| ENSGACG00000018029 | ChrIV | 11001491 | 11005265 | myoz3a | 1 | 0 | 0 | 0 | 0 | 1 |
| ENSGACG00000018030 | ChrIV | 11007288 | 11014249 | | 1 | 0 | 0 | 0 | 1 | 0 |
| ENSGACG00000018174 | ChrXIV | 11642479 | 11658752 | | 1 | 0 | 0 | 0 | 1 | 0 |
| ENSGACG00000018228 | ChrIX | 10649611 | 10672955 | | 1 | 0 | 0 | 0 | 1 | 0 |
| ENSGACG00000018233 | ChrIX | 10713754 | 10725152 | pdcd11 | 1 | 1 | 1 | 0 | 0 | 0 |
| ENSGACG00000018231 | ChrIV | 12013740 | 12034920 | abcb7 | 1 | 1 | 0 | 1 | 0 | 0 |

| | | | | | | | | | | |
|---|---|---|---|---|---|---|---|---|---|---|
| ENSGACG00000018319 | ChrIV | 12891471 | 12904168 | ergic1 | 1 | 0 | 0 | 0 | 1 | 0 |
| ENSGACG00000018371 | ChrXIV | 14084346 | 14091899 | thap1 | 1 | 0 | 0 | 0 | 1 | 0 |
| ENSGACG00000018389 | ChrIV | 13479246 | 13481484 | | 1 | 1 | 1 | 0 | 0 | 0 |
| ENSGACG00000018400 | ChrIX | 11461137 | 11465372 | | 1 | 0 | 0 | 0 | 0 | 1 |
| ENSGACG00000018496 | ChrIV | 15032996 | 15039498 | si:ch211-160j14,2 | 1 | 0 | 0 | 0 | 0 | 1 |
| ENSGACG00000018568 | ChrIX | 13057908 | 13059965 | si:dkeyp-110c7.8 | 1 | 0 | 0 | 0 | 0 | 1 |
| ENSGACG00000018572 | ChrIV | 15735707 | 15737284 | p2ry10 | 1 | 0 | 0 | 0 | 1 | 0 |
| ENSGACG00000018663 | ChrIV | 16118651 | 16126776 | tmem173 | 1 | 0 | 0 | 0 | 1 | 0 |
| ENSGACG00000018722 | scaffold_208 | 83156 | 90702 | iffo1a | 1 | 1 | 1 | 0 | 0 | 0 |
| ENSGACG00000018779 | ChrVII | 1036400 | 1060359 | col4a6 | 1 | 0 | 0 | 0 | 1 | 0 |
| ENSGACG00000018798 | ChrIX | 14552729 | 14553871 | | 1 | 0 | 0 | 0 | 0 | 1 |
| ENSGACG00000018802 | ChrVII | 1133367 | 1137045 | | 1 | 0 | 0 | 0 | 1 | 0 |
| ENSGACG00000018819 | ChrVII | 1327429 | 1330995 | | 1 | 1 | 1 | 0 | 0 | 0 |
| ENSGACG00000018862 | ChrIV | 18656262 | 18660392 | gxylt1b | 1 | 0 | 0 | 0 | 1 | 0 |
| ENSGACG00000018867 | ChrVII | 1452630 | 1459173 | khnyn | 1 | 0 | 0 | 0 | 0 | 1 |
| ENSGACG00000018927 | ChrIV | 19504962 | 19515715 | | 1 | 0 | 0 | 0 | 0 | 1 |
| ENSGACG00000018991 | ChrVII | 2259206 | 2267772 | zgc:55262 | 1 | 1 | 1 | 0 | 0 | 0 |
| ENSGACG00000019043 | ChrIX | 15241216 | 15246931 | pycr1b | 1 | 0 | 0 | 0 | 1 | 0 |
| ENSGACG00000019057 | ChrVII | 2467974 | 2472162 | mfap3l | 1 | 0 | 0 | 0 | 1 | 0 |
| ENSGACG00000019125 | ChrIV | 21818906 | 21878888 | SYN3 | 1 | 0 | 0 | 0 | 1 | 0 |
| ENSGACG00000019166 | ChrVII | 2903189 | 2904509 | | 1 | 0 | 0 | 0 | 1 | 0 |
| ENSGACG00000019183 | ChrIV | 22236334 | 22237384 | | 1 | 0 | 0 | 0 | 1 | 0 |
| ENSGACG00000019227 | ChrIV | 22899577 | 22913555 | mkrn1 | 1 | 0 | 0 | 0 | 0 | 1 |
| ENSGACG00000019253 | ChrIV | 23209606 | 23250489 | | 1 | 1 | 1 | 0 | 0 | 0 |
| ENSGACG00000019295 | ChrVII | 3222487 | 3232052 | tbc1d19 | 1 | 0 | 0 | 0 | 0 | 1 |
| ENSGACG00000019316 | ChrVII | 4025500 | 4027085 | cnpy4 | 1 | 0 | 0 | 0 | 1 | 0 |
| ENSGACG00000019362 | ChrVII | 4248635 | 4260779 | neurl4 | 1 | 0 | 0 | 0 | 1 | 0 |
| ENSGACG00000019440 | ChrIV | 24711763 | 24712875 | | 1 | 0 | 0 | 0 | 1 | 0 |
| ENSGACG00000019443 | ChrVII | 4842549 | 4846809 | mfsd8 | 1 | 0 | 0 | 0 | 1 | 0 |

| | | | | | | | | | | |
|---|---|---|---|---|---|---|---|---|---|---|
| ENSGACG00000019462 | ChrIX | 17689001 | 17750587 | | 1 | 0 | 0 | 0 | 1 | 0 |
| ENSGACG00000019485 | ChrIV | 25436299 | 25450435 | cmah | 1 | 0 | 0 | 0 | 1 | 0 |
| ENSGACG00000019509 | ChrIV | 25585353 | 25601311 | ANO4 | 1 | 0 | 0 | 0 | 1 | 0 |
| ENSGACG00000019513 | ChrIX | 17973555 | 17980601 | | 1 | 1 | 1 | 0 | 0 | 0 |
| ENSGACG00000019526 | ChrIX | 18022842 | 18029582 | | 1 | 1 | 1 | 0 | 0 | 0 |
| ENSGACG00000019550 | ChrIX | 18092917 | 18100457 | | 1 | 1 | 1 | 0 | 0 | 0 |
| ENSGACG00000019679 | ChrIV | 28725325 | 28735256 | | 1 | 0 | 0 | 0 | 1 | 0 |
| ENSGACG00000019704 | ChrIX | 18828407 | 18830791 | btbd3a | 1 | 1 | 0 | 1 | 0 | 0 |
| ENSGACG00000019718 | ChrIV | 29140176 | 29301388 | plx04 | 1 | 0 | 0 | 0 | 1 | 0 |
| ENSGACG00000019746 | ChrIV | 29700807 | 29758980 | iqsec3a | 1 | 0 | 0 | 0 | 0 | 1 |
| ENSGACG00000019759 | ChrIV | 29797492 | 29817516 | tmtc2a | 1 | 0 | 0 | 0 | 0 | 1 |
| ENSGACG00000019772 | ChrIV | 29858075 | 29865955 | PRMT8 (1 of many) | 1 | 1 | 1 | 0 | 0 | 0 |
| ENSGACG00000019855 | ChrIV | 30901709 | 30906129 | arfgap3 | 1 | 0 | 0 | 0 | 0 | 1 |
| ENSGACG00000019956 | ChrIV | 31331071 | 31344595 | sema3c | 1 | 0 | 0 | 0 | 1 | 0 |
| ENSGACG00000020002 | ChrIX | 20150078 | 20153043 | | 1 | 0 | 0 | 0 | 1 | 0 |
| ENSGACG00000020034 | ChrIV | 31999298 | 32000748 | tnnt2c | 1 | 1 | 1 | 0 | 0 | 0 |
| ENSGACG00000020043 | ChrVII | 10180846 | 10198706 | | 1 | 0 | 0 | 0 | 1 | 0 |
| ENSGACG00000020068 | ChrIV | 32329916 | 32332578 | ERI1 | 1 | 0 | 0 | 0 | 0 | 1 |
| ENSGACG00000020174 | ChrVII | 12750327 | 12755673 | capns1a | 1 | 0 | 0 | 0 | 1 | 0 |
| ENSGACG00000020401 | ChrVII | 17997085 | 18001531 | si:ch211-137i24.10 | 1 | 0 | 0 | 0 | 0 | 1 |
| ENSGACG00000020403 | ChrVII | 18007253 | 18010078 | slc25a35 | 1 | 0 | 0 | 0 | 0 | 1 |
| ENSGACG00000020493 | ChrVII | 19246409 | 19250932 | NIPS0P2 | 1 | 0 | 0 | 0 | 0 | 1 |
| ENSGACG00000020569 | ChrVII | 20326685 | 20369400 | frem2a | 1 | 0 | 0 | 0 | 0 | 1 |
| ENSGACG00000020634 | ChrVII | 21614701 | 21623583 | nectin1a | 1 | 0 | 0 | 0 | 1 | 0 |
| ENSGACG00000020724 | ChrVII | 23139706 | 23143482 | yipf5 | 1 | 0 | 0 | 0 | 1 | 0 |
| ENSGACG00000020725 | ChrVII | 23164545 | 23201619 | NR3C1 (1 of many) | 1 | 1 | 0 | 1 | 0 | 0 |
| ENSGACG00000020726 | ChrVII | 23208162 | 23253092 | ARHGAP26 | 1 | 0 | 0 | 0 | 1 | 0 |
| ENSGACG00000020734 | ChrVII | 23362870 | 23385166 | aff4 | 1 | 0 | 0 | 0 | 1 | 0 |

| | | | | | | | | | | |
|---|---|---|---|---|---|---|---|---|---|---|
| ENSGACG00000020754 | ChrVII | 24052097 | 24062908 | mybbp1a | 1 | 1 | 1 | 0 | 0 | 0 |
| ENSGACG00000020770 | ChrVII | 24240056 | 24244002 | htr3a | 1 | 0 | 0 | 0 | 1 | 0 |
| ENSGACG00000020803 | ChrVII | 25380828 | 25384895 | | 1 | 0 | 0 | 0 | 0 | 1 |
| ENSGACG00000020838 | ChrVII | 26287744 | 26291180 | | 1 | 0 | 0 | 0 | 1 | 0 |
| ENSGACG00000020848 | ChrVII | 26791004 | 26845216 | doc2b | 1 | 1 | 1 | 0 | 0 | 0 |
| ENSGACG00000020852 | ChrVII | 26876308 | 26878683 | | 1 | 0 | 0 | 0 | 1 | 0 |
| ENSGACG00000021530 | ChrIII | 13882346 | 13882414 | | 1 | 0 | 0 | 0 | 1 | 0 |
| ENSGACG00000022165 | ChrV | 11138845 | 11138931 | | 1 | 0 | 0 | 0 | 1 | 0 |

**Table S3A:** Tukey HSD post hoc test results for survival rate. Significant comparisons are highlighted in bold.

| Comparison | Estimate | Std. Error | z value | Pr(>\|z\|) |
|---|---|---|---|---|
| **WG_33 - control** | **-0.73765** | **0.26233** | **-2.812** | **0.03876** |
| **WG_6 - control** | -0.40963 | 0.27564 | -1.486 | 0.56738 |
| **TG_33 - control** | **-144.421** | **0.25161** | **-5.740** | **< 0.001** |
| **TG_06 - control** | -0.07637 | 0.26508 | -0.288 | 0.99847 |
| **WG_06 - WG_33** | 0.32802 | 0.24052 | 1.364 | 0.64713 |
| **TG_33 - WG_33** | **-0.70656** | **0.21261** | **-3.323** | **0.00769** |
| **TG_6 - WG_33** | **0.66128** | **0.22859** | **2.893** | **0.03051** |
| **TG_33 - WG_6** | **-103.457** | **0.23058** | **-4.487** | **< 0.001** |
| **TG_6 - WG_6** | 0.33327 | 0.24674 | 1.351 | 0.65554 |
| **TG_6 - TG_33** | **136.784** | **0.19398** | **7.052** | **< 0.001** |

**Table S3B:** Tukey HSD post hoc test results for standard length. Significant comparisons are highlighted in bold.

| Comparison | Estimate | Std. Error | z value | Pr(>\|z\|) |
|---|---|---|---|---|
| **WG_33 - control** | -0.002895 | 0.056225 | -0.051 | 10.000 |
| **WG_6 - control** | 0.074989 | 0.055932 | 1.341 | 0.6649 |
| **TG_33 - control** | 0.029907 | 0.059175 | 0.505 | 0.9868 |
| **TG_06 - control** | **0.172371** | **0.059205** | **2.911** | **0.0296** |
| **WG_06 - WG_33** | 0.077884 | 0.056181 | 1.386 | 0.6357 |
| **TG_33 - WG_33** | 0.032802 | 0.059405 | 0.552 | 0.9816 |
| **TG_6 - WG_33** | **0.175265** | **0.059467** | **2.947** | **0.0265** |
| **TG_33 - WG_6** | -0.045082 | 0.058994 | -0.764 | 0.9406 |
| **TG_6 - WG_6** | 0.097382 | 0.058991 | 1.651 | 0.4640 |
| **TG_6 - TG_33** | 0.142464 | 0.055749 | 2.555 | 0.0786 |

**Table S3C:** Tukey HSD post hoc test results for hepatosomatic index. Significant comparisons are highlighted in bold.

| Comparison | Estimate | Std. Error | z value | Pr(>\|z\|) |
|---|---|---|---|---|
| **WG_33 - control** | -0.13514 | 0.11511 | -1.174 | 0.76590 |
| **WG_6 - control** | 0.26415 | 0.11417 | 2.314 | 0.14025 |
| **TG_33 - control** | -0.05518 | 0.11950 | -0.462 | 0.99064 |
| **TG_06 - control** | **0.40834** | **0.11958** | **3.415** | **0.00574** |
| **WG_06 - WG_33** | **0.39929** | **0.11504** | **3.471** | **0.00464** |
| **TG_33 - WG_33** | 0.07996 | 0.12033 | 0.665 | 0.96384 |
| **TG_6 - WG_33** | **0.54348** | **0.12049** | **4.511** | **< 0.001** |
| **TG_33 - WG_6** | -0.31933 | 0.11909 | -2.681 | 0.05654 |
| **TG_6 - WG_6** | 0.14419 | 0.11907 | 1.211 | 0.74460 |
| **TG_6 - TG_33** | **0.46352** | **0.11373** | **4.076** | **< 0.001** |

**Table S3D:** Tukey HSD post hoc test results for total weight. Significant comparisons are highlighted in bold.

| Comparison | Estimate | Std. Error | z value | Pr(>\|z\|) |
|---|---|---|---|---|
| **WG_33 - control** | 0.004969 | 0.015050 | 0.330 | 0.9974 |
| **WG_6 - control** | 0.025076 | 0.014925 | 1.680 | 0.4456 |
| **TG_33 - control** | 0.001846 | 0.015738 | 0.117 | 10.000 |
| **TG_06 - control** | **0.045953** | **0.015752** | **2.917** | **0.0290** |
| **WG_06 - WG_33** | 0.020106 | 0.015039 | 1.337 | 0.6674 |
| **TG_33 - WG_33** | -0.003123 | 0.015840 | -0.197 | 0.9997 |
| **TG_6 - WG_33** | 0.040984 | 0.015871 | 2.582 | 0.0734 |
| **TG_33 - WG_6** | -0.023230 | 0.015661 | -1.483 | 0.5727 |
| **TG_6 - WG_6** | 0.020877 | 0.015659 | 1.333 | 0.6697 |
| **TG_6 - TG_33** | **0.044107** | **0.014845** | **2.971** | **0.0247** |

**Table S4:** Summary statistics for whole genome re-sequencing of wild caught sticklebacks. Six stickleback populations (KIE = Kiel, NYN = Nynäshamans, SYL= Sylt, FAL = Falsterbo, LET = Letipea, BAR = Barsta) were sampled at the corresponding latitude and longitude and salinity. Fish ID is given, along with standard length and sex.

| ID | sex | site name | latitude | longitude | salinity | standard length (mm) | total reads | reads aligend | aligned reads in pairs | high quality reads | mean coverage | sd coverage | mean insert size |
|---|---|---|---|---|---|---|---|---|---|---|---|---|---|
| S1 | female | FAL | 55.412955 | 12.931189 | 9.8 | 31 | 53099898 | 51454273 | 51059422 | 46005389 | 13.77 | 8.14 | 389.83 |
| S10 | male | SYL | 55.016166 | 8.439550 | 28.9 | 24 | 59249088 | 57721239 | 57309340 | 51392222 | 15.29 | 8.75 | 378.09 |
| S11 | male | LET | 59.5521 | 26.60826 | 4.3 | 24 | 64732112 | 62992502 | 62514646 | 55977942 | 16.47 | 9.16 | 379.76 |
| S12 | male | LET | 59.5521 | 26.60826 | 4.3 | 28 | 60505260 | 58806659 | 58363258 | 52747595 | 15.54 | 8.74 | 368.95 |
| S13 | female | LET | 59.5521 | 26.60826 | 4.3 | 40 | 46451078 | 45136076 | 44795792 | 40774872 | 12.08 | 7.18 | 371.11 |
| S14 | female | LET | 59.5521 | 26.60826 | 4.3 | 28 | 52566384 | 51070498 | 50676508 | 46196326 | 13.6 | 7.77 | 361.36 |
| S15 | female | FAL | 55.412955 | 12.931189 | 9.8 | 32 | 53175370 | 51740418 | 51356498 | 45760642 | 13.58 | 8.07 | 367.76 |
| S16 | female | FAL | 55.412955 | 12.931189 | 9.8 | 31 | 54746754 | 53085246 | 52694874 | 47758257 | 14.11 | 8.07 | 368.2 |
| S17 | female | SYL | 55.016166 | 8.439550 | 28.9 | 25 | 42687172 | 41590244 | 41276164 | 37539956 | 11.26 | 6.83 | 383.5 |
| S18 | female | SYL | 55.016166 | 8.439550 | 28.9 | 23 | 55674944 | 54215662 | 53789138 | 48770957 | 14.52 | 8.22 | 374 |
| S19 | female | KIE | 54.436621 | 10.17228 | 18 | 33 | 51339478 | 49896745 | 49534292 | 44841451 | 13.45 | 7.83 | 378.17 |
| S2 | female | FAL | 55.412955 | 12.931189 | 9.8 | 31 | 67830816 | 65879708 | 65386846 | 59023635 | 17.5 | 9.33 | 383.05 |
| S20 | female | KIE | 54.436621 | 10.17228 | 18 | 35 | 45388352 | 44131200 | 43780040 | 39724153 | 11.81 | 7.19 | 363.19 |
| S21 | male | NYN | 58.8790698 | 17.935061 | 6 | 25 | 62740566 | 60992914 | 60552808 | 55067013 | 16.42 | 8.8 | 378.55 |
| S22 | male | NYN | 58.8790698 | 17.935061 | 6 | 27 | 58127744 | 56500828 | 56070618 | 51359209 | 15.33 | 8.33 | 380.72 |
| S23 | male | BAR | 62.8630833 | 18.3975 | 4.6 | 17 | 66400620 | 64557384 | 64076342 | 57948807 | 17.25 | 9.29 | 381.66 |
| S24 | male | BAR | 62.8630833 | 18.3975 | 4.6 | 19 | 52888922 | 51354675 | 50965362 | 45750005 | 13.66 | 8.04 | 377.71 |
| S25 | female | BAR | 62.8630833 | 18.3975 | 4.6 | 19 | 55355880 | 53867446 | 53461396 | 48579009 | 14.63 | 8.13 | 383.57 |
| S26 | female | BAR | 62.8630833 | 18.3975 | 4.6 | 18 | 60545964 | 58904170 | 58457402 | 52851197 | 15.76 | 8.69 | 382.32 |
| S27 | female | LET | 59.5521 | 26.60826 | 4.3 | 27 | 48256944 | 46901465 | 46558408 | 42198012 | 12.62 | 7.42 | 376.13 |
| S28 | female | LET | 59.5521 | 26.60826 | 4.3 | 27 | 48321396 | 46997345 | 46655746 | 42196863 | 12.59 | 7.47 | 372.53 |

| | | | | | | | | | | | | | |
|---|---|---|---|---|---|---|---|---|---|---|---|---|---|
| S29 | female | FAL | 55.412955 | 12.931189 | 9.8 | 31 | 53423426 | 51720960 | 51334050 | 46751683 | 14.07 | 7.85 | 382.41 |
| S3 | female | KIE | 54.436621 | 10.17228 | 18 | 33 | 51511534 | 50109217 | 49751260 | 44785482 | 13.23 | 7.75 | 371.95 |
| S30 | female | FAL | 55.412955 | 12.931189 | 9.8 | 31 | 62572564 | 60802237 | 60355732 | 54743424 | 16.31 | 8.85 | 382.79 |
| S31 | female | SYL | 55.016166 | 8.439550 | 28.9 | 24 | 52336108 | 51000812 | 50630182 | 45718213 | 13.74 | 7.89 | 383.56 |
| S32 | female | SYL | 55.016166 | 8.439550 | 28.9 | 25 | 50659258 | 49422266 | 49071064 | 44058329 | 13.21 | 7.87 | 376.03 |
| S33 | female | KIE | 54.436621 | 10.17228 | 18 | 29 | 59992552 | 58304582 | 57854052 | 52465334 | 15.72 | 8.69 | 390.69 |
| S34 | female | KIE | 54.436621 | 10.17228 | 18 | 29 | 52958328 | 51617586 | 51231202 | 46633547 | 14.08 | 7.96 | 395.11 |
| S35 | male | NYN | 58.8790698 | 17.935061 | 6 | 39 | 54066258 | 52535615 | 52111994 | 46944359 | 13.96 | 8.12 | 385.2 |
| S36 | male | NYN | 58.8790698 | 17.935061 | 6 | 26 | 58979382 | 57446489 | 57021398 | 52137658 | 15.58 | 8.42 | 377.63 |
| S37 | female | NYN | 58.8790698 | 17.935061 | 6 | 53 | 60917462 | 59237862 | 58787416 | 53314193 | 15.88 | 8.72 | 386.29 |
| S38 | female | NYN | 58.8790698 | 17.935061 | 6 | 32 | 66539062 | 64563355 | 64037068 | 57547546 | 17.02 | 9.34 | 379.2 |
| S39 | female | BAR | 62.8630833 | 18.3975 | 4.6 | 18 | 53386772 | 51907116 | 51493034 | 46685925 | 13.92 | 7.93 | 383.5 |
| S4 | female | KIE | 54.436621 | 10.17228 | 18 | 34 | 53563882 | 52156214 | 51776690 | 46822429 | 13.9 | 8.02 | 373.12 |
| S40 | female | BAR | 62.8630833 | 18.3975 | 4.6 | 17 | 64856498 | 62844156 | 62366656 | 56901784 | 16.85 | 8.92 | 384.81 |
| S41 | female | LET | 59.5521 | 26.60826 | 4.3 | 33 | 46380788 | 45084548 | 44741886 | 40738791 | 12.03 | 7.27 | 369.47 |
| S42 | female | LET | 59.5521 | 26.60826 | 4.3 | 36 | 46539888 | 44942891 | 44542668 | 40288199 | 11.93 | 7.56 | 371.35 |
| S43 | female | FAL | 55.412955 | 12.931189 | 9.8 | 29 | 44422988 | 43191463 | 42867562 | 38818162 | 11.62 | 7.04 | 387 |
| S44 | female | FAL | 55.412955 | 12.931189 | 9.8 | 28 | 44980018 | 43519423 | 43174784 | 39637313 | 11.95 | 7.17 | 386.08 |
| S45 | male | SYL | 55.016166 | 8.439550 | 28.9 | 39 | 60809022 | 59192864 | 58753924 | 53143246 | 15.73 | 8.56 | 373.84 |
| S46 | male | SYL | 55.016166 | 8.439550 | 28.9 | 26 | 57211502 | 55688535 | 55274846 | 49799828 | 14.74 | 8.4 | 374.03 |
| S47 | male | KIE | 54.436621 | 10.17228 | 18 | 32 | 52163506 | 50721768 | 50346708 | 45023344 | 13.33 | 8.14 | 380.26 |
| S48 | male | KIE | 54.436621 | 10.17228 | 18 | 33 | 59561780 | 57906928 | 57451994 | 52022840 | 15.51 | 8.69 | 381.48 |
| S49 | male | KIE | 54.436621 | 10.17228 | 18 | 31 | 50842586 | 49392770 | 49017606 | 44150548 | 13.15 | 7.88 | 380.16 |
| S5 | male | NYN | 58.8790698 | 17.935061 | 6 | 24 | 48136868 | 46740050 | 46372224 | 42197802 | 12.59 | 7.32 | 376.67 |
| S50 | male | KIE | 54.436621 | 10.17228 | 18 | 32 | 70840292 | 68994230 | 68484716 | 61556269 | 18.36 | 9.95 | 393.8 |
| S51 | female | NYN | 58.8790698 | 17.935061 | 6 | 28 | 57912144 | 55709732 | 55285338 | 49719403 | 14.83 | 8.35 | 387.28 |
| S52 | female | NYN | 58.8790698 | 17.935061 | 6 | 43 | 45198976 | 43755361 | 43409824 | 39073539 | 11.53 | 7.18 | 376.31 |
| S53 | female | BAR | 62.8630833 | 18.3975 | 4.6 | 18 | 53204092 | 51613898 | 51197438 | 46606570 | 13.98 | 7.87 | 397.72 |
| S54 | female | BAR | 62.8630833 | 18.3975 | 4.6 | 17 | 67193056 | 65208735 | 64691376 | 58220165 | 17.28 | 9.31 | 399 |

| | | | | | | | | | | | | | |
|---|---|---|---|---|---|---|---|---|---|---|---|---|---|
| S55 | female | LET | 59.5521 | 26.60826 | 4.3 | 27 | 45627856 | 44381119 | 44043566 | 40020911 | 11.82 | 7.15 | 371.47 |
| S56 | female | LET | 59.5521 | 26.60826 | 4.3 | 27 | 49241638 | 47879579 | 47518742 | 42867415 | 12.74 | 7.61 | 382.12 |
| S57 | male | FAL | 55.412955 | 12.931189 | 9.8 | 32 | 53853956 | 52186204 | 51764358 | 46379606 | 13.81 | 8.25 | 396.87 |
| S58 | male | FAL | 55.412955 | 12.931189 | 9.8 | 31 | 48604482 | 47119809 | 46753202 | 42028951 | 12.58 | 7.77 | 394.99 |
| S59 | male | SYL | 55.016166 | 8.439550 | 28.9 | 23 | 52963100 | 51536646 | 51142932 | 45914061 | 13.66 | 8.05 | 394.22 |
| S6 | male | NYN | 58.8790698 | 17.935061 | 6 | 23 | 53610228 | 52153201 | 51764414 | 47154571 | 13.92 | 7.82 | 373.57 |
| S60 | male | SYL | 55.016166 | 8.439550 | 28.9 | 23 | 44376314 | 43229355 | 42902698 | 38295557 | 11.43 | 7.27 | 393.46 |
| S61 | male | SYL | 55.016166 | 8.439550 | 28.9 | 23 | 50150228 | 48842901 | 48471972 | 43611845 | 13.01 | 7.81 | 395.23 |
| S62 | male | SYL | 55.016166 | 8.439550 | 28.9 | 24 | 53219332 | 51778288 | 51372236 | 46356208 | 13.87 | 8.07 | 397.38 |
| S63 | male | KIE | 54.436621 | 10.17228 | 18 | 31 | 45157340 | 43911400 | 43576258 | 39284752 | 11.79 | 7.4 | 393.17 |
| S64 | male | KIE | 54.436621 | 10.17228 | 18 | 30 | 64161760 | 62357886 | 61833264 | 55332302 | 16.39 | 9.25 | 393.29 |
| S65 | female | NYN | 58.8790698 | 17.935061 | 6 | 24 | 57552266 | 55939246 | 55491846 | 50139709 | 15.04 | 8.57 | 393.1 |
| S66 | female | NYN | 58.8790698 | 17.935061 | 6 | 21 | 41545924 | 40452609 | 40146890 | 36478915 | 11 | 6.79 | 393.16 |
| S67 | female | BAR | 62.8630833 | 18.3975 | 4.6 | 18 | 40847930 | 39735959 | 39425668 | 35714454 | 10.71 | 6.78 | 385 |
| S68 | female | BAR | 62.8630833 | 18.3975 | 4.6 | 17 | 52104600 | 50722288 | 50345550 | 45557562 | 13.67 | 8.04 | 385.78 |
| S69 | male | LET | 59.5521 | 26.60826 | 4.3 | 39 | 54058430 | 52458887 | 52051058 | 46460762 | 13.62 | 8.3 | 368.57 |
| S7 | male | BAR | 62.8630833 | 18.3975 | 4.6 | 20 | 41735988 | 40611287 | 40308232 | 36537433 | 10.92 | 6.84 | 376.44 |
| S70 | male | LET | 59.5521 | 26.60826 | 4.3 | 32 | 49486134 | 48035380 | 47659980 | 42943215 | 12.68 | 7.72 | 378.01 |
| S71 | male | FAL | 55.412955 | 12.931189 | 9.8 | 33 | 53822154 | 52317878 | 51917988 | 46837175 | 13.99 | 8.22 | 388.15 |
| S72 | male | FAL | 55.412955 | 12.931189 | 9.8 | 30 | 59383166 | 57356093 | 56921304 | 50808323 | 15.14 | 8.79 | 383.64 |
| S73 | female | SYL | 55.016166 | 8.439550 | 28.9 | 26 | 41531640 | 40400645 | 40088336 | 36348810 | 10.87 | 6.81 | 388.05 |
| S74 | female | SYL | 55.016166 | 8.439550 | 28.9 | 24 | 48253694 | 46856163 | 46505036 | 42070531 | 12.48 | 7.41 | 386.77 |
| S75 | male | FAL | 55.412955 | 12.931189 | 9.8 | 28 | 48901972 | 47102565 | 46734180 | 42471863 | 12.77 | 7.62 | 398.07 |
| S76 | male | FAL | 55.412955 | 12.931189 | 9.8 | 30 | 54727256 | 52172085 | 51768210 | 46714855 | 13.99 | 8.21 | 394.86 |
| S77 | male | KIE | 54.436621 | 10.17228 | 18 | 27 | 54971942 | 53397990 | 52993618 | 47682831 | 14,25 | 8.39 | 391.34 |
| S78 | male | KIE | 54.436621 | 10.17228 | 18 | 29 | 74182672 | 72307267 | 71771586 | 65049831 | 19.4 | 10.26 | 391.66 |
| S79 | female | NYN | 58.8790698 | 17.935061 | 6 | 27 | 51125432 | 49844267 | 49475822 | 44547647 | 13.29 | 7.89 | 384.18 |
| S8 | male | BAR | 62.8630833 | 18.3975 | 4.6 | 18 | 41776818 | 40605236 | 40286314 | 36225955 | 10.81 | 6.98 | 378.65 |
| S80 | female | NYN | 58.8790698 | 17.935061 | 6 | 31 | 53096178 | 51557239 | 51160266 | 46567931 | 13.96 | 8 | 392.39 |

| | | | | | | | | | | | | | |
|---|---|---|---|---|---|---|---|---|---|---|---|---|---|
| S81 | male | BAR | 62.8630833 | 18.3975 | 4.6 | 20 | 46138468 | 44815095 | 44443792 | 40083115 | 11.95 | 7.4 | 394.98 |
| S82 | male | BAR | 62.8630833 | 18.3975 | 4.6 | 18 | 46487280 | 45206943 | 44852368 | 40245067 | 12.01 | 7.49 | 391.06 |
| S83 | male | LET | 59.5521 | 26.60826 | 4.3 | 39 | 56123940 | 54610807 | 54200548 | 48467659 | 14.28 | 8.38 | 382.43 |
| S84 | male | LET | 59.5521 | 26.60826 | 4.3 | 24 | 55394104 | 53829494 | 53424150 | 47684275 | 14.08 | 8.33 | 383.97 |
| S85 | male | NYN | 58.8790698 | 17.935061 | 6 | 28 | 57048792 | 55082164 | 54657306 | 48888229 | 14.51 | 8.43 | 393.84 |
| S86 | male | NYN | 58.8790698 | 17.935061 | 6 | 22 | 58154884 | 56657681 | 56204760 | 51137618 | 15.29 | 8.43 | 401.78 |
| S87 | female | SYL | 55.016166 | 8.439550 | 28.9 | 26 | 62919890 | 61286537 | 60817766 | 54723790 | 16.32 | 9.02 | 389.66 |
| S88 | female | SYL | 55.016166 | 8.439550 | 28.9 | 23 | 65075134 | 63336434 | 62821978 | 56786117 | 16.7 | 8.92 | 381.06 |
| S89 | female | KIE | 54.436621 | 10.17228 | 18 | 33 | 48427088 | 47150565 | 46794656 | 41828226 | 12.37 | 7.56 | 382.58 |
| S9 | male | SYL | 55.016166 | 8.439550 | 28.9 | 23 | 54509806 | 52932957 | 52517236 | 47354023 | 14.03 | 8.06 | 378.54 |
| S90 | female | KIE | 54.436621 | 10.17228 | 18 | 31 | 37045098 | 36076704 | 35816298 | 32277497 | 9.59 | 6.29 | 379.6 |
| S91 | male | FAL | 55.412955 | 12.931189 | 9.8 | 30 | 36778390 | 35393559 | 35095258 | 31680643 | 9.49 | 6.22 | 389.75 |
| S92 | male | FAL | 55.412955 | 12.931189 | 9.8 | 27 | 38582222 | 34925224 | 34645566 | 30986369 | 9.27 | 6.39 | 388.5 |
| S93 | male | LET | 59.5521 | 26.60826 | 4.3 | 24 | 60554766 | 58849331 | 58393636 | 52673246 | 15.51 | 8.83 | 380.96 |
| S94 | male | LET | 59.5521 | 26.60826 | 4.3 | 33 | 51305380 | 49375369 | 48965854 | 43399599 | 12.4 | 8.55 | 349.06 |
| S95 | male | BAR | 62.8630833 | 18.3975 | 4.6 | 17 | 70846994 | 68870460 | 68353680 | 61093010 | 18.09 | 9.81 | 389.65 |
| S96 | male | BAR | 62.8630833 | 18.3975 | 4.6 | 18 | 44379656 | 43138160 | 42810028 | 38388253 | 11.46 | 7.31 | 386.37 |

**Table S5A**: Summary statistics for the RRBS of experimental fish. Samples of the two-generation salinity acclimation experiment with sticklebacks from the Kiel population (20 PSU) are listed with their respective sex (f = female, m = male), parental and offspring salinity treatment condition and standard length. To align quality filtered reads we used *Bismark v0.17.0*[1] with *Bowtie2 v2.3.3*. The conversion rates were calculated using the *cegxQC* software[2] on the unfiltered data (see Methods for details).

| ID | sex | parental salinity (PSU) | offspring salinity (PSU) | standard length (mm) | total reads | unique aligned | unaligned | % mapping efficiency | % methylated Cs in CpGs | % methylated Cs in CHGs | % methylated Cs in CHHs | conversion rate of Cs | conversion rate of mCs |
|---|---|---|---|---|---|---|---|---|---|---|---|---|---|
| RS12 | f | 6 | 6 | 34 | 19,193,500 | 13,653,401 | 3,387,767 | 71.1 | 47.6 | 0.4 | 0.4 | 99.7 | 2.4 |
| RS28 | f | 6 | 6 | 32 | 21,245,180 | 14,488,176 | 4,014,433 | 68.2 | 55.4 | 0.5 | 0.4 | 99.6 | 2.4 |
| RS34 | m | 6 | 6 | 34 | 21,776,314 | 14,731,355 | 3,937,799 | 67.6 | 55.8 | 0.5 | 0.4 | 99.7 | 2.3 |
| RS39 | m | 6 | 6 | 35 | 15,295,670 | 10,362,419 | 2,886,758 | 67.7 | 53 | 0.4 | 0.4 | 99.6 | 2.4 |
| RS47 | m | 6 | 6 | 33 | 21,588,032 | 15,209,910 | 4,024,708 | 70.5 | 54.1 | 0.5 | 0.4 | 99.6 | 2.3 |
| RS55 | f | 6 | 6 | 32 | 14,812,705 | 10,392,489 | 2,728,068 | 70.2 | 57.9 | 0.5 | 0.4 | 99.6 | 2.3 |
| RS6 | f | 6 | 6 | 39 | 20,614,412 | 12,789,884 | 3,554,303 | 62 | 56.1 | 0.5 | 0.4 | 99.6 | 2.5 |
| RS68 | f | 6 | 6 | 35 | 15,441,713 | 11,117,645 | 2,848,520 | 72 | 56.7 | 0.5 | 0.4 | 99.6 | 2.5 |
| RS69 | m | 6 | 6 | 34 | 22,005,986 | 15,131,296 | 4,024,938 | 68.8 | 57.6 | 0.5 | 0.4 | 99.6 | 2.3 |
| RS73 | f | 6 | 6 | 35 | 19,651,081 | 13,388,904 | 3,536,242 | 68.1 | 54.8 | 0.5 | 0.4 | 99.6 | 2.5 |
| RS78 | m | 6 | 6 | 36 | 19,552,515 | 13,575,577 | 3,811,237 | 69.4 | 52.7 | 0.5 | 0.4 | 99.5 | 2.3 |
| RS9 | m | 6 | 6 | 36 | 18,521,081 | 11,376,729 | 3,324,153 | 61.4 | 48.6 | 0.4 | 0.3 | 99.6 | 2.5 |
| RS1 | f | 20 | 6 | 33 | 19,410,063 | 12,672,156 | 3,393,105 | 65.3 | 59.8 | 0.5 | 0.4 | 99.6 | 2.4 |
| RS18 | m | 20 | 6 | 33 | 22,139,187 | 14,733,925 | 4,023,405 | 66.6 | 58.3 | 0,5 | 0.4 | 99.5 | 2.5 |
| RS19 | m | 20 | 6 | 34 | 25,669,395 | 17,692,358 | 4,727,760 | 68.9 | 57.8 | 0.5 | 0.4 | 99.5 | 2.3 |
| RS22 | m | 20 | 6 | 34 | 21,431,022 | 14,848,843 | 3,781,126 | 69.3 | 60.6 | 0.5 | 0.4 | 99.7 | 2.5 |
| RS24 | f | 20 | 6 | 34 | 18,563,526 | 12,700,457 | 3,099,110 | 68.4 | 64.9 | 0.5 | 0.4 | 99.5 | 2.4 |
| RS33 | f | 20 | 6 | 31 | 19,219,593 | 12,738,183 | 3,475,010 | 66.3 | 57 | 0.5 | 0.4 | 99.7 | 2.4 |
| RS5 | f | 20 | 6 | 39 | 18,474,615 | 12,876,991 | 3,113,033 | 69.7 | 50.8 | 0.5 | 0.4 | 99.6 | 2.4 |
| RS52 | m | 20 | 6 | 34 | 19,884,602 | 14,145,657 | 3,763,333 | 71.1 | 53.7 | 0.5 | 0.4 | 99.7 | 2.6 |
| RS61 | f | 20 | 6 | 35 | 20,691,948 | 14,272,889 | 3,761,703 | 69 | 55.4 | 0.5 | 0.4 | 99.6 | 2.2 |
| RS77 | m | 20 | 6 | 34 | 17,922,261 | 12,485,632 | 3,351,014 | 69.7 | 50.8 | 0.5 | 0.4 | 99.6 | 2.5 |
| RS79 | m | 20 | 6 | 33 | 11,936,993 | 7,973,063 | 2,435,528 | 66.8 | 52.1 | 0.5 | 0.4 | 99.6 | 2.4 |
| RS11 | f | 20 | 20 | 31 | 20,243,439 | 14,268,055 | 3,663,464 | 70.5 | 46.8 | 0.4 | 0.4 | 99.6 | 2.4 |

| | | | | | | | | | | | | | |
|---|---|---|---|---|---|---|---|---|---|---|---|---|---|
| RS14 | m | 20 | 20 | 33 | 18,299,908 | 12,488,117 | 3,243,269 | 68.2 | 57.8 | 0.5 | 0.4 | 99.6 | 2.1 |
| RS2 | f | 20 | 20 | 31 | 20,137,550 | 13,212,235 | 3,509,171 | 65.6 | 61.3 | 0.5 | 0.4 | 99.5 | 2.4 |
| RS27 | m | 20 | 20 | 34 | 17,234,518 | 11,807,735 | 3,147,777 | 68.5 | 47.9 | 0.4 | 0.4 | 99.6 | 2.4 |
| RS30 | f | 20 | 20 | 33 | 20,652,539 | 14,071,954 | 3,717,746 | 68.1 | 56.5 | 0.5 | 0.4 | 99.6 | 2.4 |
| RS36 | f | 20 | 20 | 33 | 18,842,860 | 13,035,496 | 3,439,400 | 69.2 | 59.2 | 0.5 | 0.4 | 99.6 | 2.5 |
| RS46 | f | 20 | 20 | 31 | 14,955,973 | 10,166,362 | 2,653,749 | 68 | 51.4 | 0.5 | 0.4 | 99.4 | 2.4 |
| RS48 | m | 20 | 20 | 36 | 23,172,203 | 16,234,065 | 4,254,154 | 70.1 | 53.6 | 0.4 | 0.4 | 99.6 | 2.5 |
| RS59 | f | 20 | 20 | 29 | 19,697,111 | 14,065,491 | 3,437,252 | 71.4 | 57.5 | 0.5 | 0.4 | 99.7 | 2.4 |
| RS62 | m | 20 | 20 | 34 | 21,508,796 | 14,858,533 | 4,028,818 | 69.1 | 52.6 | 0.5 | 0.4 | 99.5 | 2.4 |
| RS7 | m | 20 | 20 | 34 | 15,890,812 | 10,638,953 | 3,121,759 | 67 | 55.1 | 0.5 | 0.4 | 99.6 | 2.4 |
| RS75 | m | 20 | 20 | 33 | 22,640,803 | 14,627,178 | 4,154,518 | 64.6 | 50.6 | 0.5 | 0.4 | 99.6 | 2.3 |
| RS16 | m | 20 | 33 | 36 | 23,379,805 | 16,349,654 | 4,392,411 | 69.9 | 50.3 | 0.4 | 0.3 | 99.7 | 2.5 |
| RS17 | f | 20 | 33 | 33 | 15,365,773 | 10,794,519 | 2,861,846 | 70.3 | 53.4 | 0.4 | 0.3 | 99.6 | 2.5 |
| RS23 | m | 20 | 33 | 38 | 23,257,469 | 15,971,106 | 4,134,190 | 68.7 | 63.9 | 0.5 | 0.4 | 99.4 | 2.3 |
| RS31 | f | 20 | 33 | 34 | 18,197,686 | 12,270,798 | 3,370,685 | 67.4 | 55.4 | 0.5 | 0.4 | 99.6 | 2.4 |
| RS45 | m | 20 | 33 | 34 | 19,293,724 | 13,575,990 | 3,709,491 | 70.4 | 46.6 | 0.4 | 0.4 | 99.6 | 2.4 |
| RS49 | f | 20 | 33 | 35 | 22,586,319 | 16,247,681 | 3,991,975 | 71.9 | 51.5 | 0.4 | 0.4 | 99.6 | 2.5 |
| RS56 | m | 20 | 33 | 35 | 12,837,376 | 8,486,651 | 2,417,279 | 66.1 | 56.7 | 0.5 | 0.4 | 99.7 | 2.7 |
| RS65 | f | 20 | 33 | 37 | 14,645,372 | 10,073,076 | 2,768,136 | 68.8 | 56 | 0.4 | 0.3 | 99.6 | 2.5 |
| RS66 | m | 20 | 33 | 32 | 20,732,423 | 14,118,267 | 3,957,707 | 68.1 | 60 | 0.5 | 0.4 | 99.6 | 2.4 |
| RS72 | f | 20 | 33 | 30 | 23,099,522 | 15,824,314 | 4,015,465 | 68.5 | 55.8 | 0.5 | 0.4 | 99.6 | 2.4 |
| RS8 | m | 20 | 33 | 30 | 14,834,881 | 9,428,882 | 2,726,520 | 63.6 | 59.9 | 0.5 | 0.4 | 99.5 | 2.4 |
| RS10 | f | 33 | 33 | 29 | 17,199,844 | 11,861,027 | 3,060,336 | 69 | 49.1 | 0.4 | 0.4 | 99.5 | 2.6 |
| RS13 | m | 33 | 33 | 32 | 19,322,020 | 12,524,534 | 3,508,974 | 64.8 | 49 | 0.4 | 0.4 | 99.6 | 2.5 |
| RS26 | m | 33 | 33 | 35 | 18,323,492 | 12,085,071 | 3,485,139 | 66 | 53.2 | 0.5 | 0.4 | 99.7 | 2.4 |
| RS29 | f | 33 | 33 | 32 | 20,178,134 | 13,061,973 | 3,634,233 | 64.7 | 50.1 | 0.5 | 0.4 | 99.6 | 2.4 |
| RS38 | f | 33 | 33 | 33 | 19,425,374 | 12,864,087 | 3,481,184 | 66.2 | 55.1 | 0.5 | 0.4 | 99.6 | 2.4 |
| RS42 | m | 33 | 33 | 35 | 16,966,704 | 11,139,312 | 3,146,420 | 65.7 | 54.9 | 0.5 | 0.4 | 99.7 | 2.6 |
| RS44 | m | 33 | 33 | 37 | 15,454,040 | 10,661,431 | 2,861,799 | 69 | 47.3 | 0.4 | 0.3 | 99.6 | 2.5 |
| RS50 | f | 33 | 33 | 32 | 22,655,597 | 15,608,421 | 3,964,156 | 68.9 | 60.2 | 0.5 | 0.4 | 99.7 | 2.4 |
| RS58 | f | 33 | 33 | 37 | 19,602,242 | 13,608,515 | 3,279,118 | 69.4 | 60.2 | 0.5 | 0.4 | 99.6 | 2.3 |
| RS60 | m | 33 | 33 | 36 | 19,446,537 | 13,374,949 | 3,584,079 | 68.8 | 56.8 | 0.5 | 0.4 | 99.6 | 2.4 |
| RS63 | f | 33 | 33 | 35 | 18,322,127 | 12,880,934 | 3,404,665 | 70.3 | 52.9 | 0.5 | 0.4 | 99.6 | 2.5 |
| RS71 | m | 33 | 33 | 37 | 16,379,683 | 11,242,454 | 3,138,323 | 68.6 | 58.1 | 0.5 | 0.4 | 99.6 | 2.4 |

**Table S5B**: Summary statistics for the RRBS of wild caught fish. Samples of three populations (KIE = Kiel, NYN = Nynäshamn, SYL = Sylt) collected in- and outside of the Baltic Sea salinity gradient are listed with their respective sex (f = female, m = male) and salinity of origin. To align quality filtered reads we used *Bismark v0.17.0*[1] with *Bowtie2 v2.3.3*. The conversion rates were calculated using the *cegxQC* software[2] on the unfiltered data (see Methods for details).

| ID | sex | location | salinity | total reads | unique aligned | unaligned | % mapping effiency | % methylated Cs in CpG | % methylated Cs in CHG | % methylated Cs in CHH | Conversion rate of Cs | Conversion rate of mCs |
|---|---|---|---|---|---|---|---|---|---|---|---|---|
| S3 | f | KIE | 18 | 8,753,702 | 4,874,582 | 2,438,154 | 55.7 | 57.3 | 0.7 | 0.6 | 99.1 | 2.56 |
| S4 | f | KIE | 18 | 8,571,146 | 5,056,022 | 2,321,568 | 59.0 | 63.4 | 0.8 | 0.7 | 99.2 | 2.49 |
| S5 | m | NYN | 6 | 7,232,265 | 4,454,780 | 1,997,286 | 61.6 | 62.6 | 0.7 | 0.7 | 99.3 | 2.50 |
| S6 | m | NYN | 6 | 8,181,388 | 4,890,678 | 2,307,322 | 59.8 | 59.1 | 0.7 | 0.7 | 99.2 | 2.57 |
| S9 | m | SYL | 28.9 | 7,791,070 | 4,420,213 | 2,295,644 | 56.7 | 58.1 | 0.7 | 0.6 | 99.2 | 2.50 |
| S10 | m | SYL | 28.9 | 7,753,903 | 4,145,109 | 2,254,765 | 53.5 | 57.6 | 0.7 | 0.7 | 99.3 | 2.57 |
| S17 | f | SYL | 28.9 | 11,091,390 | 7,159,358 | 2,352,179 | 64.5 | 56.4 | 0.6 | 0.5 | 99.5 | 2.48 |
| S18 | f | SYL | 28.9 | 11,411,538 | 7,481,972 | 2,501,782 | 65.6 | 59.3 | 0.6 | 0.5 | 99.4 | 2.39 |
| S19 | f | KIE | 18 | 5,744,795 | 3,662,505 | 1,369,060 | 63.8 | 58.4 | 0.6 | 0.5 | 99.4 | 2.49 |
| S20 | f | KIE | 18 | 10,006,121 | 6,408,060 | 2,200,372 | 64.0 | 58.3 | 0.6 | 0.5 | 99.4 | 2.43 |
| S21 | m | NYN | 6 | 10,724,717 | 6,639,862 | 2,498,410 | 61.9 | 58.5 | 0.6 | 0.5 | 99.4 | 2.39 |
| S22 | m | NYN | 6 | 9,966,267 | 6,734,287 | 2,092,066 | 67.6 | 64.5 | 0.6 | 0.5 | 99.3 | 2.30 |
| S32 | f | SYL | 28.9 | 8,570,141 | 4,986,526 | 2,135,955 | 58.2 | 63.1 | 0.8 | 0.7 | 99.2 | 2.47 |
| S33 | f | KIE | 18 | 9,332,513 | 5,401,129 | 2,435,485 | 57.9 | 57.8 | 0.7 | 0.6 | 99.3 | 2.52 |
| S34 | f | KIE | 18 | 7,902,135 | 4,901,313 | 2,126,350 | 62.0 | 57.3 | 0.7 | 0.6 | 99.3 | 2.44 |
| S35 | m | NYN | 6 | 9,525,328 | 5,327,975 | 2,421,326 | 55.9 | 58.9 | 0.7 | 0.6 | 99.3 | 2.39 |
| S36 | m | NYN | 6 | 14,623,815 | 9,020,345 | 3,974,462 | 61.7 | 57.2 | 0.7 | 0.6 | 99.4 | 2.43 |

| | | | | | | | | | | | | |
|---|---|---|---|---|---|---|---|---|---|---|---|---|
| S37 | f | NYN | 6 | 11,349,272 | 6,980,832 | 2,928,035 | 61.5 | 57.8 | 0.6 | 0.6 | 99.4 | 2.40 |
| S38 | f | NYN | 6 | 8,248,343 | 4,698,496 | 2,183,045 | 57.0 | 60.4 | 0.7 | 0.6 | 99.4 | 2.41 |
| S45 | m | SYL | 28.9 | 11,310,329 | 7,263,715 | 2,830,149 | 64.2 | 55.6 | 0.6 | 0.5 | 99.4 | 2.48 |
| S46 | m | SYL | 28.9 | 8,604,268 | 5,288,360 | 2,123,524 | 61.5 | 57.5 | 0.6 | 0.5 | 99.4 | 2.41 |
| S47 | m | KIE | 18 | 9,263,216 | 5,258,462 | 2,438,100 | 56.8 | 60.6 | 0.6 | 0.6 | 99.4 | 2.41 |
| S48 | m | KIE | 18 | 11,163,270 | 7,214,590 | 2,669,147 | 64.6 | 56.1 | 0.6 | 0.5 | 99.5 | 2.43 |
| S49 | m | KIE | 18 | 8,255,526 | 4,765,556 | 2,502,234 | 57.7 | 60.5 | 0.8 | 0.7 | 99.2 | 2.45 |
| S50 | m | KIE | 18 | 7,028,398 | 3,784,613 | 2,259,871 | 53.8 | 62.5 | 0.8 | 0.7 | 99.3 | 2.56 |
| S51 | f | NYN | 6 | 10,333,052 | 5,838,362 | 2,893,012 | 56.5 | 57.4 | 0.7 | 0.6 | 99.4 | 2.48 |
| S52 | f | NYN | 6 | 8,266,988 | 4,418,322 | 2,705,975 | 53.4 | 60.9 | 0.8 | 0.7 | 99.3 | 2.51 |
| S59 | m | SYL | 28.9 | 8,506,803 | 4,689,790 | 2,438,004 | 55.1 | 58.1 | 0.8 | 0.7 | 99.3 | 2.45 |
| S60 | m | SYL | 28.9 | 7,561,005 | 4,028,161 | 2,350,333 | 53.3 | 57.6 | 0.7 | 0.7 | 99.3 | 2.49 |
| S61 | m | SYL | 28.9 | 11,144,581 | 6,658,021 | 2,667,871 | 59.7 | 57.2 | 0.6 | 0.5 | 99.5 | 2.44 |
| S62 | m | SYL | 28.9 | 10,998,006 | 6,742,826 | 2,506,560 | 61.3 | 57.4 | 0.6 | 0.5 | 99.4 | 2.44 |
| S63 | m | KIE | 18 | 11,152,063 | 6,896,108 | 2,623,103 | 61.8 | 56.2 | 0.6 | 0.5 | 99.5 | 2.50 |
| S64 | m | KIE | 18 | 10,905,488 | 6,409,362 | 2,634,685 | 58.8 | 56.2 | 0.6 | 0.5 | 99.5 | 2.46 |
| S65 | f | NYN | 6 | 12,485,843 | 7,555,803 | 2,926,648 | 60.5 | 53.8 | 0.6 | 0.5 | 99.5 | 2.49 |
| S66 | f | NYN | 6 | 13,455,950 | 8,490,473 | 3,088,221 | 63.1 | 59.1 | 0.6 | 0.5 | 99.5 | 2.39 |
| S73 | f | SYL | 28.9 | 8,901,147 | 5,117,469 | 2,560,420 | 57.5 | 56.6 | 0.8 | 0.7 | 99.4 | 2.50 |
| S74 | f | SYL | 28.9 | 6,650,655 | 3,652,399 | 2,007,162 | 54.9 | 59.5 | 0.8 | 0.7 | 99.5 | 2.32 |
| S77 | m | KIE | 18 | 9,014,946 | 5,232,127 | 2,587,729 | 58.0 | 58.7 | 0.8 | 0.7 | 99.3 | 2.37 |
| S78 | m | KIE | 18 | 9,650,434 | 5,788,601 | 2,744,058 | 60.0 | 58.6 | 0.8 | 0.7 | 99.3 | 2.53 |
| S80 | f | NYN | 6 | 8,622,807 | 5,206,629 | 2,510,145 | 60.4 | 57.0 | 0.8 | 0.7 | 99.4 | 2.50 |
| S85 | m | NYN | 6 | 10,321,779 | 5,874,747 | 2,665,411 | 56.9 | 57.7 | 0.6 | 0.5 | 99.4 | 2.46 |
| S86 | m | NYN | 6 | 8,879,732 | 5,427,464 | 2,286,757 | 61.1 | 58.5 | 0.6 | 0.5 | 99.5 | 2.45 |
| S87 | f | SYL | 28.9 | 10,835,874 | 6,552,344 | 2,484,095 | 60.5 | 55.5 | 0.6 | 0.5 | 99.5 | 2.51 |
| S88 | f | SYL | 28.9 | 11,103,294 | 6,802,303 | 2,640,248 | 61.3 | 58.5 | 0.6 | 0.5 | 99.4 | 2.50 |

| | | | | | | | | | | | | |
|---|---|---|---|---|---|---|---|---|---|---|---|---|
| S89 | f | KIE | 18 | 13,340,852 | 7,663,569 | 2,949,110 | 57.4 | 57.2 | 0.6 | 0.5 | 99.4 | 2.40 |
| S90 | f | KIE | 18 | 12,725,951 | 7,916,399 | 2,971,993 | 62.2 | 55.7 | 0.6 | 0.5 | 99.4 | 2.45 |

**Table S6:** The number of DMS for each of the two pairwise population comparisons (pop-DMS), the overlap with the DMS obtained from the field (KIE) vs. experiment (control group from KIE population) comparison, which were filtered out to remove potential lab artefacts, the resulting number of pop-DMS and their associated number of genes.

| Comparison | # of pop-DMS | overlap with field (KIE) vs. experiment (control group) comparison | # of pop-DMS used in downstream analysis | # of pop-DMS associated to genes | # of genes associated to pop-DMS |
|---|---|---|---|---|---|
| KIE vs. NYN (20 vs. 6 PSU) | 1,990 | 520 | 1,470 | 1,098 | 655 |
| KIE vs. SYL (20 vs. 33 PSU) | 1,663 | 505 | 1,158 | 871 | 510 |

2. CEGX Bioinfomatics Team. Cambridge Epigenetix (CEGX), Babraham Research Campus, Cambridge. 2015.
