## Supplementary material for "Two different epigenetic pathways detected in wild three-spined sticklebacks are involved in salinity adaptation": Summary of Scripts

# Summary of scripts used by Melanie J Heckwolf and Britta S Meyer
# for analysis related to the manuscript
# 'DNA methylation facilitates local adaptation and adaptive transgenerational plasticity'
#
#################################
#1 Preprocessing of Fastq files #
#################################

#1.0 Fastqc on raw reads
for file in *.fastq.gz
do
fastqc ${file} -t 16 -o /01FASTQCresults
done

#1.1 FastqtoSAM
for i in *R1_001.fastq.gz; do
    [[ $i =~ (.*)_(.*)_(.*)_R1_001.fastq.gz ]]
    java -Xmx16G -jar picard.jar FastqToSam \
	FASTQ= $i \
	FASTQ2= ${BASH_REMATCH[1]}_${BASH_REMATCH[2]}_${BASH_REMATCH[3]}_R2_001.fastq.gz \
	OUTPUT= ${BASH_REMATCH[1]}_${BASH_REMATCH[2]}_unmapped.bam \
	READ_GROUP_NAME= ${BASH_REMATCH[1]}_${BASH_REMATCH[2]}.HGJFJBBXX.${BASH_REMATCH[3]} \
	SAMPLE_NAME= ${BASH_REMATCH[2]} \
	LIBRARY_NAME= ${BASH_REMATCH[1]}_${BASH_REMATCH[2]} \
	PLATFORM_UNIT= HGJFJBBXX.${BASH_REMATCH[3]} \
	PLATFORM= illumina
done

#1.2 MarkIlluminaAdapters
for i in *_unmapped.bam; do
  #F12149-L1_S1_unmapped.bam
    [[ $i =~ (.*)_unmapped.bam ]]
    java -Xmx8G -jar -jar ~/software/picard-tools-2.7.1/picard.jar MarkIlluminaAdapters \
  I=$i \
  O=${BASH_REMATCH[1]}_unmapped_xt.bam \
  M=${BASH_REMATCH[1]}_unmapped_xt_metrics.txt

##############################################
#2 Mapping the Fastq files to the reference #
##############################################

#2.0 index Reference genome
bwa index GasAcuV1.fa

#2.1  SamToFastq & bwa mem & MergeBamAlignment
java -Xmx16g -jar picard.jar SamToFastq \
INPUT=!file!_unmapped_xt.bam \
FASTQ=/dev/stdout \
CLIPPING_ATTRIBUTE=XT CLIPPING_ACTION=2 \
INTERLEAVE=true \
| \
bwa mem -M -t 16 -p GasAcuV1.fa /dev/stdin \
| \
java -Xmx16G -jar picard.jar MergeBamAlignment \
ALIGNED_BAM=/dev/stdin \
UNMAPPED_BAM=!file!_unmapped.bam \
OUTPUT=!file!_clean.bam \
CREATE_INDEX=true \
R=GasAcuV1.fa \
MAX_INSERTIONS_OR_DELETIONS=-1 \
ATTRIBUTES_TO_RETAIN=XS \
PRIMARY_ALIGNMENT_STRATEGY=MostDistant \
CLIP_ADAPTERS=false	\
ADD_MATE_CIGAR=true
done

############################################
#3 BAM files - manipulating and statistics #
############################################

#3.1  MarkDuplicates

java -Xmx100G -jar picard.jar MarkDuplicates \
      I=!file!_clean.bam \
      O=!file!_clean_dedup.bam \
      CREATE_INDEX=true \
      METRICS_FILE=!file!_clean_dedup_metrics.txt \
      MAX_FILE_HANDLES_FOR_READ_ENDS_MAP=1000

#3.2 statistics
#3.2.1
echo 'picard'
java -Xmx16G -jar picard.jar CollectMultipleMetrics \
      I=!file!_clean_dedup.bam \
      O=!file!_picard_multiple_metrics \
      R=GasAcuV1.fa \
      PROGRAM=CollectAlignmentSummaryMetrics \
      PROGRAM=QualityScoreDistribution \
      PROGRAM=MeanQualityByCycle \
      PROGRAM=CollectInsertSizeMetrics \
      PROGRAM=CollectBaseDistributionByCycle \
      PROGRAM=CollectGcBiasMetrics

java -Xmx16G -jar picard.jar CollectRawWgsMetrics \
     I=!file!_clean_dedup.bam \
     O=!file!_picard_raw_wgs_metrics.txt \
     R=GasAcuV1.fa \
     INCLUDE_BQ_HISTOGRAM=true

java -Xmx16G -jar picard.jar CollectWgsMetrics \
     I=!file!_clean_dedup.bam \
     O=!file!_picard_wgs_metrics.txt \
     R=GasAcuV1.fa \
     INCLUDE_BQ_HISTOGRAM=true

java -Xmx16G -jar picard.jar CollectWgsMetricsWithNonZeroCoverage \
       I=!file!_clean_dedup.bam  \
       O=!file!_picard_collect_wgs_metrics.txt \
       CHART=!file!_picard_collect_wgs_metrics.pdf  \
      R=GasAcuV1.fa \
#3.2.2
echo 'GATK'
 java -jar GenomeAnalysisTK-3.7.jar \
   -T DepthOfCoverage \
   -R GasAcuV1.fa \
   -o !file!_out \
   -I !file!_clean_dedup.bam


###############################################
#3 GATK - Haplotype calling and SNP filtering #
###############################################
#3.1 Haplotypecaller

java -Xmx96G -jar GenomeAnalysisTK-3.7.jar -T HaplotypeCaller -nct 8 \
      -R GasAcuV1.fa \
      -I !file!_clean_dedup.bam \
      -o !file!_output.raw.snps.indels.g.vcf \
      --emitRefConfidence GVCF


#3.2 joint_genotyping.sh GenotypeGVCFs
#Perform joint genotyping on gVCF files (here in a file with absolute paths all_gvcf.list) produced by HaplotypeCaller

echo GenotypeGVCFs
java -Xmx80G -jar GenomeAnalysisTK-3.7.jar -T GenotypeGVCFs \
      -R GasAcuV1.fa \
      -V all_gvcf.list \
      -o rawVariants_01.vcf \

#3.3 Filter SNPs

#3.3.1# ------- STEP 1: select SNPs -------------------------------
echo getting SNPs
java -Xmx30G -jar GenomeAnalysisTK-3.7.jar -T SelectVariants \
      -R GasAcuV1.fa \
      -V rawVariants_01.vcf \
      -selectType SNP \
      -o raw_snps_01.vcf \


#3.3.2# --------- STEP 2: get statistics --------------------------
java -Xmx30G -jar GenomeAnalysisTK-3.7.jar -T VariantsToTable \
-R GasAcuV1.fa \
-V raw_snps_01.vcf \
-o prep/raw_snps_01.filter.table \
-F ID -F CHROM -F POS -F FILTER -F QD -F FS \
-F SOR -F MQ -F MQRankSum -F ReadPosRankSum -F NO-CALL -F TRANSITION -F HET -F NCALLED \
--allowMissingData \
--showFiltered

#######################################
#3.3.2.1 R scripts to plot statistics #
#######################################
library(dplyr)
library(ggplot2)
data <- read.csv('raw_snps_01.filter.table',sep='\t')

names(data)
#https://software.broadinstitute.org/gatk/guide/article?id=3225

# ------------ plots for normal X axis ---------------------------
ggplot(data, aes(x=QD)) + geom_density(fill=rgb(0,0,.5,.5))+
  geom_vline(xintercept = c(2.5,4))#+
ggsave(paste('plots_SNPs/summary_QD_2.5_4.pdf',sep=''), width = 7,height = 3)

ggplot(data, aes(x=FS)) + geom_density(fill=rgb(0,0,.5,.5))+
  geom_vline(xintercept = c(25,60))#+
ggsave(paste('plots_SNPs/summary_FS_25_60.pdf',sep=''), width = 7,height = 3)

# ------------ plots for log X axis ---------------------------
ggplot(data, aes(x=log(FS,base = 10))) + geom_density(fill=rgb(0,0,.5,.5))+
  geom_vline(xintercept = log(c(25,50,60,75),base = 10))
ggsave('plots_SNPs/summary_log10-FS_25-50-60-75.pdf', width = 7,height = 3)

ggplot(data, aes(x=SOR)) + geom_density(fill=rgb(0,0,.5,.5))+
  geom_vline(xintercept = c(2))#+
ggsave(paste('plots_SNPs/summary_SOR_2.pdf',sep=''), width = 7,height = 3)

ggplot(data, aes(x=MQ)) + geom_density(fill=rgb(0,0,.5,.5))+
  geom_vline(xintercept = c(40,59.5))+
scale_x_continuous(limits = c(35,70))
ggsave(paste('plots_SNPs/summary_MQ_40_59.5.pdf',sep=''), width = 7,height = 3)


ggplot(data, aes(x=MQRankSum)) + geom_density(fill=rgb(0,0,.5,.5))+
  geom_vline(xintercept = c(-1.5))+
  scale_x_continuous(limits = c(-3,3))
ggsave(paste('plots_SNPs/summary_MQRankSum_-1.5.pdf',sep=''), width = 7,height = 3)

ggplot(data, aes(x=ReadPosRankSum)) + geom_density(fill=rgb(0,0,.5,.5))+
  geom_vline(xintercept = c(-2))+
  scale_x_continuous(limits = c(-15,15))
ggsave(paste('plots_SNPs/summary_ReadPosRankSum_-2.pdf',sep=''), width = 7,height = 3)


#3.3.3# --------- STEP 3: filter ---------------------------------
echo VariantFiltration
java -Xmx30G -jar GenomeAnalysisTK-3.7.jar \
   -T VariantFiltration \
   -R GasAcuV1.fa \
   -V raw_snps_01.vcf \
   --filterExpression "QD < 4.0" \
   --filterName "low_QD" \
   --filterExpression "FS > 25.0" \
   --filterName "high_FS" \
   --filterExpression "SOR > 2.0" \
   --filterName "high_SOR" \
   --filterExpression "MQ < 40.0" \
   --filterName "low_MQ" \
   --filterExpression "MQRankSum < -1.5" \
   --filterName "low_MQRankSum" \
   --filterExpression "ReadPosRankSum < -2.0" \
   --filterName "low_ReadPosRankSum" \
   -o filtered_snps.vcf

#3.3.4# ------ STEP 4: kick out unfiltered variants -------------
echo SelectVariants
java -Xmx20G -jar GenomeAnalysisTK-3.7.jar \
    -T SelectVariants \
    -R GasAcuV1.fa \
    -V filtered_snps.vcf \
    -o clean_snps.vcf \
    --excludeFiltered


####################################
#4.0 Further Filtering of the SNPs #
####################################

#4.1
#filter for quality --> genotype quality
vcftools --gzvcf clean_snps.vcf.gz --minGQ 20 --recode --recode-INFO-all --out clean_snps_gq20

#4.2
#filter for MAF 0.05%
vcftools --vcf clean_snps_gq20.recode.vcf  --maf 0.0049 --recode --recode-INFO-all --out clean_snps_gq20_maf_0.005

#4.3
#for further population genetic analyses (such as FSt and demography) we would filter for missing data, e.g. 20% missing allowed and biallelic states


##########################################
#5. Exclude SNPs for methylation analyses#
##########################################

bcftools view -H clean_snps_gq20_maf_0.005.recode.vcf -O v -o Noheader-clean_snps_gq20_maf_0.005.recode.vcf
perl snip_detect_CT.pl Noheader-clean_snps_gq20_maf_0.005.recode.vcf
perl snip_detect_GA.pl Noheader-clean_snps_gq20_maf_0.005.recode.vcf

#########################
#5.1 snip_detect_CT.pl. #
#########################
# perl script for CT

#!/usr/bin/perl
#print "base col A: ";
$pat_A=C;

#print "base col B: ";
$pat_B=T;

$col_A = 3;
$col_B = 4;

open  (FILEIN, '<', @ARGV[0]) || die  "Error: fnf.";
open  (FILEOUT,'>', 'result_pat_'.C.'_'.T.'.txt');


while(<FILEIN>){

  chomp($_);
  @eintraege = split(/\t/, $_);

  $hit_A = @eintraege[$col_A] =~ m/C/;
  $hit_B = @eintraege[$col_B] =~ m/T/;

  if($hit_A == 1 && $hit_B == 1){
    print FILEOUT @eintraege[0]."\t".@eintraege[1]."\n";
    }
}

close FILEIN;
close FILEOUT;

exit 0;

# perl script for GA

##########################
#5.2 snip_detect_GA.pl. #
##########################

#!/usr/bin/perl

$col_A = 3;
$col_B = 4;

open  (FILEIN, '<', @ARGV[0]) || die  "Error: fnf.";
open  (FILEOUT,'>', 'result_pat_'.G.'_'.A.'.txt');


while(<FILEIN>){

  chomp($_);
  @eintraege = split(/\t/, $_);

  $hit_A = @eintraege[$col_A] =~ m/G/;
  $hit_B = @eintraege[$col_B] =~ m/A/;

  if($hit_A == 1 && $hit_B == 1){
    print FILEOUT @eintraege[0]."\t".@eintraege[1]."\n";
    }
}

close FILEIN;
close FILEOUT;

exit 0;

##########################################################################
#6. R script using Methykit and GenomicRanges to filter methylation calls#
##########################################################################
#
# read methylation calls resulting from Bismark pipeline into Methylkit
myobjDB= methRead(file_path,
                    sample_id,
                    assembly = "stickleback_GasAcu1",
                    mincov = 1,
                    pipeline = "bismarkCoverage",
                    context = "CpG",
                    treatment_id,
                    dbtype = "tabix",
                    dbdir = "methylDB4")

# filter dataset for minimum coverage = 10 and remove the 99.9th percentile
filtered.myobj=filterByCoverage(myobjDB,lo.count=10,lo.perc=NULL,
                                 hi.count=NULL,hi.perc=99.9, suffix="filtered_min10x")

# normalize coverage values between samples
norm_filtered.myobj = normalizeCoverage(filtered.myobj,method="median")

# excluded all sites that were present in fewer than 9 individuals per treatment group
unite_norm_10x_9L=unite(norm_filtered.myobj, destrand=FALSE,min.per.group=9L,mc.cores=8)

# C-to-T and G-to-A-SNPs were derived from custom written perl scripts (see 5)
# from the 96 wild caught individuals
library("GenomicRanges")

#' BED to GRanges
#'
#' This function loads a BED-like file and stores it as a GRanges object.
#' The tab-delimited file must be ordered as 'chr', 'start', 'end', 'id', 'score', 'strand'.
#' The minimal BED file must have the 'chr', 'start', 'end' columns.
#' Any columns after the strand column are ignored.
#'
#' @param file Location of your file
#' @keywords BED GRanges
#' @export
#' @examples
#' bed_to_granges('my_bed_file.bed')

bed_to_granges <- function(file){
  df <- read.table(file,
                   header=F,
                   stringsAsFactors=F)

  if(length(df) > 6){
    df <- df[,-c(7:length(df))]
  }

  if(length(df)<3){
    stop("File has less than 3 columns")
  }

  header <- c('chr','start','end','id','score','strand')
  names(df) <- header[1:length(names(df))]

  if('strand' %in% colnames(df)){
    df$strand <- gsub(pattern="[^+-]+", replacement = '*', x = df$strand)
  }

  if(length(df)==3){
    gr <- with(df, GRanges(chr, IRanges(start, end)))
  } else if (length(df)==4){
    gr <- with(df, GRanges(chr, IRanges(start, end), id=id))
  } else if (length(df)==5){
    gr <- with(df, GRanges(chr, IRanges(start, end), id=id, score=score))
  } else if (length(df)==6){
    gr <- with(df, GRanges(chr, IRanges(start, end), id=id, score=score, strand=strand))
  }
  return(gr)
}


# load CT SNPs and duplicate column and save as bedfile, result_pat_C_T_gq20maf.bed is created in step 5.1
CT_gq20maf= read.csv(file="result_pat_C_T_gq20maf.bed", sep="\t", header=FALSE)
CT_gq20maf = cbind(CT_gq20maf,V3=rep(CT_gq20maf$V2))
write.table(CT_gq20maf,file="result_pat_C_T_gq20mafSTARTEND.bed",row.names = FALSE,quote = FALSE,sep="\t", col.names = FALSE)

blacklist_CT_bed<- bed_to_granges(file="result_pat_C_T_gq20mafSTARTEND.bed")

# load GA SNPs and duplicate column and save as bedfile, result_pat_G_A_gq20maf.bed is created in step 5.2
AG_gq20maf= read.csv(file="result_pat_G_A_gq20maf.bed", sep="\t", header=FALSE)
AG_gq20maf = cbind(AG_gq20maf,V3=rep(AG_gq20maf$V2))
write.table(AG_gq20maf,file="result_pat_G_A_gq20mafSTARTEND.bed",row.names = FALSE,quote = FALSE,sep="\t", col.names = FALSE)

blacklist_GA_bed<- bed_to_granges(file="result_pat_G_A_gq20mafSTARTEND.bed")


####  remove ChrM
blacklist_CT_bed_without_ChrM<- blacklist_CT_bed[!seqnames(blacklist_CT_bed) == 'chrM']
blacklist_GA_bed_without_ChrM<- blacklist_GA_bed[!seqnames(blacklist_GA_bed) == 'chrM']


# interesect bedfile and unite-file
#### create overlap --> these positions are corrected for CT SNPs
unite_norm_10x_9LGRanges <- as(unite_norm_10x_9L,"GRanges")
Overlap_CT=unite_norm_10x_9LGRanges[countOverlaps(unite_norm_10x_9LGRanges, blacklist_CT_bed_without_ChrM) == 0L]
unite_norm_10x_9L_CT <- selectByOverlap(unite_norm_10x_9L,Overlap_CT)

# make methylkit database
objDB_CT=makeMethylDB(unite_norm_10x_9L_CT,"methylBaseDB")

##### create overlap --> these positions are corrected for GA SNPs
unite_norm_10x_9L_CT_GRanges <- as(unite_norm_10x_9L_CT,"GRanges")
Overlap_GA=unite_norm_10x_9L_CT_GRanges[countOverlaps(unite_norm_10x_9L_CT_GRanges, blacklist_GA_bed_without_ChrM) == 0L]
unite_norm_10x_9L_CT_GA <- selectByOverlap(objDB_CT,Overlap_GA)


#remove sex chr.
unite_norm_10x_9L_CT_GA_Granges <- as(unite_norm_10x_9L_CT_GA,"GRanges")
unite_norm_10x_9L_CT_GA_Granges_nosex <- unite_norm_10x_9L_CT_GA_Granges[!seqnames(unite_norm_10x_9L_CT_GA_Granges) == 'chrXIX']

unite_norm_10x_9L_CT_GA_DB=makeMethylDB(unite_norm_10x_9L_CT_GA,"methylBaseDB")
unite_norm_10x_9L_CT_GA_nosex_DB <- selectByOverlap(unite_norm_10x_9L_CT_GA_DB,unite_norm_10x_9L_CT_GA_Granges_nosex)


##########################################################################
#7. R script using Methykit to identify DMS in pairwise comparisons ######
##########################################################################

# all pairwise comparisons:
# wild populations: KIE (20 PSU) vs NYN (6 PSU)
# wild populations: KIE (20 PSU) vs SYL (33 PSU)
# experiment: control vs within-generational 6 PSU
# experiment: control vs transgenerational 6 PSU
# experiment: control vs within-generational 33 PSU
# experiment: control vs transgenerational 6 PSU

# one example:
table_subset_0vs1=subset(treatment_info_table,treat_number==0 | treat_number==1)
unite_subset_0vs1=reorganize(unite_norm_10x_9L_CT_GA_nosex_DB,
                             sample.ids=as.character(table_subset_0vs1[,2]),
                             treatment=table_subset_0vs1[,3])
#METHDIFF - treatment (group: 1) - control (group: 0)
myDiff_0vs1=calculateDiffMeth(unite_subset_0vs1,mc.cores = 8)
myDiff_0vs1_significant=getMethylDiff(myDiff_0vs1,difference=15,qvalue=0.0125)


# Annotate DMS with gene prediction file (promoter, exon intron, intergenic)
# promoter definition is 1500 up and 500 downstream of TSS (transcription start site)

#ensGenegenepred.bed is a bed-file created from the UCSC-Genome table browser http://genome.ucsc.edu/index.html
#using the "ensGene" gene and gene prediction for sticklebacks (Broad/gasAcu1) in the so-called pred-format (excluding the bin column),
#which was converted with "genepredtobed ensgene.pred ensGenegenepred.bed
#and subsequently sorted  "bedtools sort -i  ensGenegenepred.bed > ensGenegenepred.bed"

gene.obj=readTranscriptFeatures("ensGenegenepred.bed",
                                up.flank = 1500, down.flank = 500, unique.prom = T)
anno_0vs1=annotateWithGeneParts(as(myDiff_0vs1_significant,"GRanges"),gene.obj)

# get association of DMS and TSS (transcription start site)
assTSS_0vs1=getAssociationWithTSS(anno_0vs1)

data_0vs1=cbind(getData(myDiff_0vs1_significant),as.data.frame(assTSS_0vs1),anno_0vs1@members)


######################################################################################################
#8. vcftools   #
######################################################################################################
#!/bin/bash

#1 comparison KIE_SYL is used to generate vcfs allowing for 40 % (= 13 IND) of missing data and a read depth of >=5
vcftools --vcf comp.KIE_SYL.gq20_maf0.05.mac1.recode.vcf.recode.vcf  \
--max-missing-count 13  \
--max-alleles 2 \
--recode \
--minDP 5 \
--out comp.KIE_SYL.gq20_maf0.05.mac1.biallelic.maxmiss13ind_DP5

###create a vcf file subset to the inducible DMS (148sites) in increasing PSU (_33PSU) with 5kb before and after DMS (_plus.minus.5kb)
vcftools --vcf comp.KIE_SYL.gq20_maf0.05.mac1.biallelic.maxmiss13ind_DP5.recode.vcf  \
  --out DP5_13miss_genes_KIE_SYL_148_5kb  \
  --recode  \
  --bed DMS_inducible_148sites_33PSU_plus.minus.5kb.txt.bed
### DMS_inducible_148sites_33PSU_plus.minus.5kb.txt.bed is a file containing the location of the 10kb window around each of the 148 inducible pop-DMS between KIE and SYL


##run fst
vcftools --vcf DP5_13miss_genes_KIE_SYL_148_5kb.recode.vcf \
  --weir-fst-pop popKIE.txt \
  --weir-fst-pop popSYL.txt \
  --keep popKIE.txt \
  --keep popSYL.txt \
  --out DP5_13miss_KIE_vs_SYL_148_5kb

#2 comparison KIE_NYN is used to generate vcfs allowing for 40 % (= 13 IND) of missing data and a read depth of >=5

vcftools --vcf comp.KIE_NYN.gq20_maf0.05.mac1.recode.vcf.recode.vcf  \
--max-missing-count 13  \
--max-alleles 2 \
--minDP 5 \
--recode \
--out comp.KIE_NYN.gq20_maf0.05.mac1.biallelic.maxmiss13ind_DP5

###create a vcf file subset to the inducible DMS (198sites) in decreasing PSU (_6PSU) with 5kb before and after DMS (_plus.minus.5kb)
vcftools --vcf comp.KIE_NYN.gq20_maf0.05.mac1.biallelic.maxmiss13ind_DP5.recode.vcf  \
  --out DP5_13miss_genes_KIE_NYN_198_5kb  \
  --recode  \
  --bed DMS_inducible_198sites_6PSU_plus.minus.5kb.txt.bed
## DMS_inducible_198sites_6PSU_plus.minus.5kb.txt.bed is a file containing the location of the 10kb window around each of the 198 inducible pop-DMS between KIE and NYN

##run fst
vcftools --vcf DP5_13miss_genes_KIE_NYN_198_5kb.recode.vcf \
  --weir-fst-pop popNYN.txt \
  --weir-fst-pop popKIE.txt \
  --keep popNYN.txt \
  --keep popKIE.txt \
  --out DP5_13miss_NYN_vs_KIE_198_5kb


###############################################
#9. R script: randomization test on Fst values#
###############################################

#randomization tests (10,000x bootstrap)
#
# script was run for 33 and 6 PSU seperately

# table_FST is a table with mean FST values per CpG site calculated in a 10kb window (+/- 5kb from CpG)
# direction = drection of inducible sites (opposite or expected)
# take mean FST over all expected and all inducible sites:
gd <- table_FST %>% group_by(direction) %>% summarize(mean_Fst=mean(mean_Fst))

# this is the difference in mean FST for the original data:
table_randomization=data.frame(group="original",difference=gd[1,2]-gd[2,2])

# bootstrap randomization test:
# calculate 10,000 differences in mean FST for randomly assigned dataset (mean FST
# is randomly assigned to opposite or expected, with replacement as it is a bootstrap randomization test)
for (i in 1:10000){
  expected=mean(sample(table_FST$mean_Fst, 101, replace = T))
  opposite=mean(sample(table_FST$mean_Fst, 47, replace = T))
  table_randomization=rbind(table_randomization,data.frame(group="random",mean_Fst=expected-opposite))
}

table_randomization_pval=subset(table_randomization,abs(table_randomization$mean_Fst) < (as.numeric(gd[1,2]-gd[2,2])))
# P-value results from:
pvalue=length(table_randomization_pval$mean_Fst)/10000
